# Transfer Matrix and Tensor Approaches to Nucleic Acid Secondary Structure Thermodynamics^*^

**DOI:** 10.64898/2026.09.30.755805

**Authors:** Mark E. Fornace, Niles A. Pierce

## Abstract

Nucleic acid secondary structure thermodynamic models have great utility for the analysis and design of nucleic acid structures, devices, and systems, yet their complexity and piecemeal development over decades pose challenges to further innovation. Employing different functional forms for each loop type within a structure, existing nearest neighbor (NN) models necessitate numerous loop-specific dynamic programming recursions in order to calculate physical quantities of interest. Here, we derive and implement a new class of models based on transfer matrices, yielding a single functional form for the free energy of all loop types (as the trace of a matrix product) and a single dynamic programming recursion for calculating physical quantities of interest. This new class of models has additional attractive properties: 1) incorporating a subensemble of states describing proximal interactions within each loop, 2) approximating logarithmic entropic scaling for all closed loops, 3) offering the opportunity to model correlations between the subensembles of neighboring loops by generalizing transfer matrices to tensors, and 4) providing a basis for systematically adjusting the number of model parameters to enhance accuracy as new empirical data become available for parameter regression. We use literature experimental data to regress parameters for representative new transfer matrix models, observing accuracy as good as or better than that of existing NN models, yet using dramatically fewer parameters (e.g., dozens vs hundreds or thousands). For complex ensembles containing an arbitrary number of interacting nucleic acid strands totaling *N* nucleotides, the new single recursion leads to exact dynamic programs for calculating diverse physical quantities (including the partition function, equilibrium base-pairing probabilities, minimum free energy and suboptimal proxy structures, Boltzmann-sampled secondary structures, and gradient of the partition function) with *O*(*N*^3^) time complexity. The simplicity of the new partition function dynamic program enables overflow-safe partition function calculations using renormalization to achieve scalability for large complexes (e.g., 50,000 nt) as well as formulation using a global matrix multiplication primitive that facilitates GPU acceleration, yielding speedups of an order of magnitude relative to NN partition function algorithms for large complexes. The new class of transfer matrix and tensor models provides an elegant framework for future efforts to advance the accuracy, speed, and scope of computational nucleic acid analysis and design.

**Relevance to Life Sciences:** The programmable chemistry of nucleic acid base-pairing is central to the biological circuits that orchestrate life and to the emerging fields of molecular programming, nucleic acid nanotechnology, and synthetic biology.

**Mathematical Content:** This paper introduces the use of transfer matrices and tensor contraction to model nucleic acid secondary structure thermodynamics, providing an extensible framework for enhancing model fidelity as new experimental training data become available, and leading to dramatically simplified dynamic programming recursions that can be formulated linear algebraically for GPU acceleration.

## 1. Introduction

Life is orchestrated by programmable biomolecules (DNA, RNA, and proteins) interacting within complex machines and circuits. These biological proofs-of-principle motivate diverse engineering efforts within the emerging disciplines of molecular programming, nucleic acid nanotechnology, and synthetic biology. In this context, the programmable chemistry of nucleic acid base pairing offers researchers a rich design space *in vitro, in situ*, and *in vivo*, enabling the rational design of complex structures and dynamic functions including sensing, amplification, signal transduction, logic, locomotion, and regulation. ^4,34,35^ These engineering efforts are greatly assisted by the use of computational algorithms to design and analyze molecular sequences before experimental testing commences in the wet lab. Notably, nucleic acid energetics can be usefully approximated using empirical models based only on secondary structure (i.e., the base-pairing state of a set of nucleic acid strands). ^11,13–15,24,25,27,29,31^ As a result, nucleic acid secondary structure models have great utility for analyzing and designing functional nucleic acid systems, and they offer the significant benefit that analysis and design problems can be rigorously formulated over well-defined structural ensembles. In this work, we revisit the secondary structure ensembles, free energy models, and dynamic programming algorithms that have paced progress in nucleic acid engineering for decades, retaining strengths and addressing shortcomings to arrive at elegant new formulations that enhance performance and offer an extensible framework for further innovation.

### 1.1. Existing Secondary Structure Complex Ensemble

The *sequence, ϕ*, of one or more interacting RNA strands totaling *N* nucleotides is specified as a 5′ to 3′ list of nucleotides *ϕ*_*i*_ ∈ {A,C,G,U} for *i* = 1, …, *N*. For DNA, *ϕ*_*i*_ ∈ {A,C,G,T}. A *secondary structure, s*, of one or more interacting RNA strands is defined by a set of base pairs, each a Watson–Crick pair (A·U and C·G) or a wobble pair (G·U). For DNA, Watson–Crick pairs are A·T and C·G and there are no wobble pairs. ^25^ Figure 1a depicts an example secondary structure for a complex of three strands. For algorithmic purposes, it is convenient to describe secondary structures using a *polymer graph* representation, constructed by ordering the strands around a circle, drawing the backbones in succession from 5′ to 3′ around the circumference with a *nick* between each strand, and drawing straight lines connecting paired bases (e.g., Figure 1b). A secondary structure is *unpseudoknotted* if there exists a strand ordering for which the polymer graph has no crossing lines (e.g., Figure 1b), or *pseudoknotted* if all strand orderings contain crossing lines. A secondary structure is *connected* if no subset of the strands is free of (i.e., unpaired to) the others. ^6^

**Figure 1.**
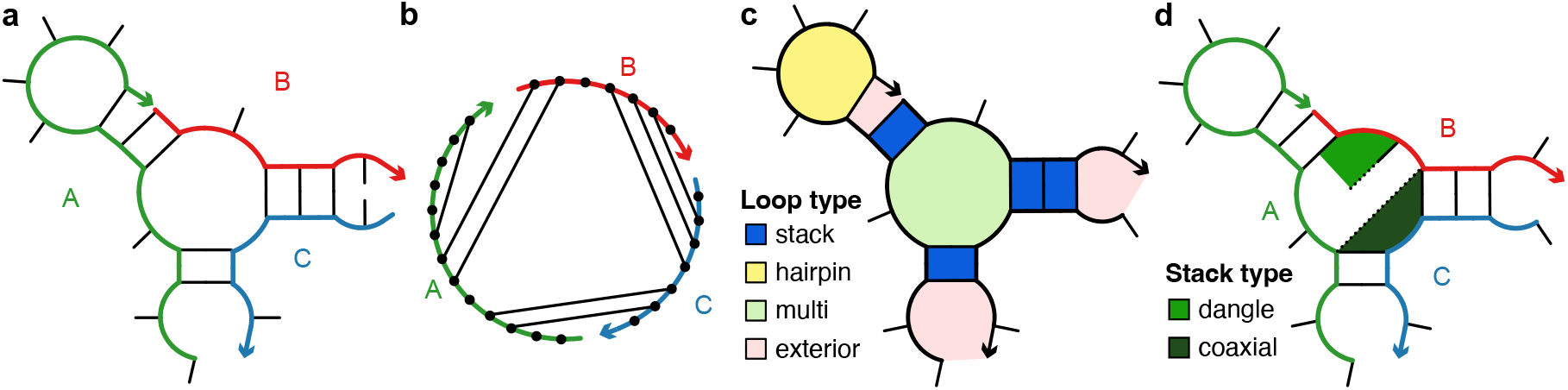
Nucleic acid secondary structure. **(a)** Connected unpseudoknotted secondary structure for a complex of 3 strands with strander ordering *π* = ABC. An arrowhead denotes the 3′ end of each strand. **(b)** Polymer graph representation of the same secondary structure. **(c)** Example loop types. **(d)** Example coaxial and dangle stacks.

Consider a complex of *L* distinct strands (each with a unique identifier in {1, …, *L*}) corresponding to strand ordering *π*. The *complex ensemble* 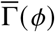 contains all connected polymer graphs with no crossing lines (i.e., all unpseudoknotted secondary structures). ^6^ As a matter of algorithmic necessity, dynamic programs used to calculate physical quantities of interest operate on complex ensemble 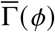 treating all strands as distinct. However, in the wet lab, strands with the same sequence are typically indistinguishable with respect to experimental observables. For comparison to experimental data, physical quantities calculated over ensemble 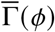 are post-processed to obtain the corresponding quantities calculated over *complex ensemble* Γ(*ϕ*) in which strands of the same species are treated as indistinguishable. ^6^ Interestingly, ensembles 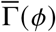 and Γ(*ϕ*) both have utility when examining the physical properties of a complex as they provide related but different perspectives, akin to complementary thought experiments. ^7^

Each secondary structure 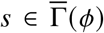 can be decomposed into a set of *loops*, each bounded by one or more closing base pairs (see the example loops shaded in Figure 1c). Within a loop, a closing base pair can form a *coaxial stack* with an adjacent closing base pair or a *dangle stack* with an adjacent unpaired base (see the example coaxial stack and dangle stack of Figure 1d). Hence, each secondary structure within the complex ensemble can be associated with a subensemble of stacking states compatible with that secondary structure, ^7^ providing a basis for physical modeling of loop free energies that goes beyond base-pairing information alone. With existing secondary structure models, however, the subensemble of stacking states is modeled only for certain loop types, and the subensemble concept does not encompass interactions between adjacent unpaired bases, leading to the oversimplification that all unstructured strands are modeled identically regardless of sequence.

### 1.2. Existing Nearest-Neighbor (NN) Free Energy Models

For sequence *ϕ* in secondary structure 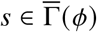, the free energy,

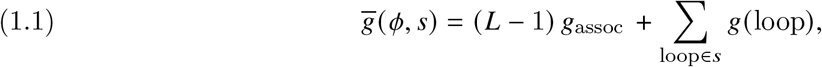

is estimated using existing models as a sum of loop free energies ^13,15,24,30,32,36^ plus a strand association penalty, ^3^ *g*_assoc_, applied *L* − 1 times for a complex of *L* strands. ^6^ To simplify the equations that follow, the free energy 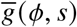 is dimensionless, and the standard dimensional free energy is simply 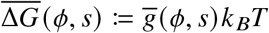, where *k* _*B*_ is the Boltzmann constant and *T* is temperature. Loop free energies, *g* (loop), taking into consideration *nearest-neighbor* (NN) information between nucleotides and base pairs, have been regressed empirically over decades by numerous research groups. ^13,15,24,30,32,36^ Different sequence- and/or size-dependent functional forms are used depending on loop type and size: *hairpin loops* closed by a single base pair, *interior loops* closed by two base pairs (*bulge loops* and *stack loops* are treated as special cases of interior loops), *multiloops* closed by three or more base pairs, and *exterior loops* containing a nick between strands and any number of base pairs (see the example loop types of Figure 1c). Many pages are required to describe all the functional forms and edge cases for the different loop types (see Section S1.7 of Reference 7). The piecemeal nature of the free energy model significantly complicates the resulting computational algorithms, necessitating loop-specific dynamic programming recursions. Due to varying levels of attention in the experimental literature, certain parameters have been regressed with significantly higher fidelity (e.g., stack loops) than others (e.g., multiloops, large hairpins, large interior loops). For reasons of computational expediency, the functional form for multiloops does not preserve logarithmic scaling for the entropic cost of loop closure with loop size, and the functional form for interior loops is highly idiosyncratic. Existing NN free energy models explicitly incorporate a Boltzmann-weighted subensemble of coaxial and dangle stacking states only within multiloops and exterior loops. ^7^ Furthermore, as new high-throughput empirical data become available for regression of enhanced parameter sets, historical NN functional forms provide no obvious knobs that can be turned to achieve higher model fidelity.

### 1.3. Existing dynamic programming algorithms

Existing dynamic programs over complex ensembles employ a unified dynamic programming framework that enables calculation of diverse physical quantities for interacting nucleic acid strands using three ingredients (see Figure 2a):^7^ 1) *recursions* that specify the dependencies between subproblems and incorporate the details of the structural ensemble and the free energy model, 2) *evaluation algebras* that define the mathematical form of each subproblem, 3) *operation orders* that specify the computational trajectory through the dependency graph of subproblems. The recursions are coded generically and then compiled with a quantity-specific evaluation algebra and operation order to generate an executable for calculating a given physical quantity (e.g., the partition function, equilibrium base-pairing probabilities, minimum free energy (MFE) proxy structures, and Boltzmann-sampled secondary structures). ^7^

**Figure 2.**
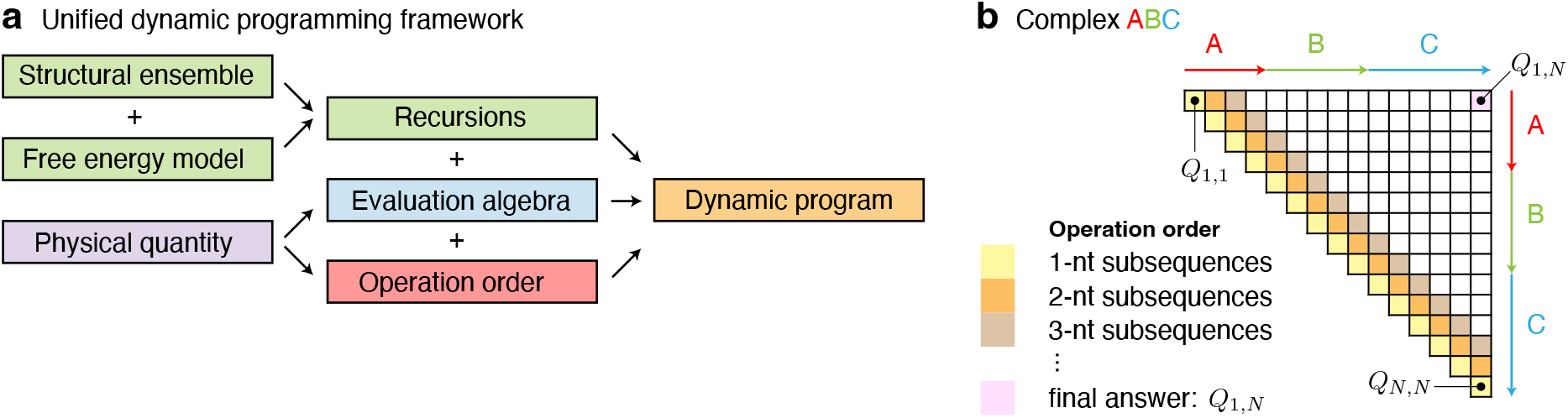
Existing dynamic programming algorithms for calculating physical quantities over the ensemble of a complex. ^7^ **(a)** Unified dynamic programming framework. **(b)** Operation order for partition function dynamic program over complex ensemble *π* = ABC containing *N* nucleotides. Adapted with permission from Reference ^7^. Copyright 2020 American Chemical Society.

For simplicity of exposition, consider calculation of the partition function as a basis for generalization to other physical quantities of interest. For a complex with sequence *ϕ*, the partition function,

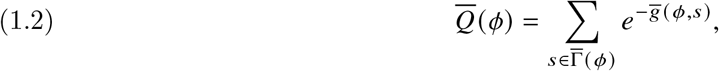

over ensemble 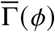 can be used to calculate the equilibrium probability,

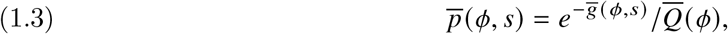

of any secondary structure 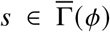. The complex ensemble size, 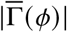, grows exponentially with the number of nucleotides, *N* := |*ϕ*|, but the partition function can be calculated in *O*(*N*^3^) time and *O*(*N*^2^) space using a dynamic programming algorithm. ^6,7,16^ The algorithm calculates the subsequence partition function *Q*_*i, j*_ for each subsequence [*i, j*] via a forward sweep from short subsequences to the full sequence, finally yielding the partition function of the full sequence, *Q*_1,*N*_ (see the operation order of Figure 2b). The recursion used to define subsequence partition function *Q*_*i, j*_ relies on additional restricted partition functions 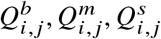, … that are also calculated recursively. To treat the different functional forms of the free energy model for all loops types and sizes with efficient vectorizable algorithms, over 20 different loop-specific recursions are used to incorporate the partition function of each secondary structure, incorporating Boltzmann-weighted contributions for the subensemble of coaxial and dangle stacking states within multiloops and exterior loops. Dozens of pages are required to describe these intricate recursions (see Section S2 of Reference 7). In the context of the *empirical* loop-based free energy model, the resulting polynomial-time algorithm calculates the *exact* partition function over the ensemble of the complex.

### 1.4. A New Approach

To address important shortcomings of existing methods, we propose fundamental changes to the structural ensemble, the free energy model, and the dynamic programming recursions (i.e., we fundamentally change all of the green boxes in Figure 2a) to enhance simplicity, accuracy, and speed, and to facilitate innovation. First, we update the structural ensemble by generalizing the subensemble concept to apply to all nucleotides within a secondary structure regardless of loop type, and propose a new class of free energy models that exploits transfer matrices and/or tensors to employ a single functional form (trace of a matrix product) for all loop types (section 2).

Second, we derive a single dynamic programming recursion for performing exact polynomial-time calculations over the new structural ensemble using the new free energy model (section 3). Third, we use literature empirical data to regress candidate parameter sets for a range of transfer matrix and tensor models and assess the accuracy and speed of the new algorithms relative to existing NN methods (section 4).

## 2. Model formulation

We seek a new class of free energy models that is simpler, more accurate, and more extensible than historical NN secondary structure models. From existing secondary structure models, we retain the complex ensemble, 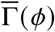, representing all unpseudoknotted connected secondary structures. However, we generalize the subensemble concept, which currently applies only to coaxial and dangle stacking states within multiloops and exterior loops, defining subensembles for all loop types over a set of local interactions more general than stacking states. As a starting point, we retain the concept of calculating the secondary structure free energy, 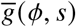, as a sum of constituent loop free energies (1.1). However, to replace numerous loop-specific functional forms, we derive a single functional form for the loop free energy, *g*(loop), that applies to all loop types and sizes. Motivated by the underlying physics of nucleic acids and prior developments in the literature, we use a subensemble beneath secondary structure to enhance fidelity by modeling the physical interactions between neighboring nucleotides. We begin by observing the similarity of a secondary structure loop to a 1D spin system modeled using transfer matrices (subsection 2.1), and then describe a new loop-based structural subensemble and free energy model expressed in terms of matrix product traces (subsection 2.2). We then show that the transfer matrix model can be generalized to a tensor model to allow modeling of correlations between neighboring loops (subsection 2.3), in which case the structure free energy, 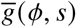, is no longer separable into independent loop free energies. Finally, we derive constraints on the formulation of transfer matrix and tensor models to ensure that they respect logarithmic entropic scaling in closed loops and automatic exclusion of disconnected structures in open loops (subsection 2.6).

### 2.1. Motivation from transfer matrix models

As a useful analogy, consider the classical 1D Ising model (see Figures 3ab), where the free energy of the system is determined solely by spin interactions between neighboring sites. ^10^ The spin, *σ*_*i*_, of site *i* is either up or down, *σ*_*i*_ ∈ {+1, −1}. Assuming a site-specific interaction strength, *J*_*i,i*+1_, between spins *σ*_*i*_ and *σ*_*i*+1_ and no magnetic field, the Hamiltonian describing the energy of a 1D system with *n* sites is 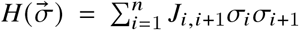 for a periodic system (Figure 3a) satisfying the periodic boundary condition 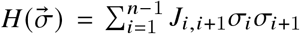 for a system with free ends (Figure 3b). In either case, the free energy over the canonical ensemble may be expressed as:

**Figure 3.**
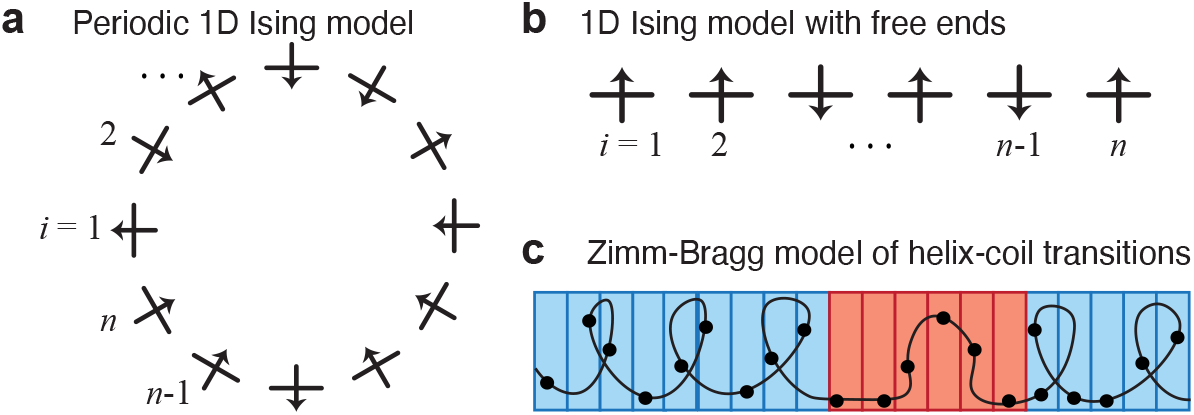
Physical models providing conceptual motivation for a transfer matrix approach to nucleic acid secondary structure thermodynamics. **(a)** Periodic 1D Ising model, analogous to a periodic nucleic acid secondary structure loop containing no nick (e.g., a hairpin loop, an interior loop, or a multiloop in Figure 1c). **(b)** A 1D Ising model with free ends, analogous to a nucleic acid secondary structure loop containing a nick (e.g., an exterior loop in Figure 1c). **(c)** Zimm-Bragg model of helix-coil transitions within biopolymers.

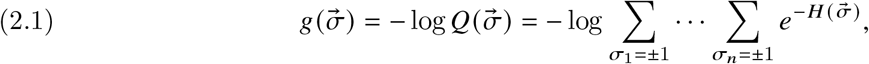

where 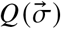 is the partition function representing a sum over 2^*N*^ system states and 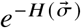 is the Boltzmann factor for system state 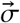.

For a periodic 1D Ising system this becomes

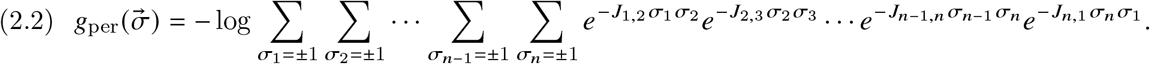

For *i* = 2, …, *n*, the individual sum, 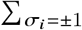, over the two spin states, *σ*_*i*_ = ±1, may be interpreted as a matrix product, *T*_*i*−1/2_*T*_*i*+1/2_, of 2×2 *transfer matrices* with:

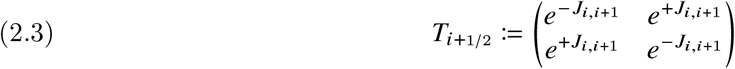

yielding

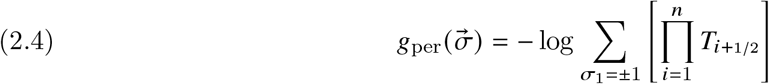

As a result of the periodic boundary condition, summation over the two spin states, *σ*_1_ = ±1, then yields the trace of a matrix product:

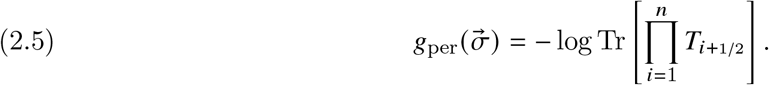

Note that each matrix multiplication, *T*_*i*−1/2_*T*_*i*+1/2_, *contracts* an intervening index *i*, effectively yielding a new transfer matrix between sites *i* − 1 and *i* + 1. Evaluation of the product in (2.5) thus yields an effective transfer matrix between 1 and *n* + 1 and the trace sums only the diagonal entries (for which either *σ*_1_ =*σ*_*n*+1_ =+1 or *σ*_1_ =*σ*_*n*+1_ =−1) because the off-diagonal entries (for which *σ*_*n*+1_ ≠ *σ*_1_) do not satisfy the periodic boundary condition and hence do not contribute to the free energy. The trace of a matrix product is invariant to cyclic permutation (e.g., Tr[*ABC*] = Tr[*BC A*]), so the functional form respects the periodicity of the physical system.

For a 1D Ising system with free ends, (2.1) becomes

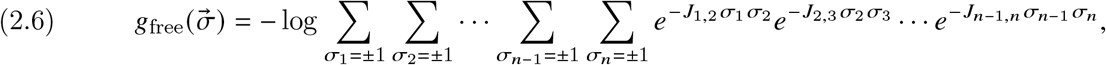

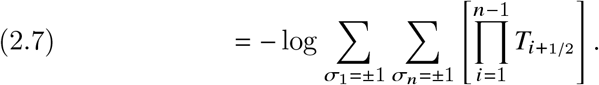

Summation over the four combinations of spin states at the free ends then yields the scalar free energy

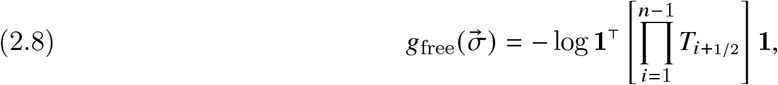

where **1** denotes a column vector of ones. Note that in the case of free ends, evaluation of the product in (2.8) contracts all middle indices to yield an effective transfer matrix between sites 1 and *n*; the vectors of ones sum both diagonal and off-diagonal entries because all four combinations of spin states (*σ*_1_ = ±1 with *σ*_*n*_ = ±1) contribute to the free energy. The Zimm-Bragg model uses an analogous transfer matrix approach to model helix-coil transitions in biopolymers (Figure 3c). These 1D Ising models illustrate that for a physical system with energetics dependent on local interactions between neighboring sites, the free energy of the system can be naturally expressed using transfer matrices, whether the system is periodic (2.5) or has free ends (2.8).

### 2.2. A transfer matrix free energy model with independent loop free energies

We now adapt this transfer matrix concept to estimate the free energy of a loop within a nucleic acid secondary structure. For comparison to the 1D Ising models of Figure 3ab, we number the nucleotides within a loop from *i* = 1, …, *n* (Figure 4ab). We introduce a transfer matrix 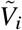 for each unpaired nucleotide *i* and a transfer matrix 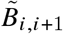 for each base pair *i* · *i* + 1. To facilitate matrix products, we define 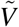 and 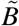 to be square *γ* × *γ* matrices. Analogous to (2.3), each matrix entry may be interpreted as a non-negative Boltzmann factor representing local interactions within the subensemble of the loop secondary structure (see Section 2.5 for details).

**Figure 4.**
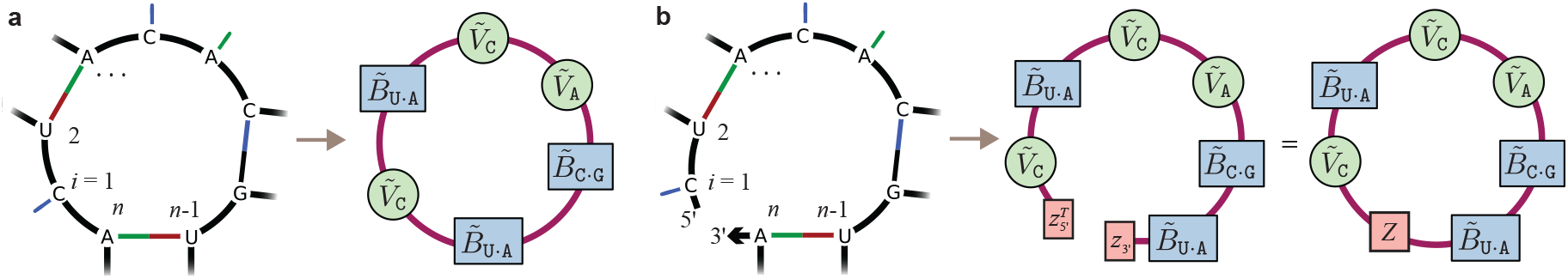
New loop-based free energy model based on transfer matrices. Nucleotides within a loop are numbered *i* = 1, …, *n*. **(a)** An example multiloop with three closing base pairs (left) and the corresponding transfer matrix network (right); cf. the periodic 1D Ising model of Figure 3a. Each unpaired base, *i*, is represented by a transfer matrix 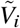 and each base pair, *i* · *i* + 1, is represented by a transfer matrix 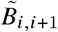. The loop free energy is given by (2.9). **(b)** An example exterior loop with three closing base pairs and a nick between 5′ and 3′ ends (left) and the corresponding transfer matrix network (middle); cf. the 1D Ising model with free ends of Figure 3b. The 5′ and 3′ ends are represented by vectors 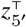 and 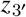. The loop free energy is given by (2.10) or equivalently by (2.11) with 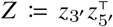. Hence, a loop with free ends can be treated equivalently as a periodic loop incorporating transfer matrix *Z* (right).

As a first example, by analogy to the periodic 1D Ising model (Figure 3a) with free energy given by (2.5), for the multiloop of Figure 4a (left), the loop free energy is calculated as (Figure 4a; right):

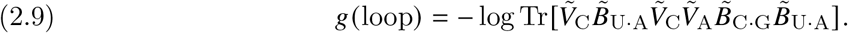

As a second example, by analogy to the 1D Ising model with free ends (Figure 3b) and free energy given by (2.8), for the exterior loop of Figure 4b (left) containing a nick before nucleotide *i* = 1, the loop free energy is calculated as (Figure 4a; middle):

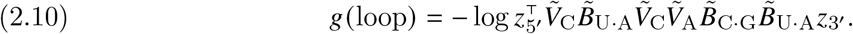

In (2.8), the role of the **1**^⊤^ and **1** vectors at the free ends is to sum all of the elements in the matrix product to yield a scalar free energy. Here, we generalize these vectors of ones to be the vectors 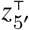 and *z*_3_′ with each entry interpreted as a non-negative Boltzmann factor representing local interactions between the end of a strand (5′ or 3′) and the adjacent entity (unpaired base or base pair). To unify the formulation and cast the free energy for the non-periodic case with free ends as the trace of a matrix product, we restate (2.10) as (Figure 4b; right):

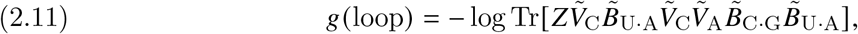

where 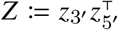 is a *γ* ×*γ* rank-one matrix with non-negative entries. This restatement exploits the fact that taking the trace of the scalar, *g*(loop), in (2.10) does not change its value and that the trace is invariant to cyclic permutations for products of vectors and matrices with compatible dimensions. Hence, the loop with free ends may be equivalently treated as a periodic loop incorporating rank-one transfer matrix *Z*, which enforces the effect of the free ends by preventing correlations between the 5′ and 3′ nucleotides on either side of the nick. As a result, for either a periodic loop (2.9) or a loop with free ends (2.11), the loop free energy, *g*(loop), may be calculated as the trace of a matrix product.

Recall that a nucleic acid secondary structure *s* may comprise a variety of loop types (Figure 1c). Exterior loops contain a single nick (with 5′ and 3′ ends on either side of the nick) and represent the non-periodic loop case with free ends. Non-exterior loops, including hairpin loops, interior loops (with stack loops and bulge loops as special cases), and multiloops, contain no nick and represent the periodic loop case.

We now formulate a single functional form for the free energy of any nucleic acid loop regardless of loop type. We calculate the loop free energy as the trace of a matrix product, where for each nucleotide *i* = 1, …, *n* in the loop we incorporate: a matrix *Z* if there is a nick before *i*, a matrix 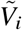 if base *i* is unpaired, a matrix 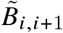 if base *i* is paired to base *i* + 1.

To arrive at a compact formulation, let us define the conditional matrix

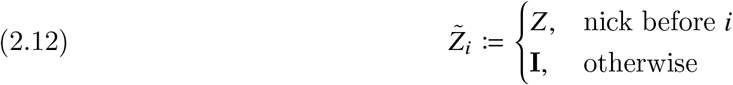

representing the presence or absence of a nick before nucleotide *i*, and the conditional matrices

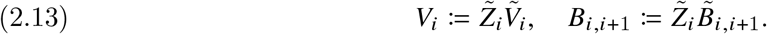

where *V*_*i*_ represents unpaired base *i* possibly preceded by a nick and *B*_*i,i*+1_ represents base pair *i* ·*i* +1 possibly preceded by a nick. With these definitions, the loop free energy may then be expressed

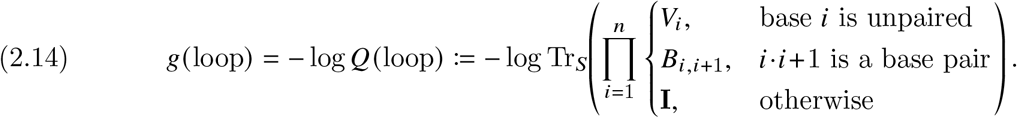

where *Q*(loop) denotes the loop partition function over the subensemble of the loop. For future reference, we use a custom trace operator, Tr_*S*_ (*w*) := Tr(*Sw*), incorporating a constant weight matrix *S* (for now, assume *S* **I**).

Let 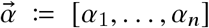 denote a list of base indices for a loop of 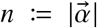 bases. For example, 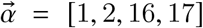 for loop 1 in the example secondary structure of Figure 5a. The secondary structure free energy, 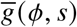, is then given by a sum of loop free energies:

**Figure 5.**
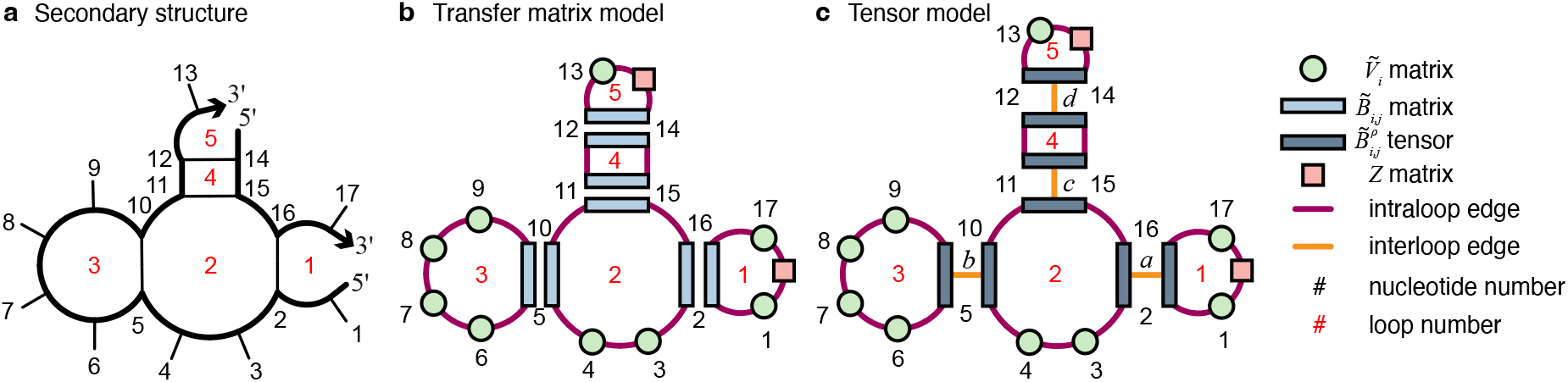
Example calculation of the secondary structure free energy, 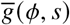, using a transfer matrix model (*ξ* = 1) or a tensor model (*ξ* ≥ 2). **(a)** Example secondary structure. **(b)** Using a transfer matrix model, the structure free energy (2.17) is a sum of independent loop free energies, where for each loop a trace contracts subensemble information to a scalar. Each intraloop edge in the transfer matrix network connects two neighboring transfer matrices within a loop. **(c)** Using the tensor model (with *ξ* ≥ 2 subtypes for each base pair), the structure free energy (2.22) is no longer a sum of independent loop free energies, where for the two neighboring loops on either side of a base pair, the base-pair subtype index (e.g., *ρ*_*a*_ ∈ [*ξ*]) appears inside the traces of the two neighboring loops, introducing a correlation between subensemble information for the two loops. Each interloop edge in the tensor network (e.g., *ρ*_*a*_, *ρ*_*b*_, …) connects two base pair tensors within neighboring loops.

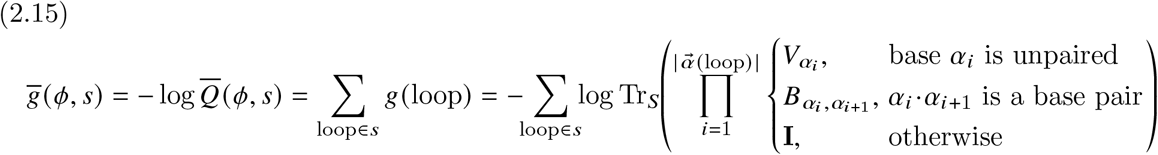

or equivalently as a product of loop partition functions:

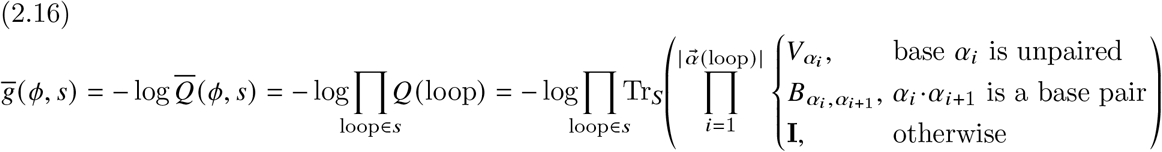

where 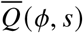 is the partition function over the subensemble of secondary structure *s*. We will later see that the (*L* − 1)*g*_assoc_ term of (1.1) can be absorbed into the loop free energies and so need not be included as an extra term in (2.15) and (2.16).

For example, for the secondary structure of Figure 5a consisting of two strands and five loops, the secondary structure free energy

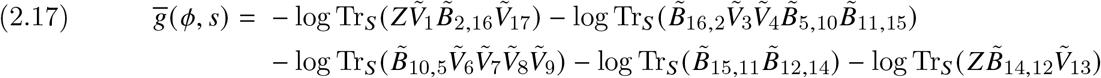

contains a sum of five independent loop free energies, each comprising a trace of a matrix product. In contrast to existing NN free energy models which employ different functional forms for different loop types and sizes and model a subensemble of coaxial and dangle stacking states only within multiloops and exterior loops, the new transfer matrix model uses a single functional form for all loop types and models a subsensemble of local interactions within all loop types.

As a limiting case of the new model, it is instructive to consider the simplest form of (2.14) when *γ* = 1. In this case, matrix operations may be replaced by their scalar analogs, and the loop free energy simplifies to a sum of a free energy contribution per nick, per unpaired base, and per base pair

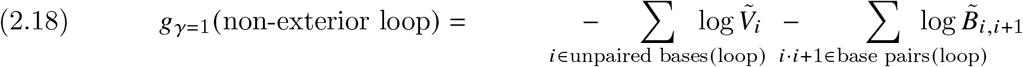

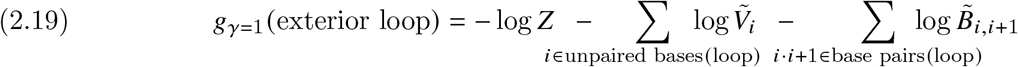

The secondary structure free energy (2.15) for *L* interacting strands simplifies to a sum of contributions per nick, per unpaired base, and per base pair:

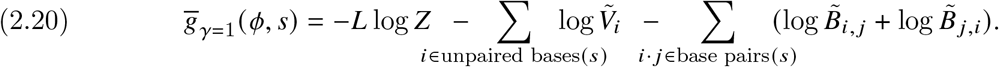

Hence, for the scalar model with *γ* = 1, there is no structural subensemble beneath secondary structure within each loop and correlations are not modeled between neighboring entities within each loop. For the transfer matrix model (*γ* ≥ 2), within each loop, each product of two matrices models local interactions between neighboring entities (each entity a nick, an unpaired base, or a base pair). Hence, correlations between neighboring entities are modeled within the subensemble of each loop. However, for each loop, the trace contracts the subensemble information for that loop to a scalar, so correlations are not modeled between the structural subensembles of different loops, which we now address.

### 2.3. Generalization to a tensor free energy model with correlated loop free energies

We now generalize the new transfer matrix model to a tensor model that enables modeling of correlations between neighboring loops within a secondary structure. In the transfer matrix model, note that each base pair *i* · *j* contributes twice to the structure free energy (using matrices 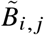 and 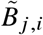), serving as a closing base pair within one loop on either side of the base pair (e.g., see Figure 5b and equation (2.17)). In essence, a single base pair is modeled using two different transfer matrices that contribute independently to two neighboring loops. This suggests the opportunity to use the base pair as a physical and logical link between the structural subensembles on either side of the base pair and thus enable modeling of correlations between neighboring loops. For this purpose, we generalize the two base pair matrices to be tensors (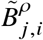and 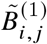) of order 3 by introducing a third index *ρ* ∈ [*ξ*] that corresponds to the addition of an interloop edge between the two tensors defining each base pair (e.g., see the interloop edges of Figure 5c).

For *ξ* = 1, we recover the transfer matrix model of Section 2.2 where the structural subensemble for each loop is modeled independently. For *ξ* ≥ 2, we may interpret *ρ* as a base-pair subtype. For example, for the neighboring loops on either side of a base pair, the respective matrices are 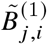 and 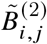 for base-pair subtype *ρ* = 1 and 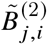 and 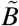 for base-pair subtype *ρ* = 2, thus enabling modeling of correlations between the structural subensembles of neighboring loops. The modeling of base-pair subtypes is supported by observations of multiple conformations for nucleic acid base pairs ^12,17^ and incorporating structural correlation across base pairs offers the opportunity to capture backbone steric effects that persist across base pairs (e.g., leading to A-, B-, or Z-form double helices) while also increasing flexibility for future work.

Using this tensor model, the free energy of a secondary structure is then

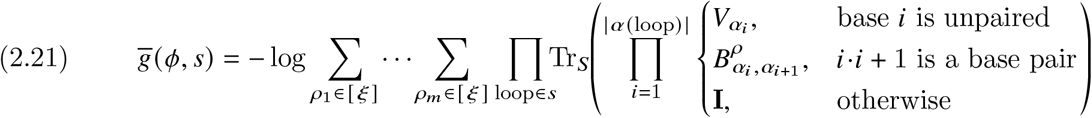

where *m* is the number of base pairs in *s* and *ρ* denotes a base-pair subtype. Compared to (2.16), the difference is that each subtype index (*ρ*_1_, *ρ*_2_, …) introduces correlations between the subensembles of a pair of neighboring loops, with summation performed over *m* ^*ξ*^ combinations of base-pair subtypes.

Revisiting the example secondary structure of Figure 5a, the tensor model of Figure 5c yields the secondary structure free energy

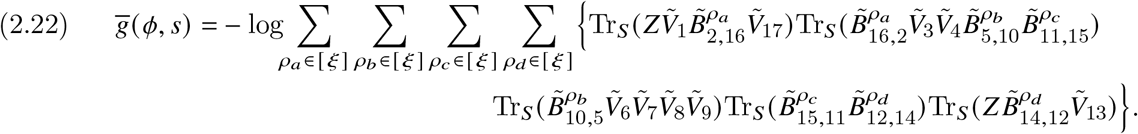

### 2.4. Tensor network contraction interpretation

For the transfer matrix model (2.16) (with independent loop subensembles) and the tensor model (2.21) (with correlated subensembles between neighboring loops), evaluation of the secondary structure free energy occurs via a set of matrix multiplications, traces, and sums. This process may alternatively be interpreted as contracting all the edges in the corresponding tensor network to produce a scalar free energy. A *tensor network* is an undirected graph of nodes, in which each node represents a tensor (of any degree ≥ 0) and each edge represents an interaction between tensors. For the tensor networks of Figures 5bc, each multiplication of two matrices or each matrix trace contracts one *intraloop edge* within a loop, and each sum over alternative base-pair subtypes (*ρ* ∈ [*ξ*]) collapses one *interloop edge* between neighboring loops, yielding a scalar secondary structure free energy. Thus, these new transfer matrix and tensor models for secondary structure free energy may be interpreted in the language of *tensor network contraction*. ^1,21^ Tensor network contraction is exploited across multiple fields including statistical mechanics, ^19^ quantum chemistry ^5,8^ and machine learning. ^26^ Theoretically, a connected tensor network can exactly express any discrete joint probability distribution given tensors of sufficiently large dimension^20^ (e.g., sufficiently large *γ* at intraloop edges and sufficiently large *ξ* at interloop edges).

For finite-dimensional tensors, the accuracy of the model improves if the topology of the network mirrors the essential physics of the problem. Base-pairing is highly non-local with respect to position in sequence, *ϕ*, so for a given secondary structure, *s*, we hard-wire non-local base-pairing information into the tensor network using 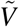 base-pair tensors, in combination with 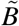 matrices to represent unpaired bases and *Z* matrices to represent nicks between strands. Beneath secondary structure, we then model a structural subensemble within each constituent loop (each loop consisting of some combination of 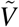 tensors, 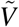 matrices, and zero or one *Z* matrices) using tensor contraction to evaluate local interactions between neighboring entities (each a nick, unpaired base, or base pair) within a loop, and optionally to model correlations between the subensembles of neighboring loops on either side of a base pair. Such local interactions can naturally encompass the existing notions of coaxial stacking and dangle stacking, but need not be explicitly defined a priori, allowing the dominant interactions to be discovered implicitly by the process of regressing to experimental data. Note that contraction of an intraloop edge models interactions that are local within a loop but generally non-local with respect to position along the sequence due to the nonlocal nature of base-pairing. Hence, the transfer matrix or tensor network layers local subensemble analysis beneath the non-local topology of the secondary structure in order to mirror the physics of the system. Unlike existing NN physical models, transfer matrix models use a single functional form for all loop types, model a structural subensemble for all loop types, and with the generalization to tensors, enable modeling of correlations between the subensembles of neighboring loops within a secondary structure.

### 2.5. Definition of a proximity state and a subensemble state within the transfer matrix and tensor models

Consider a tensor network comprising nodes connected by edges. For the 1D periodic Ising model (e.g., Figure 3a), the spin sites reside at the nodes (*i* = 1, 2, …) (each node with 2 alternative spin states, *σ*_*i*_ = ±1, corresponding to spin up or spin down) and the transfer matrices reside at the edges (e.g., *T*_1/2_, *T*_3/2_, …) between adjacent nodes. A system state is defined as a set of spin states, one per node. The partition function over the ensemble of all system states is calculated using nested sums over the alternative spin states at the nodes. The sum at node *i*, 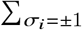, is implemented as a product of transfer matrices, *T*_*i*−1/2_*T*_*i*+1/2_that contracts node *i* in the network to yield a new effective transfer matrix between nodes *i* − 1 and *i* + 1. For product *T*_*i*−1/2_*T*_*i*+1/2_, the spin states at the different nodes are represented in the matrix entries as follows:

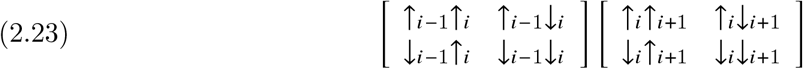

where the spin state of node *i* matches between the rows of the first matrix and the columns of the second matrix, providing the basis for contraction of node *i*. Each entry in these transfer matrices is a non-negative Boltzmann factor calculated at an edge in the network based on the interaction between the depicted pair of spin states for the nodes at either end.

For our tensor secondary structure free energy model (e.g., Figure 5c), in contrast to the 1D Ising model, transfer matrices are defined at nodes (e.g., 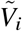 for *i* = 1, 2, …), *proximity states, σ*_*i*+1/2_ (any of *γ*), are defined at intraloop edges (e.g., *σ*_1/2_, *σ*_3/2_, …) between adjacent nodes, and base-pair subtypes *ρ* (any of *ξ*) are defined at interloop edges (e.g., *ρ*_*a*_, *ρ*_*b*_, …). A *subensemble state, s*, is defined as a set of proximity states, one per edge. The partition function over all subensemble states may be expressed using nested sums over the alternative proximity states at the edges. The sum over alternative proximity states, 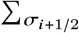, for an intraloop edge, 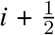, is implemented as a product of transfer matrices, for example, 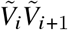, that contracts the edge 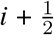 in the network to yield a new effective transfer matrix between edges 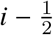 and 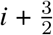. For example, if *γ* = 2 and the two proximity states at edge 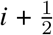 are “stack” 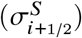 or “no stack”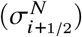, the proximity states at the different edges are represented in the matrix entries for product 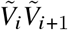 as follows:

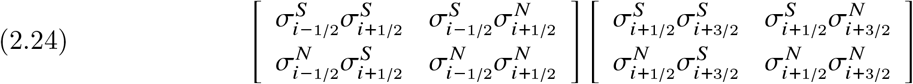

where the proximity state of edge 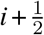 matches between the rows of the first matrix and the columns of the second matrix, providing the basis for contracting edge 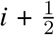. Each entry, (*a, b*), in these transfer matrices is a non-negative Boltzmann factor calculated at a node in the network based on the interaction between the pair of depicted proximity states of the edges on either side:

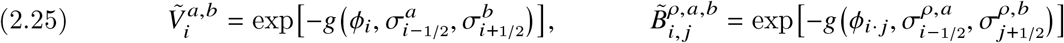

where *a* ∈ [*γ*], *b* ∈ [*γ*].

If there is a nick before node *i*, the alternative proximity states at edge 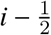 represent interactions of the 5′ end of the strand with the unpaired base or base pair at *i*. For example, if *γ* = 2, the proximity states at the different edges are represented in the vector and matrix entries for product 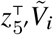 as follows:

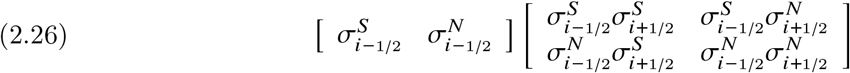

where the proximity state of edge 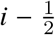 matches between the row vector and the columns of the matrix, providing the basis for contracting edge 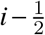. Each entry in the row vector is a non-negative Boltzmann factor calculated at the 5′ end of the strand based on the depicted proximity state of the adjacent edge:

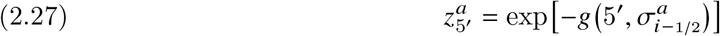

where *a* ∈ [*γ*]. Analogous observations apply for the product 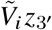 at the 3′ end of the strand. Hence, using the 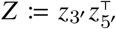 transfer matrix at a nick, the system state is defined as a set of states, one per edge, where *Z* is connected by edges to the 5′ and 3′ entities (each entity a 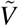 or 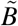 transfer matrix).

Let 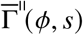 denote the ensemble of all subensemble states for secondary structure *s*. The secondary structure free energy, (2.21), may then be restated as

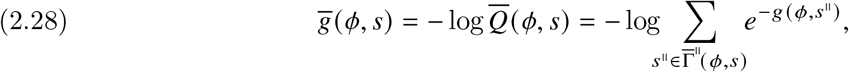

where 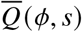 is the partition function over ensemble 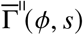 For the secondary structure ensemble, 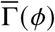, the corresponding ensemble of all subensemble states is denoted 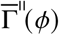. The partition function, (1.2), over ensemble 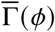, can then be restated as

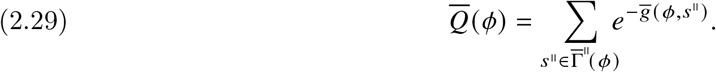

### 2.6. Incorporating physical constraints using block matrices

We now define some additional constraints on the definition of our transfer matrices and tensors in order to enhance accuracy, efficiency, and simplicity. First, for all closed loops (i.e., all non-exterior loops corresponding to those loops containing no nick), we show that logarithmic scaling of entropic cost with loop size can be accurately modeled by using a custom trace with diagonal weight matrix *S* and block diagonal 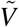 and 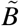 matrices (subsection 2.6.1). Second, for all exterior loops (i.e., open loops containing exactly one nick), we show that the block structure of 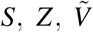 and 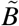 matrices can be defined so as to automatically exclude disconnected structures from the complex ensemble, enabling simple and efficient implementations (subsection 2.6.2). Third, we show that the matrix layout constraints for non-exterior loops and exterior loops can be combined into different blocks within unified transfer matrices and tensors to yield the full formulation of our transfer matrix and tensor models, with different blocks automatically contributing in the appropriate loop types.

#### 2.6.1. Approximate logarithmic entropic scaling for non-exterior loops

While tensor networks are well-suited to modeling local interactions between adjacent unpaired bases and/or base pairs within a loop, it is more challenging to model long-range effects within a loop. In particular, the entropic cost of loop closure for any non-exterior loop (i.e., a loop containing no nick) should scale logarithmically with loop size. Jacobson-Stockmayer theory^9,22^ states that the free energy associated with the entropic cost of loop closure follows the form:

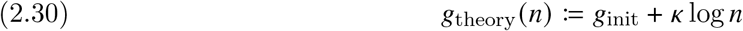

for a chain of *n* units, with *κ* = 1.5 for an ideal chain in three dimensions, and *g*_init_ a constant independent of *n*. The stiffness of nucleic acid polymers results in different empirical values of *κ*, with many RNA and DNA NN models using *κ* = 1.75. ^7,15,25,27^

It is challenging to accurately model the logarithmic scaling of (2.30) using transfer matrices of small dimension. To see this, consider the free energy of a loop of *n* entities, each an unpaired base or base pair, under the simplifying assumption that entropic effects are the same for each entity independent of sequence identity,

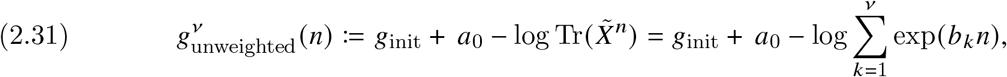

where 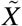 is a ν × ν matrix representing either 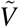 or 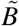. Here, we have introduced a free constant, *a*_0_, to assist with satisfying the boundary condition 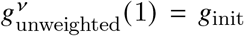. Furthermore, we have assumed that 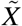 is positive-definite such that the trace may be written in terms of 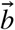, the eigenvalues of log 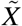; using this functional form is convenient because it guarantees the non-negativity of the eigenvalues of 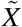 following parameter regression. Fits of (2.31) to (2.30) are shown in Figure 6a, demonstrating that the quality of fit improves only slowly with increasing ν. Note that in order to achieve *O*(*N*^3^) time complexity for dynamic programming algorithms operating on a complex of *N* nucleotides, existing NN free energy models incorporate loop free energies for multiloops that are linear in the number of unpaired nucleotides and closing base pairs (akin to *g*(*n*) := *a*_0_ − *b*_1_*n* in the terminology of (2.31)), and are unable to capture the logarithmic entropic scaling of (2.30).

**Figure 6.**
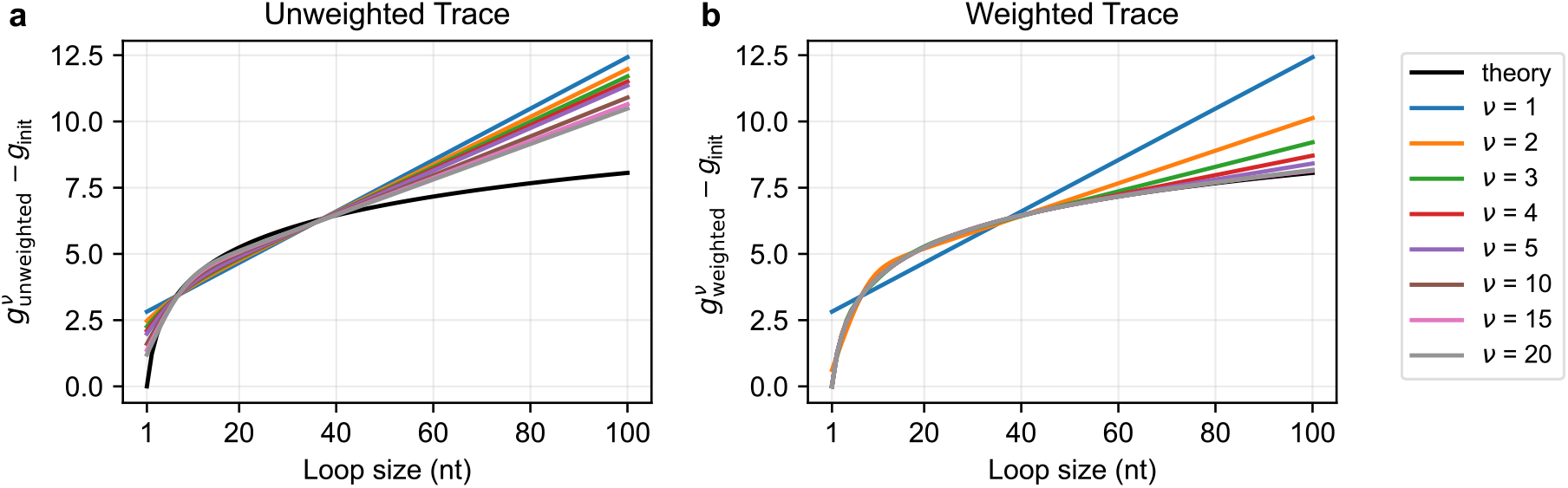
Fits of transfer matrix models of dimension ν to Jacobson-Stockmayer logarithmic scaling of entropic cost as a function of loop size for *κ* = 1.75. **(a)** Unweighted trace. **(b)** Weighted trace 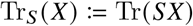 using a diagonal weight matrix *S*. Fits performed by minimizing 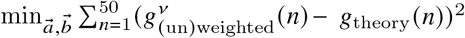 for loops of *n* = 1, …, 50 nt; results plotted beyond the fitting region. See Section S7.1 for details.

Now consider a free energy model where the loop free energy

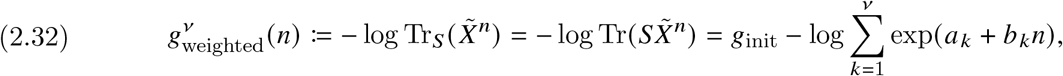

is computed using weighted trace, Tr_*S*_ (*A*) := Tr(*SA*), incorporating a constant diagonal matrix *S*. Here, 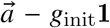 are the eigenvalues of log *S* and it is assumed that *S* and 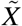 are simultaneously diagonalizable. Fits of (2.32) to (2.30) are shown in Figure 6b, demonstrating that allowing *S* ≠ **I**, the quality of fit improves rapidly with ν; in effect, each exponential is individually weighted with a free constant *a*_*i*_. We may also interpret (2.32) as defining diagonal weight and transfer matrices

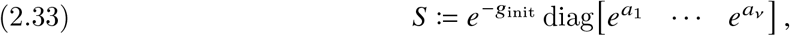

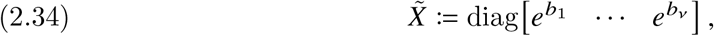

that are well-suited to approximating the non-local effect of logarithmic scaling of entropic cost for closed loops. To additionally benefit from off-diagonal transfer matrix entries for modeling local interactions, we use the Kronecker product, ⊠, to define a diagonal weight matrix and block-diagonal transfer matrix

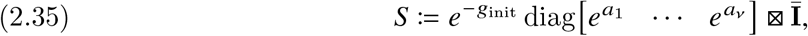

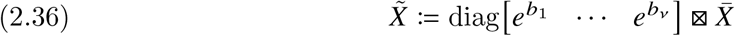

where Ī and 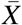 are *μ* × *μ* matrices. Noting that Tr(*A* ⊠ *B*) = Tr(*A*)Tr(*B*) we then have

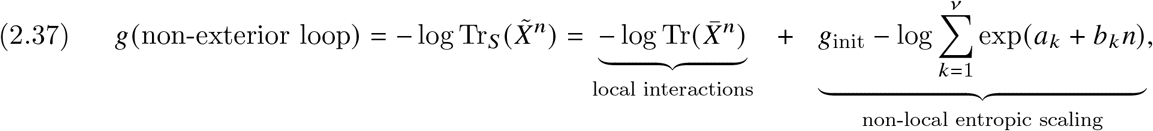

so that the weighted trace of a transfer matrix product models the loop free energy as the sum of a local term (trace of a transfer matrix product) and a non-local term that approximates the logarithmic entropic cost of loop closure (leveraging the diagonal weights of *S* and 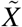).

##### Lemma 2.1.

*For weighted trace* Tr_*S*_ (*AB*) := Tr(*SAB*) *and block diagonal square matrices A and B [as for* 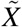 *in (2*.*36)], the necessary and sufficient condition for invariance of* Tr_*S*_ *with respect to cyclic permutations of its arguments is S diagonal with uniform constant entries α*_*k*_ *inside block k [as for S in (2*.*35)]*.

*Proof*. Necessity follows from Tr(*SAB*) ≠ Tr(*SBA*) for single-block *A* and *B* and *S* ≠ *α***I** due to the non-commutivity of the standard trace. Sufficiency follows from the fact that the blocks do not interact and that within block *k*, Tr(*S*_*k*_ *A*_*k*_ *B*_*k*_) = *α*_*k*_**I** Tr(*A*_*k*_ *B*_*k*_) = *α*_*k*_**I** Tr(*B*_*k*_ *A*_*k*_) due to the cyclic invariance of the standard trace to its arguments.

#### 2.6.2. Exclusion of disconnected structures from exterior loops

By definition, the ensemble of a complex contains only connected secondary structures, so no subset of strands in a complex of *L* strands can be free of (i.e., unpaired to) the others. For example, within a test tube ensemble containing strand species A, B, C, the 1-stranded complex *π* = A, 2-stranded complex *π* = BC, and 3-stranded complex *π* = ABC are all separate complex ensembles. By definition, all non-exterior loops within a secondary structure contain zero nicks and exterior loops contain exactly one nick. If an exterior loop contained more than one nick, that would correspond to a disconnected structure (since the strand between the two nicks would not be paired to any other strands in the complex), which is hence excluded from the complex ensemble. To keep track of the nick locations and avoid contributions from disconnected structures, previous dynamic programming algorithms for calculating the partition function over the ensemble of a complex, 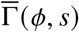, have either employed conditionals (which negatively impacted efficiency) or separate intrastrand and interstrand recursions^7^ (which complicated implementation). Here, we show that to calculate the free energy of exterior loops, interplay between the sparsity patterns of the trace weight matrix, *S*, and the transfer matrix for a nick, *Z*, can be used to automatically zero out contributions from any loop containing no nick (no *Z* matrix) or more than one nick (more than one *Z* matrix), thus excluding disconnected structures from the complex ensemble, while enabling simple and efficient implementations.

For our transfer matrix model, consider a loop consisting of a word, *w*, of transfer matrices 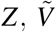, and/or 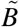(or 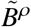 if *ξ* > 1). We would like to encode the rule that Tr_*S*_ (*w*) ≠ 0 if and only if exactly one instance of *Z* appears in *w*, thus automatically excluding contributions from non-exterior loops (that contain no *Z* matrix) and disconnected structures (that contain more than one *Z* matrix). Further, we must ensure that Tr_*S*_ (*w*) is invariant to cyclic permutations of *w* so that the calculated exterior loop free energy does not change if the transfer matrices in the loop are cyclically permuted. Consider the 2 × 2 matrices

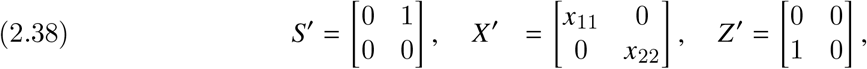

where *S*′ models the layout of *S, X*′ models the layout of both 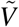 and 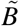, and *Z*′ models the layout of *Z*. We then have the following lemma.

##### Lemma 2.2.

*For matrices S*′, *Z*′, *and X*′ *with the sparsity patterns of* (2.38) *and w a word of Z*′ *and X*′, Tr_*S*_′ [*w*] ≠ 0 *if and only if there is exactly one instance of Z in w, and* Tr_*S*_′ [*w*] *is invariant to cyclic permutations of w if and only if x*_11_ = *x*_22_.

*Proof*. If *w* contains no *Z*′ matrices, the product *S*′*w* has the same sparsity pattern as *S*′ so 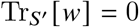. If *w* contains exactly one *Z*′ matrix, the sparsity patterns of *S*′ and *Z*′ interact to yield

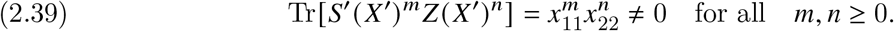

If *w* contains more than one *Z*′ matrix, the sparsity pattern of *Z*′ ensures that *w* = 0 so 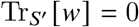. Invariance to cyclic permutations of *w* is trivial if *w* does not contain exactly one *Z* matrix; otherwise, from (2.39), invariance holds if and only if *x*_11_ = *x*_22_.

Analogous claims hold if the non-zero scalars in *S*′, *Z*′, and *X*′ are replaced with non-zero blocks. The toy matrices demonstrate how blocks within 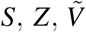 and 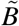 can be parametrized with sparsity patterns that make no contribution in non-exterior loops (which contain no *Z* matrices), maintain invariance with respect to cyclic permutation for exterior loops (which contain exactly one *Z* matrix), and automatically exclude contributions from disconnected loops (which contain two or more *Z* matrices).

#### 2.6.3. Full formulation of transfer matrix and tensor secondary structure free energy models

We now combine the block structures derived for non-exterior loops (Section 2.6.1) and exterior loops

(Section 2.6.2) to specify the form of the tensors using to evaluate the secondary structure free energy (2.21) (see Table 1 for a notation summary). The weight matrix, *S*, for custom trace Tr_*S*_ and transfer matrix, *Z*, for a nick are square *γ* × *γ* block matrices:

**Table 1.** Summary of transfer matrix and tensor model notation and motivations.

| Quantity | Meaning | Example physical motivation |
| --- | --- | --- |
| $\tilde{V}_i$ | Transfer matrix for unpaired base $i$ | π-π stacking |
| $\tilde{B}_{i,j}^\rho$ | Tensor for base pair $i \cdot j$ of subtype $\rho$ | H-bonds, π-π stacking |
| $Z$ | Transfer matrix for a nick | Statistical independence across nick, 5' and 3' stacking |
| $S$ | Weight matrix for custom trace $\text{Tr}_S$ | Prevent disconnected structures, weight entropic terms |
| $\gamma$ | Transfer matrix size: $\gamma = (\nu + 2)\mu$ | Model interactions at edges in tensor network |
| $\nu$ | Number of blocks for non-exterior loops | Loops containing no nicks (no $Z$ matrices) |
| $\mu$ | Block size | Subensemble states per block at each intraloop edge |
| $\xi$ | Dimension of $\rho$ tensor index | Base-pair subtypes at each interloop edge |

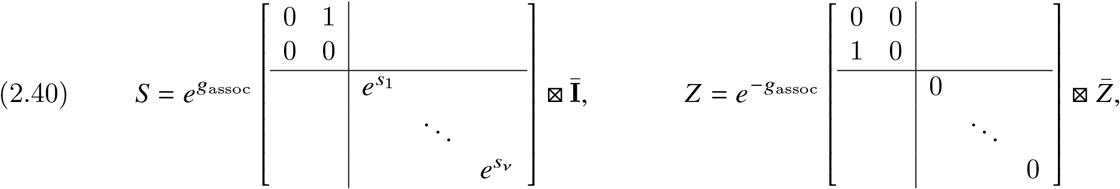

with an upper left partition for exterior loop contributions (2 × 2 blocks with the layouts of *S*′ and *Z*′ to exclude disconnected structures from the complex ensemble) and a lower right partition for non-exterior loop contributions (*S* containing ν diagonal blocks with weights 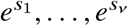). Here, Ī and 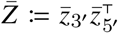 are of dimension *μ* × *μ*, so *γ* = (ν + 2) *μ*.

In what we term the *fixed-weight formulation*, the transfer matrices 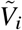 for unpaired base *i* and 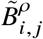 for base pair *i* · *j* are square *γ* × *γ* block-diagonal matrices with the same partitioning (upper left for exterior loop contributions and lower right for non-exterior loop contributions):

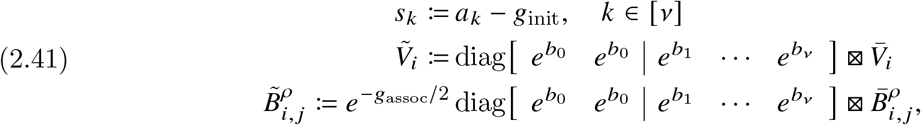

where 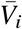 and 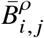 are *μ* × *μ* matrices with non-negative entries. The weighted trace block weights, 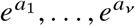 (with 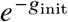 as a floating prefactor), and the transfer matrix block weights, 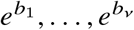, are fixed to be the values fit using (2.32) in Figure 6b to approximate logarithmic entropic cost for closed loops. Following Lemma 2.2, the block weight, 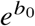, is identical in the two diagonal blocks in the top left partition to ensure invariance of the weighted trace, Tr_*S*_, to cyclic permutations of exterior loops, and *b*_0_ = 0 to zero out the loop closure term (since exterior loops are not closed).

As a basis for comparison, we also consider a more flexible *floating-weight formulation*, that has the same transfer matrix dimensions, block structure, and partitioning:

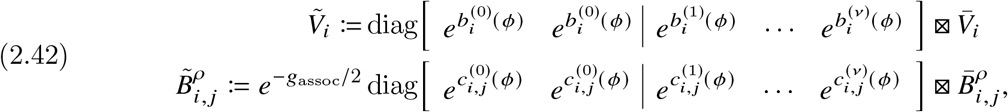

but allows the block weights 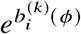 and 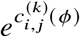, to be sequence-dependent and to float during parameter regression, with the weighted trace block weights, 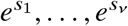, also allowed to float during regression. This formulation has the representational capacity to approximate logarithmic entropic cost of closed loops, but will exhibit this property only if it emerges during regression to the experimental data.

With either of these two formulations, for a structure, 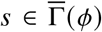, containing *L* strands, the *g*_assoc_ terms in *S, Z*, and 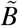 contribute (*L* − 1)*g*_assoc_ to the structure free energy (2.15), consistent with (1.1). To see this, note that there are *L* contributions of *g*_assoc_ through *Z, n*_bp_(*s*) contributions of *g*_assoc_ through 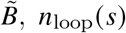 contributions of −*g*_assoc_ through *S*, and that *n*_loop_(*s*) = *n*_bp_(*s*) + 1, where *n*_bp_(*s*) and *n*_loop_(*s*) denote the number of base pairs and loops in secondary structure *s*, respectively.

The unified transfer matrices of (2.40)–(2.42) have several notable properties. Due to sparsity pattern of *S* and *Z*, the upper left partition of 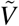 and 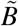 contributes to a loop free energy via (2.21) if and only if a loop is an exterior loop containing exactly one *Z* matrix (i.e., the upper left partition is not active in non-exterior loops [no *Z* matrices] or in disconnected structures [two or more *Z* matrices]). Furthermore, the lower right partition of 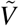 and 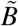 contributes to a loop free energy if and only if a loop is a non-exterior loop containing zero *Z* matrices (since *Z* contains all zeros in the lower right partition) with the capacity to approximate logarithmic entropic cost for closed loops (due to the diagonal weights in 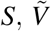 and 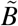).

For example, for the secondary structure of Figure 5a consisting of *L* = 2 strands and five loops, (2.17) becomes

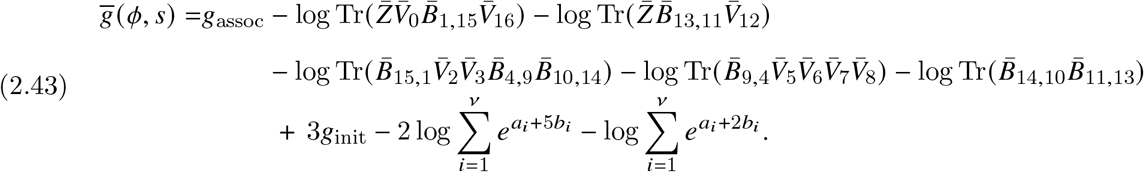

where the first row contains the strand association penalty and terms for local interactions within exterior loops 0 and 4, the second row contains terms for local interactions within non-exterior loops 1, 2, 3, and the third row contains terms for non-local entropic scaling of non-exterior loops 1, 2, 3.

For RNA, this formulation comprises 4 transfer matrices for unpaired bases 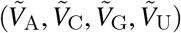 and 6 tensors for Watson–Crick and wobble base pairs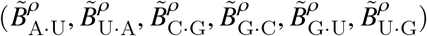. For DNA, there are 4 transfer matrices for unpaired bases 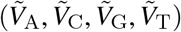 and 4 tensors for Watson–Crick base pairs (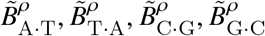; there are no wobble pairs for DNA). Conceptually (e.g., for the pseudocode of Algorithms 3.1 and 3.2), *V*_*i*_ and 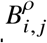 can be pre-computed for all *i* ∈ [*N*], *j* ∈ [*N*], *ρ* ∈ [*ξ*], incorporating a *Z* matrix using (2.13) if there is a nick before *i*, while also prohibiting mismatches and sterically infeasible small hairpin loops (see Section S2.6 for details); for production code, *V*_*i*_ and 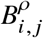 are computed on-demand to avoid incurring the storage overhead of pre-computation.

## 3. Polynomial-Time Algorithms

Having fully specified a new class of tensor free energy models (2.21) (and the subsidiary class of transfer matrix models corresponding to *ξ* = 1), we now describe efficient polynomial-time algorithms for calculating physical quantities of interest for complex ensemble 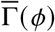 containing a total of *N* := |*ϕ*| nucleotides and the subensemble states 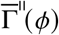. First, we describe an *O* (*N*) algorithm for calculating 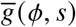, the free energy of any single secondary structure 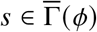 over all subensemble states 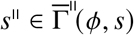) (Section 3.1). We then describe an *O*(*N*^3^) dynamic programming algorithm that uses a single recursion to calculate 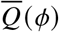, the partition function over the complex ensemble of all secondary structures 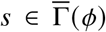 taking into account all subensemble states 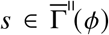) (Section 3.2). Exploiting the unified dynamic programming framework^7^ of Figure 2a, we combine this generic recursion with quantity-specific evaluation algebras and operation orders to calculate diverse physical quantities of interest (Section 3.3).

### 3.1. *O*(*N*) Algorithm for Calculating the Secondary Structure Free Energy

To calculate the secondary structure free energy 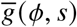 using (2.21) we must traverse each loop in secondary structure *s*, calculating the appropriate transfer matrix products and trace for each loop, while also summing over alternative base-pair subtypes if *ξ* > 1 to incorporate correlations between the subensembles of neighboring loops. Secondary structure *s* is specified as a list for *i* = 1, …, *N* where *s*_*i*_ = *i* if base *i* is unpaired and *s*_*i*_ = *j* if base *i* is paired to base *j*. Algorithm 3.1 traverses the loops in *s* via a stack data structure (last in, first out) using counter *k* to index the loops in the stack. The stack operates on element *Q*_*k*_, a *γ* × *γ* matrix representing the transfer matrix state for loop *k*. The structure *s* is traversed in a single pass along the backbone from 5′ to 3′ with the contribution for each position, *i* ∈ [*N*], handled in one of three ways (consider, for example, the secondary structure of Figure 5a corresponding to the tensor network of Figure 5c). First, if base *i* is unpaired, *Q*_*k*_ is right multiplied by *V*_*i*_ to incorporate the contribution of unpaired base *i* into loop *k*. Second, if base *i* is paired to base *j* with *i* < *j* (i.e., a base pair is opened), a new element, *Q*_*k*+1_, is pushed onto the stack to open a first base pair for a new neighboring loop *k* + 1. Third, if base *i* is paired to base *j* with *i* > *j* (i.e., a base pair is closed), the newest element on the stack is popped by closing loop *k* and incorporating this result into the ongoing calculation of the neighboring preceding loop *k* − 1. In this case, the contribution of base pair *i* · *j* is incorporated in loop *k* by left multiplying *Q*_*k*_ by 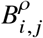 and taking the trace to complete the loop calculation, and in loop *k* − 1 by right multiplying *Q*_*k*−1_ by 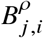; for tensor algorithms with *ξ* > 1, these contributions are summed over alternative base-pair subtypes *ρ* ∈ [*ξ*]. The contribution of a nick before *i* is incorporated by the *Z* transfer matrix included in the definition of *V*_*i*_ or 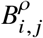 (depending on whether *i* is unpaired or paired) via (2.13). Note that in the context of the stack data structure, indices *k* − 1, *k*, and *k* + 1 provide relative indexing of the loops currently in the stack, so a given value of *k* may be pushed to open a loop, then popped upon closing the loop and pushed again to open a new loop as the backbone is traversed from 5′ to 3′ (e.g., when traversing the secondary structure of Figure 5a, index *k* = 3 is pushed and popped for loop 3 and then pushed and popped again for loop 4).

By calculating the secondary structure free energy with a single traversal of *s* for *i* ∈ [*N*], the time complexity of the algorithm is *O*(*N*) even for tensor models (*ξ* > 1) that model correlations between neighboring loop subensembles. In terms of the underlying tensor network, calculation of the scalar 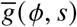 corresponds to contracting every edge in the network to yield a scalar free energy. Within each loop, each intraloop edge is contracted using a product of adjacent transfer matrices, except for the last edge, which is contracted using a trace. For tensor models (*ξ* > 1), the interloop edge between neighboring loops is contracted as a sum over alternative base pairing states. In contrast to existing NN free energy models that treat unpaired bases in an exterior loop (e.g., an unstructured strand) as a zero free energy reference state independent of length and sequence, these unstructured segments are modeled by the new transfer matrix and tensor models over a subensemble of states as a product of sequence-dependent transfer matrices.

#### Algorithm 3.1

*O*(*N*) algorithm to calculate 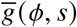, the free energy of secondary structure *s*

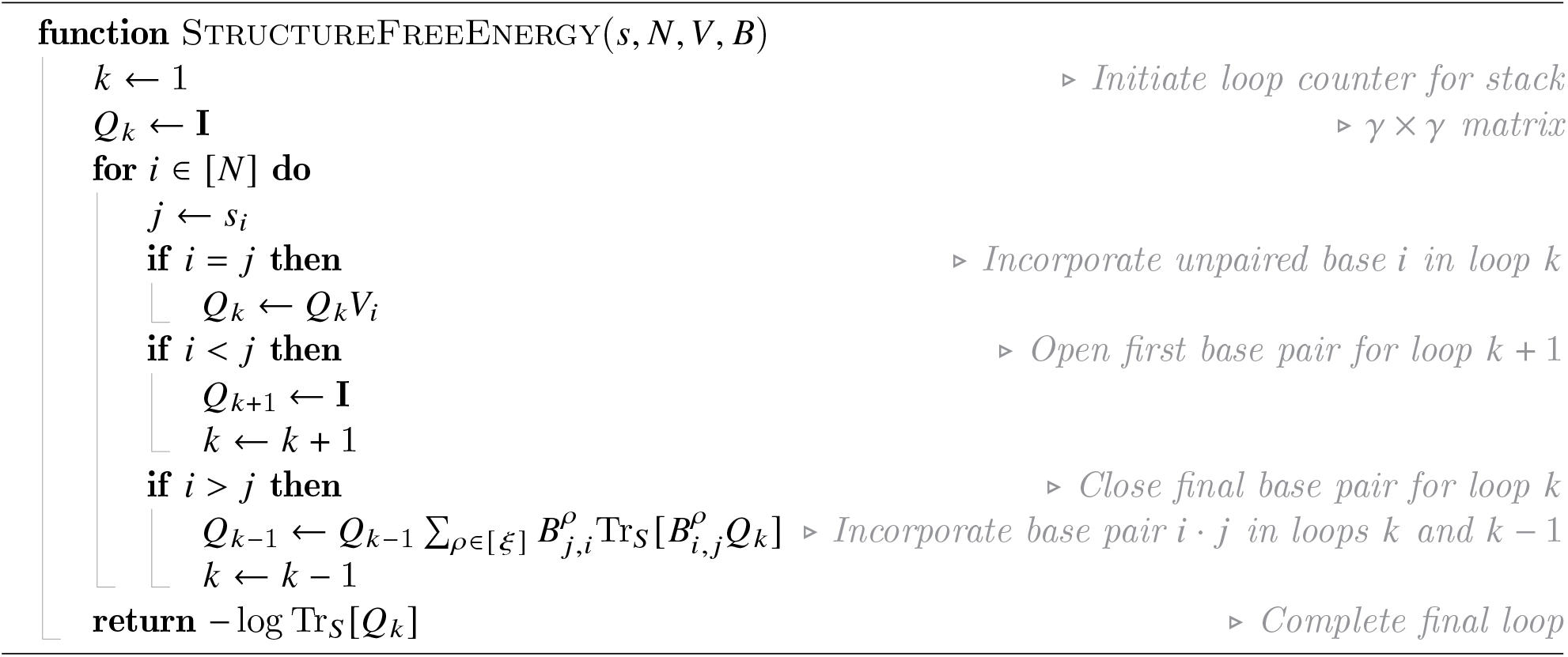

### 3.2. *O*(*N*^3^) Dynamic Programming Algorithm for Calculation of the Complex Partition Function

We now define an algorithm for calculating the partition function, 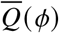, over the ensemble of all secondary structures 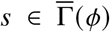 subject to the tensor secondary structure free energy model (2.21), which encompasses the subensemble states 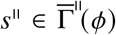. Following a dynamic programming paradigm, the algorithm calculates the partition function element, *Q*_*i, j*_, for subsequence [*i, j*], in terms of subsequence partition function elements previously calculated for shorter subsequences, ultimately yielding *Q*_1,*N*_ for the full sequence. Here, *Q*_*i, j*_ is a *γ*×*γ* matrix, and the partition function for the full sequence 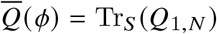. During this buildup procedure, the algorithm proceeds by incrementally adding a new 3-most base (see the operation order of Figure 7a), considering the possible base-pairing states of the new base, and incorporating the partition function contributions over the associated subensembles states. For subsequence [*i, j*] with new 3′-most base *j*, there are two scenarios: 1) either the new base *j* is unpaired, or 2) the new base *j* is base-paired to any preceding base *k* in the interval. Considering these two scenarios, calculation of the partition function element *Q*_*i, j*_ for subsequence [*i, j*] in terms of previously computed subsequence partition function elements may be represented equivalently by the *recursion diagram* of Figure 7b and by the *recursion equation*:

**Figure 7.**
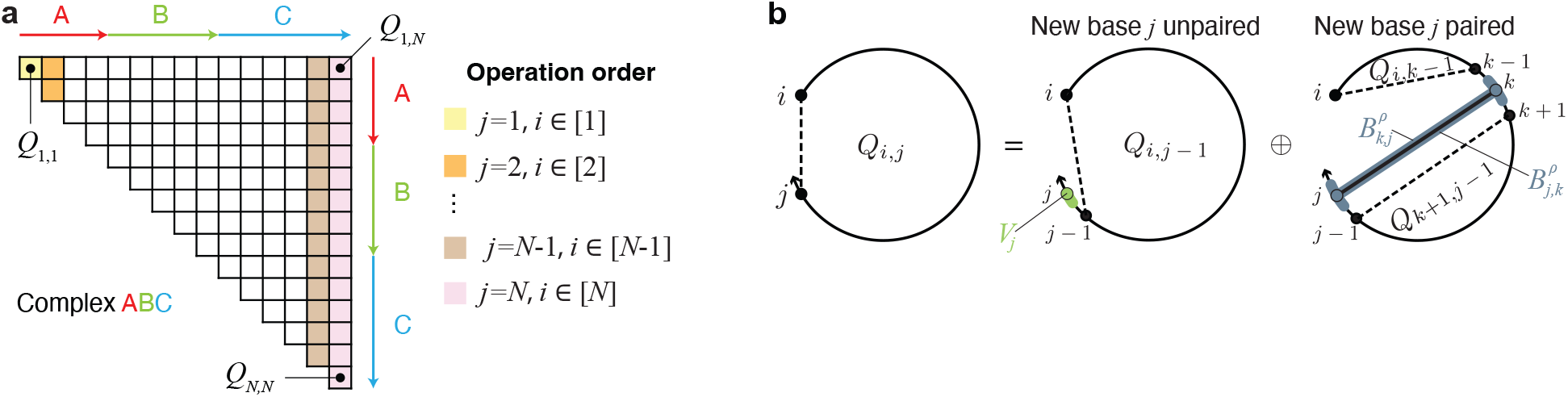
Dynamic programming algorithm for calculating the complex partition function using a single recursion (see the pseudocode of Algorithm 3.2). Calculation over ensemble 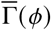 with subensemble states 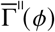 defined by the new class of tensor models (2.21) (and the subsidiary class of transfer matrix models corresponding to *ξ* = 1). **(a)** Operation order. A forward sweep of 3′-most base *j* from 1 to *N*, considering all subsequences [*i, j*] for each *j*. **(b)** Recursion diagram corresponding to the recursion equation (3.1). *Q*_*i, j*_ respresents the partition function element for subsequence [*i, j*]. A solid straight line indicates a base pair and a dashed line demarcates a region without implying that the connected bases are paired. Shading corresponds to transfer matrices and/or tensors (*V* or *B*) incorporated at the current level of recursion. There are two cases: either the new base *j* is unpaired (incorporating a transfer matrix *V*_*j*_ and making use of previously computed subsequence partition function *Q*_*i, j*−1_) or the new base *j* forms a base pair with any preceding base *k* in the interval (in-corporated by the tensors 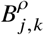 and 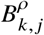 into the neighboring loops on either side of the base pair and making use of previously computed subsequence partition functions *Q*_*k*+1, *j*−1_ and *Q*_*i,k*−1_ in the respective loops).

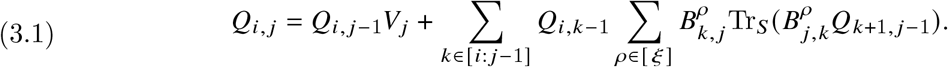

In the first scenario, where the new base *j* is unpaired, the transfer matrix *V*_*j*_ right multiplies *Q*_*i, j*−1_, the previously computed partition function element for subsequence [*i, j* −1]. In the second scenario, where the new base *j* is paired to any preceding base *k* in the interval, the tensors 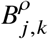 and 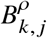 are incorporated into the neighboring loops on either side of base pair *k* · *j*, making use of previously computed subsequence partition functions *Q*_*k*+1, *j*−1_ and *Q*_*i,k*−1_ in the respective loops. On one side of the base pair *k* · *j*, tensor 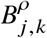 left multiplies *Q*_*k*+1, *j*−1_ and closes a loop, with 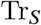 contracting the final edge in the loop to yield a scalar, and on the other side of base pair *k* · *j, Q*_*i,k*−1_ left multiplies 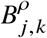 to build up partition function information for a new loop that will be closed in a future recursion level. Summation is performed over each alternative base-pair subtype *ρ* ∈ *ξ* for each alternative base *k* ∈ [*i* : *j*−1]. The recursion implicitly incorporates the contributions of a nick in an exterior loop via the definitions of *V*_*i*_ and 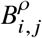 using (2.13), which conditionally include a *Z* matrix if there is a nick before *i*. The interplay of the sparsity patterns of the *S* and *Z* matrices automatically excludes contributions from loops containing more than one nick, which would correspond to disconnected structures that are not in the complex ensemble.

#### Algorithm 3.2

*O*(*N*^3^) algorithm to calculate the complex partition function 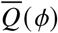.

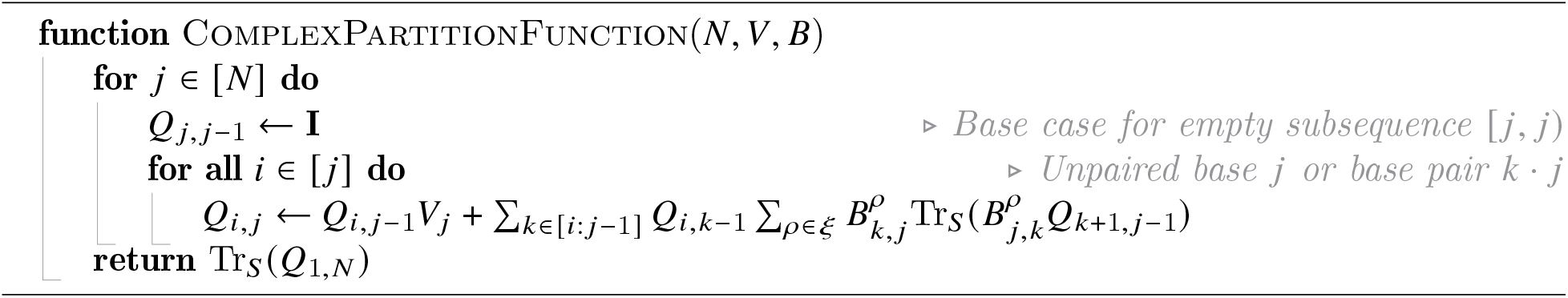

Algorithm 3.2 displays pseudocode. The time complexity of the algorithm, which we will improve below, is *O*(*ξγ*^3^*N*^3^) (base indices *i, j, k* and base-pair subtype index *ρ* on the right hand side with matrix multiplication a *γ*^3^ operation) and the space complexity is *O*(*γ*^2^*N*^2^) (base indices *i, j* on the left hand side with each recursion element, *Q*_*i, j*_, a *γ* × *γ* matrix).

Remarkably, only the single recursion (3.1) is needed to exactly calculate the complex partition function in *O*(*N*^3^) over ensemble 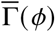 subject to the tensor free energy model (2.21). This is possible because the tensor free energy model treats all loop types using a single functional form. For production code, the time complexity can be reduced from *O*(*ξγ*^3^*N*^3^) to *O*(*γ*^3^*N*^3^ + *ξγ*^2^*N*^2^) by evaluating the inner sum of (3.1) using an intermediate recursion so that there is no asymptotic dependence on *ξ* as *N* → ∞ (see Section S4.2 for details). Moreover, the new models and algorithms account for subensemble states within all loop types and can approximate the logarithmic entropic scaling in all closed loops (i.e., all non-exterior loops). By contrast, existing state-of-the-art partition function algorithms that operate over complex ensemble 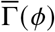 subject to existing NN free energy models require more than 20 recursions to treat the piecemeal functional forms used for different loop types and sizes with efficient vectorizable algorithms; furthermore, existing algorithms incorporate a subensemble of stacking states only in multiloops and exterior loops, and omit logarithmic entropic scaling in multiloops in order to enable polynomial-time complexity. ^7^

Note that existing dynamic programming algorithms operate on a subsequence partition function element, *Q*_*i, j*_, that is the scalar partition function of subsequence [*i, j*], whereas the new dynamic programming algorithm operates on a subsequence partition function element, *Q*_*i, j*_ that is a *γ* × *γ* matrix, corresponding to partition function contributions for subsequence [*i, j*]. In contrast to existing algorithms that make a forward sweep from short subsequences to long subsequences (operation order of Figure 2b), the new algorithm makes a forward sweep of the 3′-most base *j* from 1 to *N* (operation order of Figure 7a). This approach offers two important advantages for optimized implementation of production code. First, it facilitates straightforward prevention of numerical overflow using a per-base weighting factor (see Section S10.2). Second, the sole component of the algorithm that contributes *O*(*N*^3^) complexity can be written as a single global matrix product, enabling straightforward GPU acceleration for the first time (see Section S6.3.2).

### 3.3. A unified dynamic programming framework for calculating diverse physical quantities

The unified dynamic programming framework^7^ of Figure 2a defines a dynamic program to calculate a physical quantity of interest using three ingredients: a set of model-specific recursions, a quantity-specific evaluation algebra, and a quantity-specific operation order. The recursion equation of (3.1) enables calculation of the complex partition function, 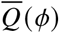, subject to the physical model comprising the structural ensemble 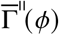and the tensor free energy model (2.21). We now abstract this quantity-specific (partition function) recursion to yield a generalized recursion equation

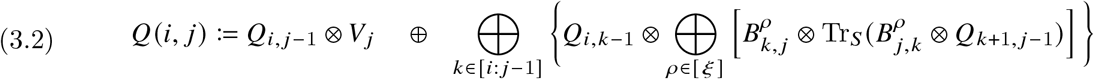

employing the generalized weighted trace

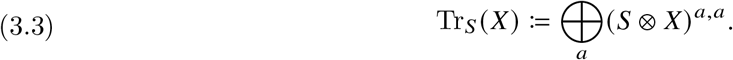

This single generic recursion (which specifies the dependencies between subproblems and incorporates the details of the structural ensemble and free energy model) can be combined with a quantity-specific evaluation algebra (to specify the mathematical form of each subproblem) and a quantity-specific operation order (to define the computational trajectory through the dependency graph of subproblems) to yield a dynamic program for calculating diverse physical quantities of interest (Table 2).

**Table 2.**
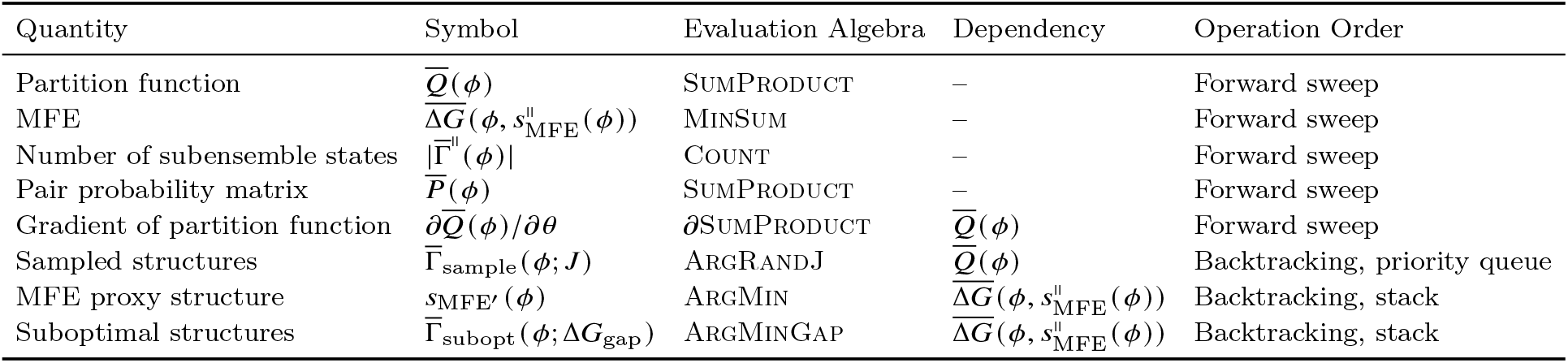
Algorithmic ingredients for calculating diverse physical quantities.

An evaluation algebra ^7^ operating on transfer matrix or tensor recursion elements, is an algebraic structure comprising: 1) a semiring *R* equipped with binary matrix operators ⊕ and ⊗ and an identity matrix, I, of dimension *γ* containing 1 on the diagonal and 0 off the diagonal, 2) a map *W* (*g*) from free energy, *g*, to *R* with the property *W* (0) = 1, and 3) a map *Q*(*i, j*) from recursion indices (*i, j*) to a *γ* × *γ* matrix in *R*. Using an evaluation algebra, transfer matrices and tensors are defined element-wise by the quantity-specific map, *W* (*g*), so the partition function-specific definitions of (2.25) and (2.27) become:

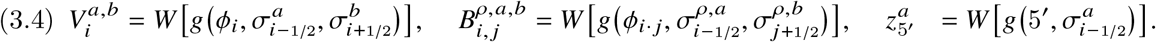

to enable calculation of different physical quantities.

Table 3 displays evaluation algebras for calculating diverse physical quantities over the complex ensemble. For example, the partition function is calculated using the SumProduct evaluation algebra: 1) ⊕ is standard matrix addition, ⊗ is standard matrix multiplication, 0 is 0, 1 is 1; 2)*W* (*g*) := exp(−*g*) is the Boltzmann factor with the property *W* (0) = 1; 3) *Q* is the trivial lookup operator *Q*(*i, j*) ↦→ *Q*_*i, j*_ that returns a *γ* × *γ* matrix corresponding to recursion indices (*i, j*). Alternatively, the free energy, 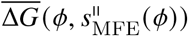, of the minimum free energy (MFE) subensemble state, 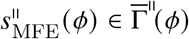, can be calculated using the MinSum evaluation algebra: 1) ⊕ reinterprets matrix addition with each subsidiary scalar addition replaced by scalar minimization, ⊗ reinterprets matrix multiplication with each subsidiary scalar multiplication replaced by scalar addition and each subsidiary scalar addition replaced by scalar minimization, 0 is ∞, 1 is 0; 2) *W* (*g*) := *g* is the free energy with the property *W* (0) = 0; 3) *Q* is the trivial lookup operator *Q*(*i, j*) ↦→ *Q*_*i, j*_. Either evaluation algebra can be used to define the mathematical form of the subproblems displayed in the generic recursion diagram of Figure 7 and recursion equation (3.1). Compared to calculating the partition function using SumProduct, when calculating the MFE using MinSum, every sum of alternative partition function contributions is replaced with a minimization over alternative free energy contributions and every product of exponentiated free energies is replaced with a sum of free energies. The same operation order is used to calculate the partition function or the MFE (Figure 2b and Table 2), performing a forward sweep in which the 3′ base is incremented from 1 to *N*.

**Table 3.**
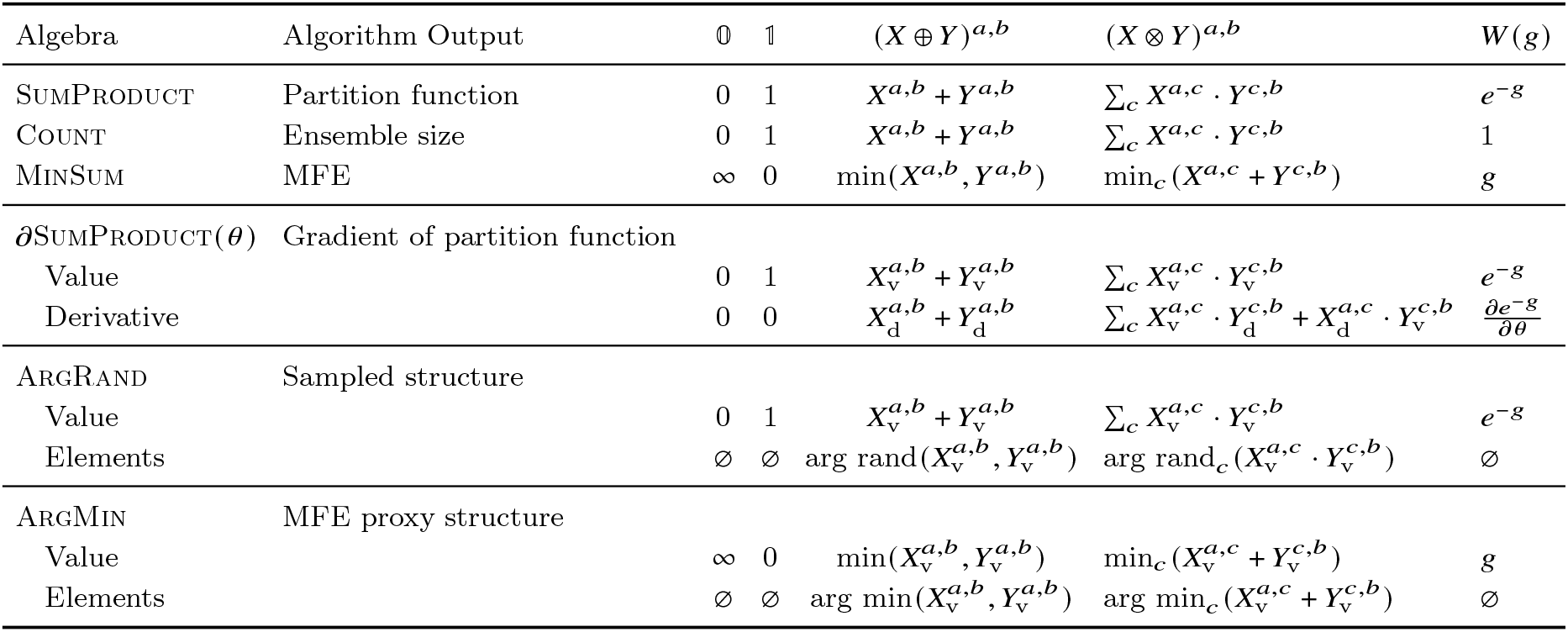
Evaluation algebras for dynamic programming algorithms operating on a complex ensemble. Operations are performed for all matrix entries with indices (*a, b*) within a given evaluation algebra domain. SumProduct yields the partition function. Count yields the number of subensemble states. MinSum yields the free energy of the MFE subensemble state. *∂*SumProduct yields the gradient of the partition function with respect to an arbitrary parameter, with each element a partition function contribution 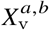 and the associated derivative contribution 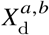. ArgRand yields a Boltzmann sampled secondary structure, with each element a partition function contribution 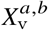 and the associated recursion elements 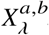. ArgMin yields the MFE proxy structure containing the MFE subensemble state, with each element a free energy contribution 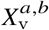 and the associated recursion elements 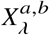. See Section S5 for details.

Table 3 presents evaluation algebras for calculating additional physical quantities including the MFE proxy structure *s*_MFE_′ (defined as the secondary structure containing the MFE subensemble state), the set of suboptimal proxy structures 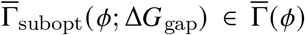 containing subensemble states within a specified Δ*G*_gap_ of the MFE subensemble state, the set of *J* Boltzmann-sampled secondary structures, 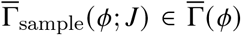, and the gradient of the partition function, 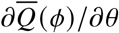, with respect to an arbitrary parameter *θ* (which provides the basis both for calculating fundamental physical quantities and for gradient-based parameter regression without the use of finite differences). These evaluation algebras are combined with quantity-specific operation orders (see Section S6 for details) to efficiently traverse the recursive subproblems, performing either a forward sweep or else backtracking through a dependency tree of previously computed recursion elements.

## 4. Model and algorithm performance

We now assess the representational capacity, accuracy, scalability, and speed of these new transfer matrix and tensor models and algorithms.

### 4.1. Representational capacity

Existing NN free energy models employ different functional forms and lookup tables for different loops types and sizes, requiring extensive documentation to describe all the functional forms and edge cases. As a reality check, we were curious whether the new transfer matrix models, which use a single functional form (trace of a matrix product) for all loop types and sizes, are flexible enough to encompass the representational capacity of the piecemeal NN models. To provide a holistic assessment, we regressed model parameters by optimizing the Kullback-Leibler (K-L) divergence (D_KL_) for a range of transfer matrix models using either the fixed-weight formulation (2.41) or the floating-weight formulation (2.42) relative to reference NN free energy models for RNA and DNA. The non-negative D_KL_ objective function was defined based a *box ensemble* containing random sequences interacting in a dilute solution (see Section S8 for details). As a basis for comparison, we also calculated the K-L divergence between historical and reference NN models for a given material (*D*_KL_ ≈ 0.5). Figure 8 demonstrates that RNA transfer matrix models achieve the representational capacity of existing NN models (*D*_KL_ < 0.5) as the number of blocks, ν, and block size, *μ*, are increased (see Figure S10 for the corresponding DNA results). The flexible-weight formulation is able to more closely approximate the existing NN reference model than the historical NN model of the same material, while the fixed-weight model is somewhat constrained from approaching the existing NN reference model given that it seeks to approximate logarithmic entropic costs for closed loops, a feature not present in the existing NN reference model for small hairpin or interior loops or for multiloops of any size.

**Figure 8.**
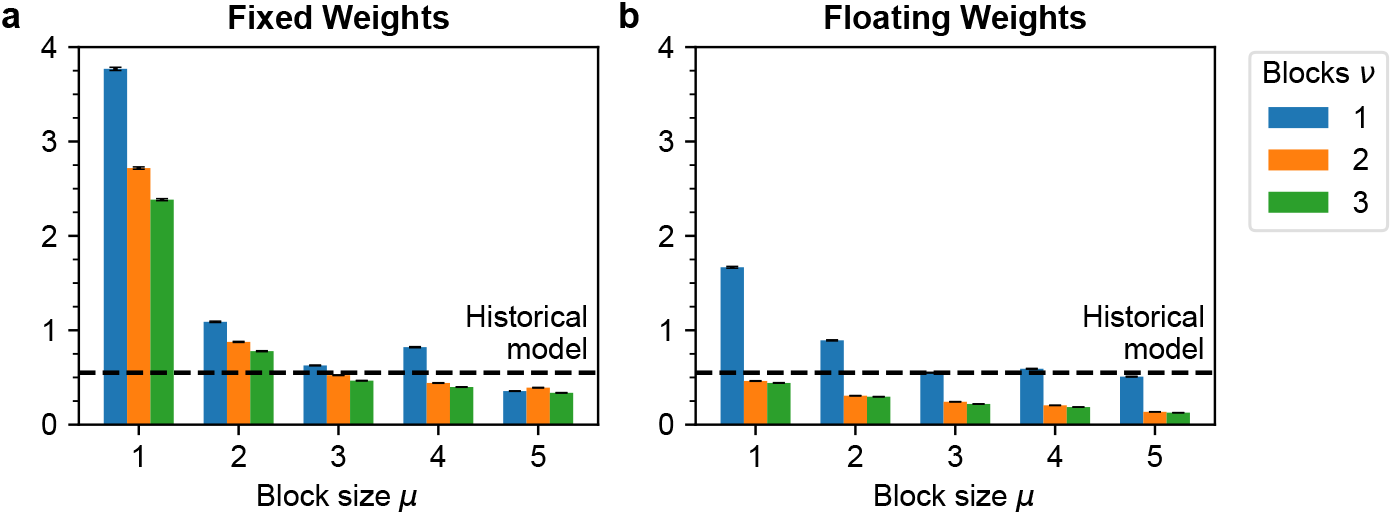
Representational capacity of new RNA transfer matrix models relative to NN reference model: a) Fixed-weight formulation (2.41). b) Floating-weight formulation (2.42). *D*_KL_ for test set (mean ± SEM over 100,000 box ensembles) using parameters optimized on training set of 180,000 box ensembles for transfer matrix models with ν ∈ {1, 2, 3} blocks and *μ* ∈ {1, 2, 3, 4, 5} block size with *ξ* = 1 (no correlation between loop subensembles). Dashed lines represent mean *D*_KL_ for a historical NN model on the test set. Reference model: `rna06` (RNA) with `stacking` subensemble. Historical model: `rna95` (RNA) parameters with `no stacking` subensemble. See Figure S10 for corresponding DNA results.

### 4.2 Accuracy

Historically, the parameters for existing RNA and DNA NN free energy models were regressed based on a large number of melt experiments for different sequences. ^13,15,24,30,32^ Here, to regress model parameters and assess accuracy, we collected large data sets of literature melt experiments for RNA (895 melts) and DNA (464) in standard salt conditions (1.0 M Na^+^). For each material, we assigned ≈80% of experiments to a training set to use for parameter regression, and the other ≈20% of experiments to a test set to assess model accuracy as well as to monitor overfitting. For each melt experiment, the literature reports the reaction free energy and enthalpy, 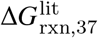(at reference temperature *T*_37_ := 310.15 K) and 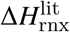, based on a fit to the absorbance melt curve. For a given transfer matrix or tensor model, calculated values of the reaction free energy and enthalpy, 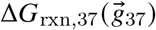 and 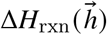, were used to regress the free energy and enthalpy model parameters, 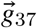 and 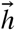. Temperature dependent free energy parameters are then given by

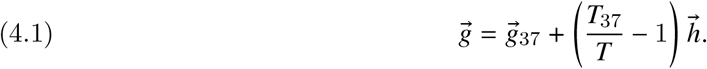

We regressed parameters for the fixed-weight formulation (2.41) and floating-weight formulation (2.42) for a range of transfer matrix and tensor models with different (ν, *μ, ξ*) values (see the RNA models of Table S8). Based on the literature melt data, we monitored the root mean square error (RMSE) for the test set divided by that of the training set, seeking ratios lower than ≈1.1 as a proxy for avoidance of overfitting. Figure 9 illustrates RNA model accuracy relative to experiment for an existing NN reference model (panel a) and for representative transfer matrix models ν, *μ, ξ* = (2, 2, 1) using the fixed-weight formulation (panel b) or the floating-weight formulation (panel c) for reaction free energy (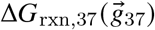; top) and reaction enthalpy (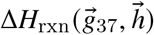; bottom). There are several notable points. First, relative to the existing NN reference model, accuracy is improved using the fixed-weight formulation (test set MAE decreases from 0.9 to 0.8 kcal/mol for reaction free energy and from 11.1 to 8.9 kcal/mol for reaction enthalpy) or the floating-weight formulation (test set MAE decreases from 0.9 to 0.7 kcal/mol for reaction free energy and from 11.1 to 8.0 kcal/mol for reaction enthalpy). Second, the existing RNA NN model uses 12,596 parameters comprising 553 unique values compared to 51 parameters for the fixed-weight formulation and 77 parameters for the floating-weight formulation. Third, accuracy tends to improve significantly with increasing block size (*μ* = 2 for the selected model), but only modestly when increasing the number of blocks (ν = 2 for the selected model). Fourth, Figure 10 displays the entropic cost of loop closure for multiloops, revealing that the fixed-weight formulation (panel a) closely approximates the theoretical logarithmic scaling as ν increases, but that the floating-weight formulation (panel b) optimizes away from logarithmic scaling to enhance accuracy relative to the experimental data. Fifth, over-fitting is observed as ν, *μ*, and *ξ* are increased in various combinations (see the test/training ratios in Table S8). Sixth, models with *ξ* > 1 that use different base-pair subtypes to model correlation between structural subensembles in neighboring loops led to marginal accuracy improvements for some models (Table S8), motivating the choice *ξ* = 1 based on the current experimental training set; future regression with larger experimental training sets may motivate adoption of models with *ξ* > 1 that model loop correlations. For all of the above points, similar trends are evident for the corresponding DNA models (Table S10 and Figures S12 and S9), with DNA models being more susceptible to overfitting due to use of a smaller experimental training set.

**Figure 9.**
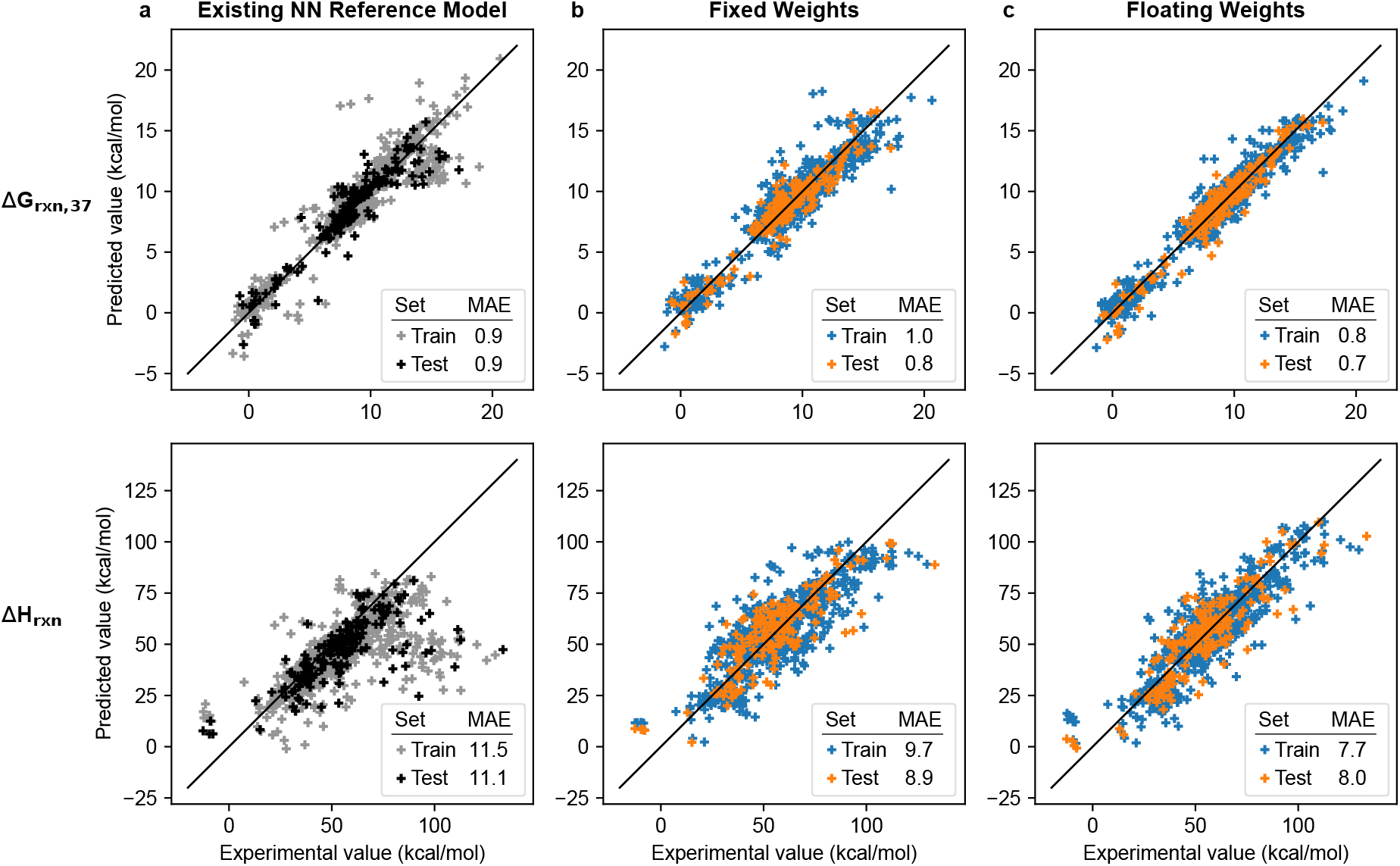
Predicted vs experimental RNA reaction free energies (Δ*G*_rxn,37_; top) and enthalpies (Δ*H*_rxn_; bottom). a) Existing nearest-neighbor (NN) reference model (panel a; NUPACK 4.1.0.1 with rna06 parameters and `stacking` subensemble). b,c) New transfer matrix models (ν, *μ, ξ*) = (2, 2, 1) using either the fixed-weight formulation (panel b) or floating-weight formulation (panel c). Mean absolute error (MAE) in kcal/mol. Training set: 716 melts. Test set: 179 melts. Salt conditions: 1.0 M Na^+^. See Section S9.1 for additional details.

**Figure 10.**
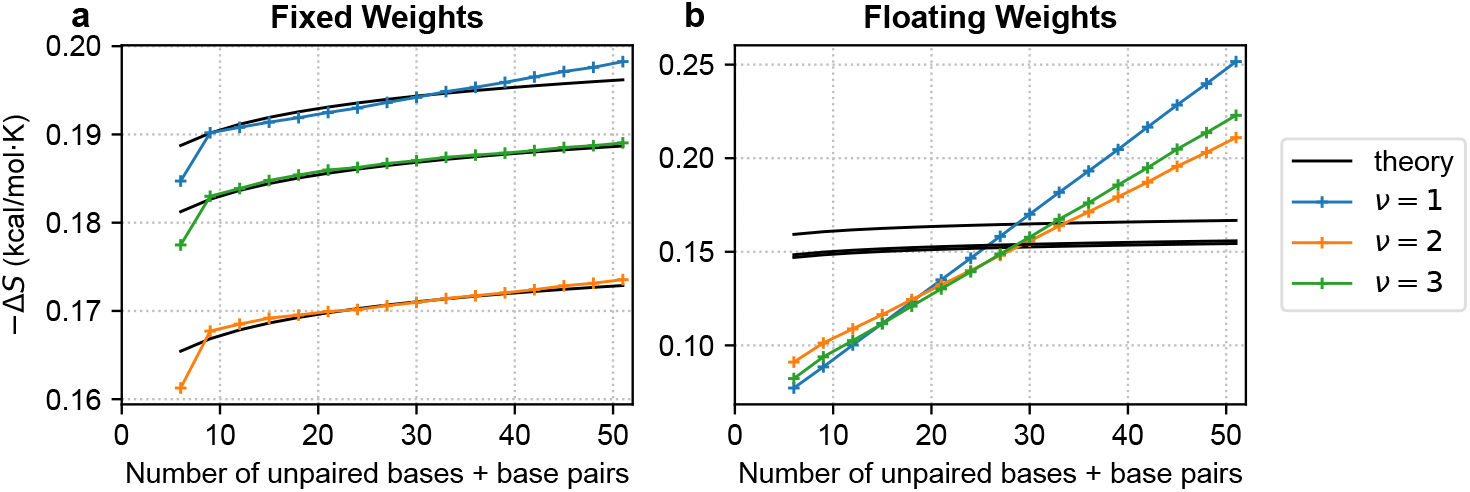
Assessment of logarithmic scaling of entropic cost with loop size for RNA multiloops using transfer matrix models with: a) fixed-weight formulation, b) floating-weight formulation. Loop size defined as the number of transfer matrices in the loop transfer matrix network (i.e., number of unpaired bases + base pairs). 100 random sequences for each combination of loop type and size. Means over 10,000 random sequences for each loop size. Theory depicted forL*κ* = 1.75. Fit of theory curve to transfer matrix results performed for each ν by minimizing 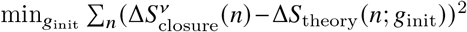 over loop size *n* with respect to free theory parameter *g*_init_. Transfer matrix model: ν, *μ, ξ* = (2, 2, 1). Conditions: RNA, 37 ^°^C, 1 M Na^+^. See Section S7.2 for details.

### 4.3. Scalability

One of the challenges with calculating the partition function is the prevention of overflow ^6,7^ as the size of the complex, *N* := |*ϕ*|, increases. The maximum expressible numbers for single-precision (32-bit), double-precision (64-bit), and quad-precision (128-bit) arithmetic, are ≈ 10^38^, ≈ 10^308^, and ≈ 10^4932^, leading to partition function overflow for single-stranded RNA (ssRNA) at ≈240 nt, ≈1700 nt, and ≈27,000 nt, respectively. For random double-stranded RNA (dsRNA), which has a deeper well in the free energy landscape, partition function overflow occurs for smaller complexes (≈70 nt, ≈560 nt, and ≈8700 nt, respectively). To overcome these limitations, the renormalization approach of Section S10.2 enables straightforward overflow-safe partition function calculations with negligible effect on cost. Figure 11 demonstrates partition function calculations for complexes of up to 50,000 nt for either ssRNA or dsRNA. By comparison, existing algorithms based on NN free energy models achieve scalable partition function calculations by employing an overflow-safe evaluation algebra that operates separately on the mantissa and exponent, leading to a ≈2–2.5× increase in cost. ^7^

**Figure 11.**
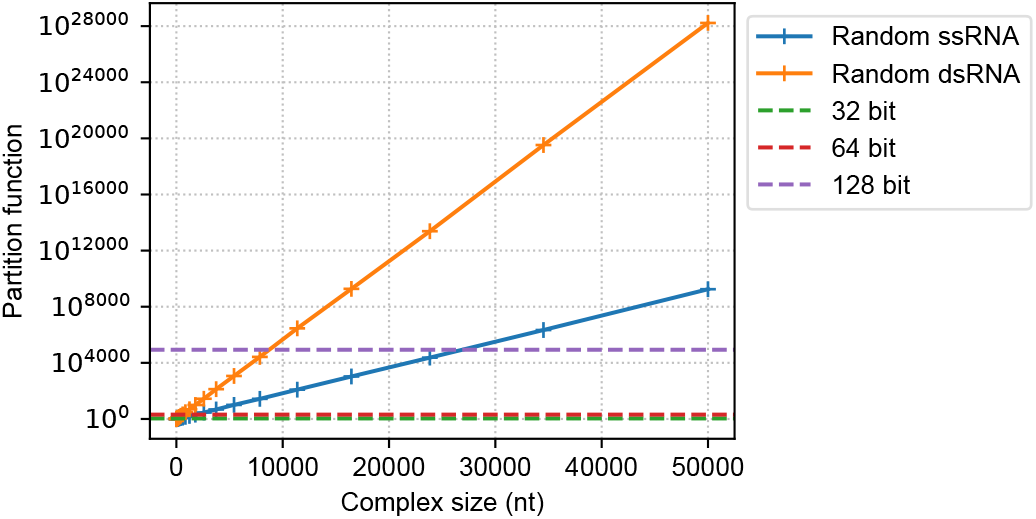
Overflow-safe partition function calculations on complex ensembles including very large complexes. Dashed lines denote the overflow thresholds for single-, double-, and quadruple-precision arithmetic. Partition function calculations performed using renormalization (Algorithm **??**). Random single-stranded RNA (ssRNA) test set: 1-stranded complex with random sequence. Random double-stranded RNA (dsRNA) test set: 2-stranded complex comprising a random sequence and its complement. Geometric means of partition functions over 3 sets of sequences per complex size. Transfer matrix model: fixed-weight formulation with ν, *μ, ξ* = (2, 2, 1). Conditions: RNA, 37 ^°^C, 1.0 M Na^+^. See Section S10.2 for details.

### 4.4. Speed

Figure 12a profiles the cost of partition function calculations using a new transfer matrix dynamic program on complex ensembles up to 30,000 nt. The production implementation blends CPU parallel execution for small complexes with GPU execution for large complexes. For large complexes, GPU acceleration leads to an order of magnitude speedup relative to CPU parallel execution of the transfer matrix algorithm or the existing vectorized NN partition function algorithm (Figure 12b). Figure S14 illustrates the relative cost of calculating the partition function, equilibrium pair probability matrix, and minimum free energy (MFE) of a subensemble state using the evaluation algebras and operation orders of Table 2. Using production implementations (blended CPU/GPU), the cost of calculating the equilibrium pair probability matrix is ≈2–3× the partition function cost (Figure S14b). Relative to CPU parallel execution of the partition function algorithm, the MFE cost using CPU parallel execution is ≈2–3× higher (Figure S14c). Relative to the production partition function algorithm (blended CPU/GPU), the MFE cost using parallel CPU execution (GPU implementation not available) increases with complex size (e.g., ≈30× higher for 10,000 nt; Figure S14b).

**Figure 12.**
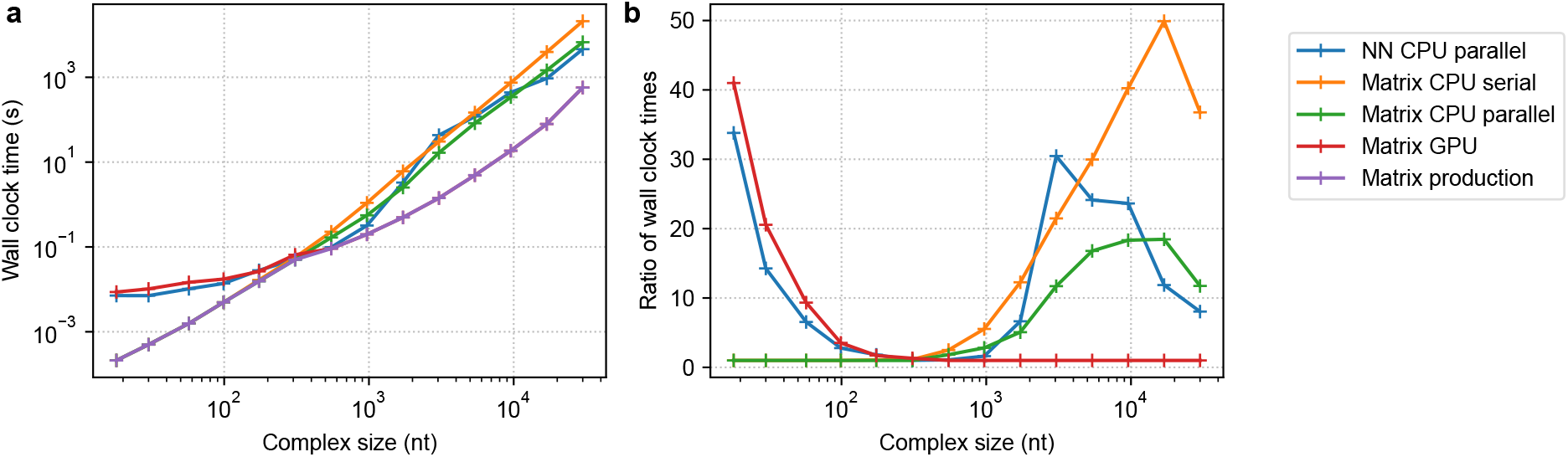
Speed of partition function calculations on complex ensembles up to 30,000 nt. (a) Computational cost. (b) Computational slowdown relative to production transfer matrix algorithm. Transfer matrix model: fixed-weight formulation with ν, *μ, ξ* = (2, 2, 1) (subsensemble of states in all loop types); one of four algorithm implementations: CPU serial, CPU parallel, GPU, or production (blend of CPU parallel for small complexes and GPU for large complexes). Existing NN reference model and algorithm: CPU parallel implementation (vectorized, overflow-safe evaluation algebra, coaxial and dangle stacking subensembles in multiloops and exterior loops; NUPACK 4.1.0). Calculation of the partition function for a complex of 3 strands, each with a different random sequence of uniform length. Conditions: RNA, 37 ^°^C, 1.0 M Na^+^.

## 5. Conclusions

This paper introduces transfer matrix models as a fundamentally new approach to modeling nucleic acid secondary structure thermodynamics. In a major departure from existing nearest-neighbor (NN) models, which use different functional forms for different loop types and sizes within a secondary structure, the new models use the trace of a matrix product as a single functional form for all loops. As a further contrast, a structural subensemble is modeled beneath secondary structure for all loop types. Additionally, by generalizing transfer matrices to tensors, we enable modeling for the first time, of correlations between the structural subensembles of neighboring loops. Introducing a custom weighted trace operator and a block structure to transfer matrices and tensors enables approximation of logarithmic entropic scaling in all closed loops (a feature lacking in multiloops using NN models) and automatic exclusion of disconnected structures in open loops (i.e., exterior loops containing exactly one nick). A secondary structure, *s*, for a complex with sequence, *ϕ*, corresponds to a transfer matrix or tensor network, with evaluation of the secondary structure free energy, 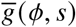, corresponding to contraction of all edges in the network to yield a scalar free energy. The single functional form of the new transfer matrix and tensor models enables a single dynamic programming recursion to replace over 20 loop-type-specific NN recursions to achieve *O*(*N*^3^) calculation of the complex partition function, 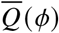, for a complex containing *N* nucleotides. Furthermore, this simplicity enables the partition function dynamic program to be expressed as a global matrix multiplication primitive that enables straightforward GPU acceleration, yielding order-of-magnitude speedups for large complexes. The single generic recursion encapsulating the details of the structural ensemble and free energy model can be combined with quantity-specific evaluation algebras and operation orders to calculate diverse physical quantities over the complex ensemble (equilibrium base pairing probabilities, number of subensemble states, minimum free energy proxy structures, Boltzmann-sampled secondary structures, and gradient of the partition function).

Despite their simplicity, transfer matrix models have the representational capacity to closely reproduce existing NN RNA and DNA models. Benchmarking new models on experimental RNA and DNA data, the new models exhibit accuracy as good as or better than that of existing NN models, using dozens of parameters rather than the hundreds or thousands used in NN models. For the experimental training and test sets employed here, use of tensors to model correlations between the structural subensembles of neighboring loops did not yield compelling accuracy benefits. Additional effort is warranted using larger and more comprehensive experimental training and tests sets to more fully explore the optimal formulation and accuracy of transfer matrix and tensor models. If justified by future experimental data, the number of blocks *ν*, block size *μ*, and number of base-pair subtypes *ξ*, can be increased over time to increase the number of subensemble states and the expressive power of the model. Here, we formulated transfer matrix and tensor models for nucleic acid secondary structure thermodynamics by defining a transfer matrix, *V*, for each unpaired base type (4 types for RNA) and a transfer matrix or tensor, *B*, for each base-pair type (6 types for RNA). However, within this new modeling framework, there are many other alternative formulations that merit consideration, including: defining transfer matrices at the edges rather than the nodes of the secondary structure graph (e.g., at the edge between unpaired bases or between an unpaired base and a paired base), defining a transfer matrix for each of the two individual bases within a base pair (e.g., one matrix for C and one matrix for G within a C·G base pair), defining blocks of different sizes within a transfer matrix or tensor (in contrast to using a uniform block size *μ*), defining a transfer matrix for each tuple of bases instead of for individual bases (e.g., for sequence ACGUU defining one matrix for each tuple ACG, CGU, and GUU). Overall, the new transfer matrix and tensor models represent a major simplifying departure from the piecemeal approach of existing NN models, providing the foundation for a new generation of modeling and algorithmic innovation in analyzing and designing nucleic acid systems.

## Acknowledgments

We thank M. Gonzalvo for help gathering literature experimental data and comments on the manuscript, J. Huang for implementation advice, N.J. Porubsky for discussions on stacking recursions and backtracking operation orders, A. Nanjundiah for discussions of literature experimental data, C.W. Wang for comments on the manuscript, and M.J. Lindsey for discussions of tensor methodologies.

## Notation

Subsequence [*i, j*] denotes bases *i*, …, *j* and subsequence [*i, i* −1] denotes an empty subsequence. In equations, *i* ∈ [*N*] denotes *i* = 1, …, *N, k* ∈ [*i*:*j*] denotes *k* = *i*, …, *j*, and *k* ∈ [*i*: *i*−1] denotes *k* = ∅. In pseudocode we use “**for all**” to indicate unordered iteration and “**for**” to indicate necessarily ordered iteration.

## Implementation

Algorithms are coded in C++17 making use of the `OpenBLAS` library ^2,33^ for CPU implementation, the `cuBLAS` library^18^ for GPU implementation, and the armadillo matrix template library^23^ for other matrix operations. To enhance efficiency, all operations on the (sparse) tensors 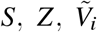, and 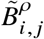 of (2.40)–(2.42) are implemented using explicit block representations, which facilitates parallelization.

## Computation

Divergence fitting regressions were run on `g3.xl` instances (AMD Milan 7713 CPU, NVIDIA A100 SXM4 40GB GPU) using the Jetstream2 cluster^28^. All other computations were run on the NERSC Perlmutter cluster (CPU: AMD EPYC 7763 with up to 64 cores and 256 GB RAM, GPU: 2× Nvidia A100 with 80 GB RAM).

## SOFTWARE

Algorithms can be run interactively via the NUPACK web app (nupack.org) or scripted locally using the NUPACK Python module, subject to the NUPACK software license agreement.^*^ Please direct questions, comments, feature requests, and bug reports to

## Supporting information

Additional model details, diverse physical quantities, recursions details, evaluation algebras, operation orders, logarthmic scaling of the entropic cost of loop closure, representational capacity to reproduce existing nearest-neighbor models, regression of model parameters based on experimental melt data, additional studies, algorithm validation, data files.

## Footnotes

* NUPACK updates employing models and algorithms from the current work will be made available in conjunction with publication in a peer-reviewed journal.

