## Supporting Information for "Transfer Matrix and Tensor Approaches to Nucleic Acid Secondary Structure Thermodynamics^*^"

### List of Algorithms

S1 Optimized $O(N^3)$ algorithm to calculate the complex partition function $Q(\phi)$ with reduced time complexity prefactor. . . . . . . 12
S2 Optimized $O(N^3)$ algorithm to calculate the complex partition function $\overline{Q}(\phi)$ with reduced time-complexity prefactor using production code conventions (cf. main text conventions of Algorithm S1 and $\chi$ conventions of Algorithm S3). . . . . . . 22
S3 Optimized $O(N^3)$ algorithm to calculate the complex partition function $\overline{Q}(\phi)$ with reduced time-complexity prefactor using $\chi$ conventions (cf. main text conventions of Algorithm S1 and production code conventions of Algorithm S2). . . . . . . 22
S4 Renormalization for prevention of partition function overflow using main text conventions (cf. production code conventions of Algorithm S5 and $\chi$ conventions of Algorithm S6) . . . . . . 23
S5 Renormalization for prevention of partition function overflow using production code conventions (cf. main text conventions of Algorithm S4 and $\chi$ conventions of Algorithm S6) . . . . . . 24
S6 Renormalization for prevention of partition function overflow using $\chi$ conventions (cf. main text conventions of Algorithm S4 and production code conventions of Algorithm S5). . . . . . . 24
S7 Optimized $O(N^3)$ algorithm to calculate the complex partition function $\overline{Q}(\phi)$ with renormalization to avoid overflow and efficient linear algebra primitives with optional GPU acceleration using main text conventions (cf. production code conventions of Algorithm S8 and $\chi$ conventions of Algorithm S9). . . 25

S8 Optimized $O(N^3)$ algorithm to calculate the complex partition function $\overline{Q}(\phi)$ with renormalization to avoid overflow and efficient linear algebra primitives with optional GPU acceleration using production code conventions (cf. main text conventions of Algorithm S7 and $\chi$ conventions of Algorithm S9). . . 26
S9 Optimized $O(N^3)$ algorithm to calculate the complex partition function $\overline{Q}(\phi)$ with renormalization to avoid overflow and efficient linear algebra primitives with optional GPU acceleration using $\chi$ conventions (cf. main text conventions of Algorithm S7 and production code conventions of Algorithm S8). . 27
S10 Backtrack-free $O(N^3)$ algorithm to calculate the equilibrium pair probability matrix $\overline{P}(\phi)$. . . . . . . 29
S11 Simultaneous Boltzmann sampling . . . . . . . . . . . . . . . . . . . . . . . . . . . . . . . . . . . . . . 32
S12 Suboptimal structure generation . . . . . . . . . . . . . . . . . . . . . . . . . . . . . . . . . . . . . . 35
S13 Stochastic gradient descent with Hessian information . . . . . . . . . . . . . . . . . . . . . . . . . . 43
S14 Trust region radius update . . . . . . . . . . . . . . . . . . . . . . . . . . . . . . . . . . . . . . . . . 52
S15 Exhaustive enumeration of all secondary structures $s \in \overline{\Gamma}(\phi)$ for sequence $\phi$ . . . . . . . . . . . . . . . 61

### S1 Presentation of models and algorithms

The supplementary information of Fornace et al.[19] provides a detailed description of a unified dynamic programming framework for calculating diverse physical quantities over complex and test tube ensembles using nearest-neighbor (NN) models and algorithms. To facilitate comparison, here we follow the same presentation as closely as possible, modifying text, equations, algorithms, and figures as necessary to describe the new transfer matrix and tensor models and algorithms.

### S2 Additional details for transfer matrix and tensor free energy models

#### S2.1 Definition of Watson–Crick and wobble base pairs

Base pairs are either Watson–Crick pairs:

- RNA: rG·rC and rA·rU,
- DNA: dG·dC and dA·dT,

or wobble pairs:

- RNA: rG·rU,

Note that for DNA, dG and dT are considered to form a mismatch dG×dT and not a wobble pair.[59]

#### S2.2 Dimensionless free energy

For a complex with sequence $\phi$, the (dimensional) free energy of secondary structure, $s \in \overline{\Gamma}(\phi)$, treating all strands as distinct, is

$$\overline{\Delta G}(\phi, s) = \overline{\Delta H}(\phi, s) - T\overline{\Delta S}(\phi, s), \tag{S1}$$

where $\overline{\Delta H}(\phi, s)$ and $\overline{\Delta S}(\phi, s)$ are assumed to be independent of temperature. To simplify notation, models and algorithms are presented in terms of the dimensionless free energy

$$\overline{g}(\phi, s) := \overline{\Delta G}(\phi, s)/(k_B T), \tag{S2}$$

where $k_B$ is the Boltzmann constant and $T$ is temperature.

#### S2.3 Temperature dependence of the free energy model

For the new transfer matrix and tensor models, the free energy parameters, $\vec{g}_{37}$, at reference temperature $T_{37} :=$ 310.15 K, and enthalpy parameters, $\vec{h}$, assumed to be temperature independent, are regressed (see Section S9 for details) and the temperature-dependent free energy parameters are calculated using

$$\vec{g} = \vec{g}_{37} + \left(\frac{T_{37}}{T} - 1\right)\vec{h}. \tag{S3}$$

In this equation, terms are non-dimensionalized as follows:

$$\begin{aligned} g &= \Delta G/(k_B T), \\ g_{37} &= \Delta G_{37}/(k_B T_{37}), \\ h &= \Delta H/(k_B T_{37}). \end{aligned} \tag{S4}$$

where $\Delta G_{37} := \Delta H - T_{37}\Delta S$. Note that parameter vectors, $\vec{g}_{37}$ and $\vec{h}$, are non-dimensionalized at a known fixed temperature, $T_{37}$, so that they can be stored in parameter files. On the other hand, the temperature-dependent parameter vector, $\vec{g}$, is non-dimensionalized on the fly at a user-specified temperature, $T$, for straightforward calculation of $\tilde{V}$ and $\tilde{B}$ using (2.25).

##### S2.4 Distinguishability issues

For a complex of $L$ strands, the ensemble $\overline{\Gamma}(\phi)$ treats each strand as distinct while the ensemble $\Gamma(\phi)$ treats strands with the same sequence as indistinguishable.[13,19] Both ensembles have conceptual utility as they provide different perspectives when examining the physical properties of a complex. In laboratory experiments, strands with the same sequence are typically indistinguishable, so calculations over ensemble $\Gamma(\phi)$ are crucial for comparison to experimental data (e.g., equilibrium secondary structure probabilities and equilibrium complex concentrations). On the other hand, calculations over ensemble $\overline{\Gamma}(\phi)$ can sometimes provide information that is valuable precisely because it cannot be measured experimentally (e.g., equilibrium base-pairing probability matrix).

For a secondary structure $s \in \Gamma(\phi)$ with an $R$-fold rotational symmetry there is an $R$-fold reduction in distinguishable conformational space, so the secondary structure free energy, $\overline{g}(\phi, s)$, given by (2.15) or (2.21) must be adjusted by a symmetry correction:[13]

$$g(\phi, s) = \overline{g}(\phi, s) + g^{\mathrm{sym}}(\phi, s) \tag{S5}$$

where

$$g^{\mathrm{sym}}(\phi, s) = \log R(\phi, s). \tag{S6}$$

All of the dynamic programs described in the present work operate on ensemble $\overline{\Gamma}(\phi)$ using free energy model (2.15) or (2.21) where each strand is treated as distinct. This is a matter of algorithmic necessity, as the free energy model (S5) used for ensemble $\Gamma(\phi)$ contains a symmetry correction that depends on the global rotational symmetry $R$ of each secondary structure $s \in \Gamma(\phi)$. For efficiency reasons, the dynamic programs avoid explicitly enumerating each structure, instead operating on local loop free energies to incorporate information for multiple structures simultaneously while operating only on local loop free energies. As a result, the dynamic programs cannot incorporate a different global rotational symmetry correction for each structure because they never have access to global structural information. However, to facilitate comparisons to experimental data, physical quantities calculated by a dynamic program using physical model (2.15) or (2.21) over ensemble $\overline{\Gamma}(\phi)$ can be post-processed by leveraging the Distinguishability Correction Theorem of Dirks et al.[13] to enable exact conversion to physical quantities calculated using physical model (S5) over ensemble $\Gamma(\phi)$; see Section S5 of Reference 19 for details. Interestingly, ensembles $\overline{\Gamma}(\phi)$ and $\Gamma(\phi)$ both have utility when examining the physical properties of a complex as they provide related but different perspectives, akin to complementary thought experiments.

##### S2.5 Floating-block formulation for transfer matrix and tensor models

As a basis for comparison, we also consider a more flexible *floating-block formulation* that further generalizes the floating-weight formulation of (2.42). As for the floating-weight formulation, $S$ and $Z$ are defined by (2.40) with diagonal weights, $e^{s_1}, \ldots, e^{s_\nu}$, in $S$ permitted to float during regression. The definitions for $\tilde{V}$ and $\tilde{B}$ have the same partitioning (upper left for exterior loop contributions and lower right for non-exterior loop contributions) and block structure:

$$\tilde{V}_i = \left[\begin{array}{cc|ccc} V_i^{(0)} & & & & \\ & V_i^{(0)} & & & \\ \hline & & V_i^{(1)} & & \\ & & & \ddots & \\ & & & & V_i^{(\nu)} \end{array}\right], \qquad \tilde{B}^\rho_{i,j} = e^{-g^{\mathrm{assoc}}/2} \left[\begin{array}{cc|ccc} B_{i,j}^{\rho,(0)} & & & & \\ & B_{i,j}^{\rho,(0)} & & & \\ \hline & & B_{i,j}^{\rho,(1)} & & \\ & & & \ddots & \\ & & & & B_{i,j}^{\rho,(\nu)} \end{array}\right]. \tag{S7}$$

but with the diagonal blocks in $\tilde{V}$ and $\tilde{B}$ permitted to float during regression so that each block can have different non-negative values. This formulation has the representational capacity to approximate logarithmic entropic cost of closed loops, but will exhibit this property only if it emerges during parameter regression. Following Lemma 2.2, the two diagonal blocks in the top left partition of $\tilde{V}$ are identical (and likewise for $\tilde{B}$) to ensure invariance of the custom trace, $\mathrm{Tr}_S$, to cyclic permutations of exterior loops.

##### S2.6 Transfer matrix and tensor definitions for bases that cannot pair

For RNA, there are 4 transfer matrices for unpaired bases ($\tilde{V}_{\mathrm{A}}, \tilde{V}_{\mathrm{C}}, \tilde{V}_{\mathrm{G}}, \tilde{V}_{\mathrm{U}}$) and 6 tensors for Watson–Crick and wobble base pairs ($\tilde{B}^\rho_{\mathrm{A\cdot U}}, \tilde{B}^\rho_{\mathrm{U\cdot A}}, \tilde{B}^\rho_{\mathrm{C\cdot G}}, \tilde{B}^\rho_{\mathrm{G\cdot C}}, \tilde{B}^\rho_{\mathrm{G\cdot U}}, \tilde{B}^\rho_{\mathrm{U\cdot G}}$). For DNA, there are 4 transfer matrices for unpaired bases

$(\tilde{V}_{\mathrm{A}}, \tilde{V}_{\mathrm{C}}, \tilde{V}_{\mathrm{G}}, \tilde{V}_{\mathrm{T}})$ and 4 tensors for Watson–Crick base pairs ($\tilde{B}^{\rho}_{\mathrm{A\cdot T}}, \tilde{B}^{\rho}_{\mathrm{T\cdot A}}, \tilde{B}^{\rho}_{\mathrm{C\cdot G}}, \tilde{B}^{\rho}_{\mathrm{G\cdot C}}$; there are no wobble pairs for DNA).

For any two bases that form a mismatch (e.g., A×A, A×C, C×A) the corresponding $\tilde{B}$ tensor is set to $\mathbb{0}$ (e.g., $\tilde{B}^{\rho}_{\mathrm{A\times A}} = \mathbb{0}$, $\tilde{B}^{\rho}_{\mathrm{A\times C}} = \mathbb{0}$, $\tilde{B}^{\rho}_{\mathrm{C\times A}} = \mathbb{0}$). This way, the evaluation algebra will appropriately prevent contributions from bases that cannot base pair during the dynamic programming recursions without using conditionals to check for base-pairing feasibility.

Even if $\phi_i$ and $\phi_j$ are suitable for forming a base pair (e.g., $\phi_i$ = C, $\phi_j$ = G), if formation of the base pair would close a hairpin loop, they can only base pair if $|j - i| \geq 4$ because the smallest feasible hairpin loop has 3 unpaired nucleotides in the loop.[41,59] Hence, in dynamic programming recursions, if $|j - i| < 4$ and subsequence $[i, j]$ contains no nick (i.e., formation of the base pair would lead to a hairpin loop rather than an exterior loop), we set $\tilde{B}^{\rho}_{i,j} = \mathbb{0}$ so as to prevent contributions from infeasible hairpin loops.

#### S2.7 Definition of model parameter sets and number of model parameters

As described in Section 2.6.3, the $\gamma \times \gamma$ trace matrix $S$, $\gamma \times \gamma$ transfer matrices $\tilde{V}$, and $\gamma \times \gamma \times \xi$ tensors $\tilde{B}$ (which reduce to transfer matrices if $\xi = 1$), are defined to have block structures as follows:

- Fixed-weight formulation: block structure defined by (2.40) and (2.41).
- Floating-weight formulation: block structure defined by (2.40) and (2.42).
- Floating-block formulation: block structure defined by (2.40) and (S7).

A given transfer matrix or tensor model is defined as a tuple $(\nu, \mu, \xi)$ where $\gamma = (\nu + 2)\mu$. The number of subensemble states that are modeled, and the number of parameters that are used to do so, can be adjusted by changing the following:

- $\nu$ (the number of blocks in a non-exterior loop): increasing $\nu$ is expected to increase the capacity of the model to approximate logarithmic scaling of entropic cost with loop size in non-exterior loops (i.e., all loops that are closed and do not contain a nick).
- $\mu$ (block size): increasing $\mu$ is expected to increase the capacity to model local interactions between neighboring nucleotides and to propagate information between next-nearest neighbors, next-next-nearest neighbors, etc.
- $\xi$ (number of base pair subtypes): increasing $\xi$ is expected to increase the capacity to model correlations between the structural subensembles of neighboring loops on either side of a base pair.

The number of base types, $n_{\mathrm{basetype}}$ is 4 for RNA (A,C,G,U) or DNA (A,C,G,T) and the number of base pair types $n_{\mathrm{bptype}}$ is 6 for RNA (A·U, U·A, C·G, G·C, G·U, and U·G) and 4 for DNA (A·T, T·A, C·G, G·C). The number of model parameters for different formulations are itemized by type in Table S1 with totals as follows:

$$|\vec{g}| = (n_{\mathrm{basetype}} + \xi n_{\mathrm{bptype}})\mu^2 + 2(\nu + \mu + 1) + 1 \qquad \text{fixed-weight formulation} \qquad \text{(S8)}$$

$$|\vec{g}| = (n_{\mathrm{basetype}} + \xi n_{\mathrm{bptype}})\mu^2 + 2\mu + (1 + n_{\mathrm{basetype}} + n_{\mathrm{bptype}})(\nu + 1) \qquad \text{floating-weight formulation} \qquad \text{(S9)}$$

$$|\vec{g}| = (n_{\mathrm{basetype}} + \xi n_{\mathrm{bptype}})(\nu + 1)\mu^2 + 2\mu + \nu + 1 \qquad \text{floating-block formulation} \qquad \text{(S10)}$$

A regressed parameter set comprises a vector of free energy parameters, $\vec{g}_{37}$, and a vector of enthalpy parameters, $\vec{h}$, which are used to define the temperature-dependent free energy parameter set, $\vec{g}$, using (S3). Each parameter in $\vec{g}$ is the negative logarithm of a non-zero scalar, vector element, or matrix element itemized in Table S1. Hence, upon negation and exponentiation of a parameter in $\vec{g}$, the resulting scalar, vector element, or matrix element is in $R_{>0}$.

Note that we ignore the presence/absence of 3′ terminal phosphates for the purposes of parameter regression.[25] If desired, future regressions could regress values for both a $z_{3'}$ (without 3′ terminal phosphate) and $z_{3'}^{\mathrm{phosphate}}$ (with 3′ terminal phosphate) to capture distinct stacking of the 3′ terminus on the final base or base pair with/without a 3′ terminal phosphate.

**Table S1:** Number of parameters for different model formulations.

| Term | Parameters | | |
|---|---|---|---|
| | Fixed Weights | Floating Weights | Floating Blocks |
| $e^{-g_{\text{assoc}}}$ | 1 | 1 | 1 |
| $e^{a_1}, \ldots, e^{a_\nu}$ | $\nu$ | — | — |
| $e^{s_1}, \ldots, e^{s_\nu}$ | — | $\nu$ | $\nu$ |
| $\bar{z}_{5'}$ | $\mu$ | $\mu$ | $\mu$ |
| $\bar{z}_{3'}$ | $\mu$ | $\mu$ | $\mu$ |
| $e^{-g_{\text{init}}}$ | 1 | — | — |
| $e^{b_0}, \ldots, e^{b_\nu}$ | $\nu + 1$ | — | — |
| $e^{b_i^{(0)}(\phi)}, \ldots, e^{b_i^{(\nu)}(\phi)}$ | — | $(\nu + 1)\, n_{\text{basetype}}$ | — |
| $e^{c_{i,j}^{(0)}(\phi)}, \ldots, e^{c_{i,j}^{(\nu)}(\phi)}$ | — | $(\nu + 1)\, n_{\text{bptype}}$ | — |
| $\bar{V}_i$ | $\mu^2\, n_{\text{basetype}}$ | $\mu^2\, n_{\text{basetype}}$ | — |
| $\bar{B}_{i,j}^{\rho}$ | $\mu^2 \xi\, n_{\text{bptype}}$ | $\mu^2 \xi\, n_{\text{bptype}}$ | — |
| $\tilde{V}_i$ | — | — | $(\nu + 1)\mu^2\, n_{\text{basetype}}$ |
| $\tilde{B}_{i,j}^{\rho}$ | — | — | $(\nu + 1)\mu^2 \xi\, n_{\text{bptype}}$ |

#### S2.8 Representative RNA and DNA parameter sets for transfer matrix models

NUPACK 4.2 supports the following temperature-dependent parameter sets for transfer matrix models:*

- `rna-matrix-26`: RNA parameters for transfer matrix model $(\nu, \mu, \xi) = 2, 2, 1$ with the floating-weight formulation in 1.0 M $Na^+$.
- `dna-matrix-26`: DNA parameters for transfer matrix model $(\nu, \mu, \xi) = 2, 2, 1$ with the floating-weight formulation in 1.0 M $Na^+$.
- `custom`: Custom parameters based on user-specified parameters representing nucleic acids or synthetic nucleic acid analogs in experimental conditions of choice.

To enable temperature-dependent calculations (see Section S2.3), parameter sets contain (dimensionless) free energy parameters, $\vec{g}_{37}$, at reference temperature $T_{37} \coloneqq 310.15$ K, and (dimensionless) enthalpy parameters, $\vec{h}$, assumed to be temperature independent.

*Will be made available in conjunction with publication in a peer-reviewed journal.

### S3 Calculation of diverse physical quantities

#### S3.1 Quantities defined over a complex ensemble

We provide dynamic programs to calculate the following physical quantities for a complex ensemble with sequence $\phi$:

- the partition function,

$$\overline{Q}(\phi) = \sum_{s \in \overline{\Gamma}(\phi)} e^{-\overline{\Delta G}(\phi,s)/kT}, \tag{S11}$$

  over ensemble $\overline{\Gamma}(\phi)$ treating all strands as distinct. The equilibrium probability of any secondary structure $s \in \overline{\Gamma}(\phi)$ is then

$$\overline{p}(\phi, s) = e^{-\overline{\Delta G}(\phi,s)/kT} / \overline{Q}(\phi). \tag{S12}$$

  Post-processing $\overline{Q}(\phi)$ yields the partition function $Q(\phi)$ over ensemble $\Gamma(\phi)$ treating strands of the same species as indistinguishable.[13, 19]

- the base-pairing probability matrix $\overline{P}(\phi)$ with entries $\overline{P}^{i,j}(\phi) \in [0, 1]$ corresponding to probability

$$\overline{P}^{i,j}(\phi) = \sum_{s \in \overline{\Gamma}(\phi)} \overline{p}(\phi, s) S^{i,j}(s) \tag{S13}$$

  that base pair $i \cdot j$ forms at equilibrium within ensemble $\overline{\Gamma}(\phi)$, treating all strands as distinct. Here, $S(s)$ is a *structure matrix* with entries $S^{i,j}(s) = 1$ if structure $s$ contains base pair $i \cdot j$ and $S^{i,j}(s) = 0$ otherwise. Abusing notation, the entry $S^{i,i}(s)$ is 1 if base $i$ is unpaired in structure $s$ and 0 otherwise; the entry $P^{i,i}(\phi) \in [0, 1]$ denotes the equilibrium probability that base $i$ is unpaired over ensemble $\overline{\Gamma}(\phi)$. Hence $S(s)$ and $\overline{P}(\phi)$ are symmetric matrices with row and column sums of 1.

- the free energy of the minimum free energy (MFE) subensemble state $s''_{\text{MFE}}(\phi) \in \overline{\Gamma}''(\phi)$ treating all strands as distinct:

$$\overline{\Delta G}(\phi, s''_{\text{MFE}}(\phi)) = \min_{s'' \in \overline{\Gamma}''(\phi)} \overline{\Delta G}(\phi, s''). \tag{S14}$$

- the MFE proxy structure

$$s_{\text{MFE}'} = \{s \in \overline{\Gamma}(\phi) \mid s''_{\text{MFE}}(s) \in s\}. \tag{S15}$$

  defined as the secondary structure containing the MFE subensemble state. The free energy of the MFE proxy structure is reported as

$$\Delta G(\phi, s_{\text{MFE}'}(\phi)), \tag{S16}$$

  treating strands of the same species as indistinguishable, which satisfies the inequality

$$\Delta G(\phi, s_{\text{MFE}'}(\phi)) \geq \Delta G(\phi) \tag{S17}$$

  relative to the complex free energy. If there is more than one MFE subensemble state, the algorithm returns all corresponding MFE proxy structures. Optionally, the structure search of equation (S15) may be performed over ensemble $\Gamma(\phi)$ instead of $\overline{\Gamma}(\phi)$, so that the MFE proxy structure is determined treating strands of the same species as indistinguishable.

- the set of suboptimal proxy structures

$$\overline{\Gamma}_{\text{subopt}}(\phi; \Delta G_{\text{gap}}) = \{s \in \overline{\Gamma}(\phi) \mid s'' \in s, \overline{\Delta G}(\phi, s'') \leq \overline{\Delta G}(\phi, s''_{\text{MFE}}(\phi)) + \Delta G_{\text{gap}}\} \tag{S18}$$

  with subensemble states within a specified $\Delta G_{\text{gap}} \geq 0$ of the MFE subensemble state. Optionally, the structure search of equation (S18) may be performed over ensemble $\Gamma(\phi)$ instead of $\overline{\Gamma}(\phi)$, so that the MFE proxy structure and the suboptimal proxy structures in the requested energy gap are determined treating strands of the same species as indistinguishable.

- a set of $J$ secondary structures Boltzmann-sampled from ensemble $\overline{\Gamma}(\phi)$ treating all strands as distinct:

$$\overline{\Gamma}_{\text{sample}}(\phi; J) \in \overline{\Gamma}(\phi). \tag{S19}$$

  If strands of the same species are treated as indistinguishable, the same sampled secondary structures may alternatively be interpreted as the set of $J$ secondary structures Boltzmann-sampled from ensemble $\Gamma(\phi)$:

$$\Gamma_{\text{sample}}(\phi; J) \in \Gamma(\phi). \tag{S20}$$

##### S3.2 Quantities defined over a test tube ensemble

We provide algorithms to calculate the following physical quantities for a test tube ensemble containing an arbitrary set of strand species $\Psi^0$ interacting to form an arbitrary set of complex species $\Psi$:

- the set of equilibrium concentrations $x_\Psi \equiv x_c \ \forall c \in \Psi$, (specified as mole fractions) that are the unique solution to the strictly convex optimization problem:[13]

$$\min_{x_\Psi} \sum_{c \in \Psi} x_c (\log x_c - \log Q_c - 1) \tag{S21a}$$

$$\text{subject to} \quad \sum_{c \in \Psi} A_{i,c} x_c = x_i^0 \quad \forall i \in \Psi^0, \tag{S21b}$$

  expressed in terms of the previously calculated set of partition functions $Q_\Psi$. Here, the constraints impose conservation of mass: $A$ is the stoichiometry matrix such that $A_{i,c}$ is the number of strands of type $i$ in complex $c$, and $x_i^0$ is the total concentration of strand $i$ present in the test tube. Based on dimensional analysis,[13] the convex optimization algorithm operates on mole fractions, but for convenience, accepts molar strand concentrations $[i]^0 = x_i^0 \rho_{H_2O}$ as inputs and returns molar complex concentrations $[c] = x_c \rho_{H_2O}$ as outputs, where $\rho_{H_2O}$ is the molarity of water.
- the ensemble pair fractions for the test tube ensemble, for example

$$f_A(i_A \cdot j_B) \tag{S22}$$

  denotes the fraction of A strands that form base pair $i_A \cdot j_B$ (correspondingly $f_B(i_A \cdot j_B)$ denotes the fraction of B strands that form base pair $i_A \cdot j_B$). In order to calculate these base-pairing observables, it is first necessary to calculate the set of equilibrium concentrations $x_\Psi$ and the set of base-pairing probability matrices $\overline{P}_\Psi$.

#### S4 Recursions incorporating the complex ensemble and free energy model

Recursions specify the dependencies between subproblems and incorporate the details of the structural ensemble and the free energy model. The recursions described here can be combined with a quantity-specific evaluation algebra (Section 3.3) and a quantity-specific operation order (Section S6) to calculate diverse physical quantities. Each recursion corresponds to an efficient iteration through a conditional ensemble of substructures within a given subsequence that are compatible with a specified set of constraints.

##### S4.1 Conventions for recursion diagrams, equations, and pseudocode in the main text

A recursion is represented in two ways: graphically, as a set of recursion diagrams, and algebraically, as an equation defining the recursion as a specific combination of contributions. The recursion diagrams employ the following conventions:

- Solid circular arcs depict the nucleic acid backbone. An arrowhead denotes the $3'$ end of a strand.
- Dots indicate particular nucleotide positions that define the bounds of recursive contributions. If a dot is labeled with a nucleotide index, the same index is used in the corresponding recursion. If a dot is adjacent to a dot labeled $i$, the implied index of the unlabeled dot is either $i-1$ or $i+1$ (indices increase from $5'$ to $3'$).
- A straight line delimits the boundary for a given contribution. A solid straight line indicates that the connected nucleotides are base-paired. A dashed straight line indicates that the connected nucleotides may or may not be base-paired.
- Shading indicates that the shaded region in a recursion explicitly incorporates a *recursion energy* using a transfer matrix or tensor (green shading for $V$, blue shading for $B$).

A recursion equation provides a mathematical description of the conditional ensemble depicted graphically in a recursion diagram. Recursion equations employ the following conventions:

- For each physical quantity, an appropriate evaluation algebra (Section 3.3) is used to define the generic operators that appear in the recursion equations: $\mathbb{0}$, $\mathbb{1}$, $\oplus$, $\otimes$, $W$, and $Q$. For example, to calculate the partition function, we have:

$$\mathbb{0} \to 0, \quad \mathbb{1} \to 1, \quad \oplus \to +, \quad \otimes \to \times, \quad W(g) \to e^{-g}, \quad Q(r,i,j) \to Q^{r}_{i,j}. \tag{S23}$$

  where the last right-hand side indicates that $Q^r_{i,j}$ is a lookup of the relevant stored matrix element. Note that element $Q^r_{i,j}$ is a $\gamma \times \gamma$ matrix with entries $Q^{r;a,b}_{i,j}$ for $a \in [\gamma], b \in [\gamma]$.
- A recursion equation for subsequence $[i,j]$ corresponding to element $i,j$ in a triangular block is denoted $Q(r,i,j)$ for a recursion of type $r \in \{\varnothing, e\}$. Here, $\phi$ is the sequence of the complex and $i$ and $j$ are base indices. Note that in the Supporting Information we use $Q^{\varnothing}_{i,j}$ to denote $Q_{i,j}$ so that each recursion has an explicit recursion type $r$.
- If a recursion diagram contains a shaded region denoting a recursion energy, $g$, the corresponding recursion equation will incorporate the recursion energy via a term $W(g)$ used to define a $V$ or $B$ matrix entry (3.4).
- After it is evaluated for the first time, $Q(r,i,j)$ is used to yield $Q^r_{i,j}$ in subsequent recursions. In the evaluation algebras that generate scalars (SumProduct, MinSum, Count), the output of $Q^r(i,j)$ is synonymous with the value $Q^r_{i,j}$ that is stored in the recursion matrices. However, other evaluation algebras involve different treatment of the output of $Q(r,i,j)$. The ways in which recursion outputs are utilized for each physical quantity are described in Section S6.

##### S4.2 Optimized $O(N^3)$ algorithm with reduced time-complexity prefactor

Algorithm 3.2 calculates the complex partition function, $Q(\phi)$, using a single recursion displayed in the recursion diagram of Figure 7b and recursion equation (3.1). The time complexity of Algorithm 3.2, evident from the right hand side of the recursion diagram and equation, is $O(\xi\gamma^3 N^3)$, where $N^3$ comes from iterating over base indices $i, j, k$, and the prefactor contains $\gamma^3$ due to multiplying $\gamma \times \gamma$ matrices and $\xi$ due to iterating over base pair subtype index $\rho$. The space complexity of Algorithm 3.2, evident from the left hand side of the recursion diagram and equation, is $O(\gamma^2 N^2)$, where $N^2$ comes from storing recursion element, $Q^{\varnothing}_{i,j}$, for base indices $i, j$, and the prefactor, $\gamma^2$, comes from $Q^{\varnothing}_{i,j}$ being a $\gamma \times \gamma$ matrix.

Here, we show that by using two recursions instead of one, it is possible to reduce the time complexity to $O(\gamma^3 N^3 + \xi\gamma^2 N^2)$, which has the notable property that for large $N$, the complexity is asymptotically independent of $\xi$. The optimized algorithm with lower time complexity is described in the pseudocode of Algorithm S1, which uses the same operation order as Algorithm 3.2 (see Figure S1a) , adding a new $3'$-most base $j$ at each iteration. The new recursion diagrams are depicted in Figure S1b and the corresponding recursion equations are displayed in Figure S1c.

**$Q^e$ recursion: the "extend" operation.** The $Q^e_{i,j}$ recursion performs an *extend* operation by extending the subsequence $[i, j]$ to have a new $3'$-most base $j$. Considering each possible value for $i$, the recursion encompasses two scenarios:

- $i = j$ and the new base $j$ is unpaired: the transfer matrix $V_j$ is incorporated.
- $i < j$ and the new base $j$ is base-paired to base $i$: the tensors $B^\rho_{j,i}$ and $B^\rho_{i,j}$ are incorporated into the neighboring loops on either side of base pair $i \cdot j$, making use of the previously computed subsequence partition function $Q^\varnothing_{i+1,j-1}$. On one side of the base pair $i \cdot j$, tensor $B^\rho_{j,i}$ left multiplies $Q^\varnothing_{i+1,j-1}$ and closes a loop, with $\mathrm{Tr}_S$ contracting the final edge in the loop to yield a scalar, and on the other side of base pair $i \cdot j$, $B^\rho_{j,i}$ contributes partition function information for a new loop that will be closed in another recursion. Summation is performed over each alternative base pair subtype $\rho \in \xi$ for each alternative base $i < j$.

The time complexity of the $Q^e$ recursion is $O(\xi\gamma^2 N^2)$, where $N^2$ comes from iterating over base indices $i, j$, and the prefactor contains $\gamma^2$ due to taking the trace of a product of $\gamma \times \gamma$ matrices and $\xi$ due to iterating over base pair subtype index $\rho$. Note that taking the product of $\gamma \times \gamma$ matrices has $\gamma^3$ time complexity, but taking the standard trace, Tr, reduces this complexity to $\gamma^2$ (since only the diagonal entries of the product need be computed), and taking the customized trace, $\mathrm{Tr}_S$, retains this $\gamma^2$ complexity due to the sparsity of $S$ (2.40).

**$Q^\varnothing$ recursion: the "combine" operation.** The $Q^\varnothing_{i,j}$ recursion represents the partition function for subsequence $[i, j]$ with new $3'$-most base $j$, performing the *combine* operation by combining the previously computed subsequence partition function $Q^\varnothing_{i,k-1}$ with the newly computed extension $Q^e_{k,j}$ for any base $k$ in the interval. The time complexity of the $Q^\varnothing$ recursion is $O(\gamma^3 N^3)$, where $N^3$ comes from iterating over base indices $i, j, k$, and the prefactor contains $\gamma^3$ due to multiplying $\gamma \times \gamma$ matrices.

**Explicit model dependence.** Note that the only explicit dependence on the free energy model (incorporated using $V$ transfer matrices and $B^\rho$ tensors) occurs in the $O(N^2)$ extend operation of the $Q^e$ recursion, and that the $Q^\varnothing$ recursion, which is the sole $O(N^3)$ component of the algorithm, contains no explicit dependence on the free energy model (no $V$ or $B^\rho$ terms), and so is particularly amenable to GPU acceleration using linear algebra primitives.

**Implicit treatment of nicks.** The $Q^e$ and $Q^\varnothing$ recursions implicitly incorporate the contributions of a nick in an exterior loop via the definitions of $V_i$ and $B^\rho_{i,j}$ using Eq. (2.13), which conditionally include a $Z$ matrix if there is a nick before $i$. Furthermore, the interplay of the sparsity patterns of the $S$ and $Z$ matrices automatically exclude contributions from loops containing more than one nick, which would correspond to disconnected structures that are not in the complex ensemble.

**Algorithm S1** Optimized $O(N^3)$ algorithm to calculate the complex partition function $Q(\phi)$ with reduced time complexity prefactor.

```
function ComplexPartitionFunction(N, V, B)
    for j ∈ [N] do
        Q^∅_{j,j−1} ← I                                  ▷ Base case for empty subsequence [j, j)
        for all i ∈ [j − 1] do
            Q^e_{i,j} ← Σ_{ρ∈ξ} B^ρ_{i,j} Tr_S(B^ρ_{j,i} Q^∅_{i+1,j−1})   ▷ Extend subsequence to j with base pair i · j
        Q^e_{j,j} ← V_j                                  ▷ Extend subsequence to j with unpaired base j
        for all i ∈ [j] do
            Q^∅_{i,j} ← Σ_{k∈[i:j]} Q^∅_{i,k−1} Q^e_{k,j}   ▷ Combine previously computed Q^∅ with newly computed Q^e
    return Tr_S(Q^∅_{1,N})
```

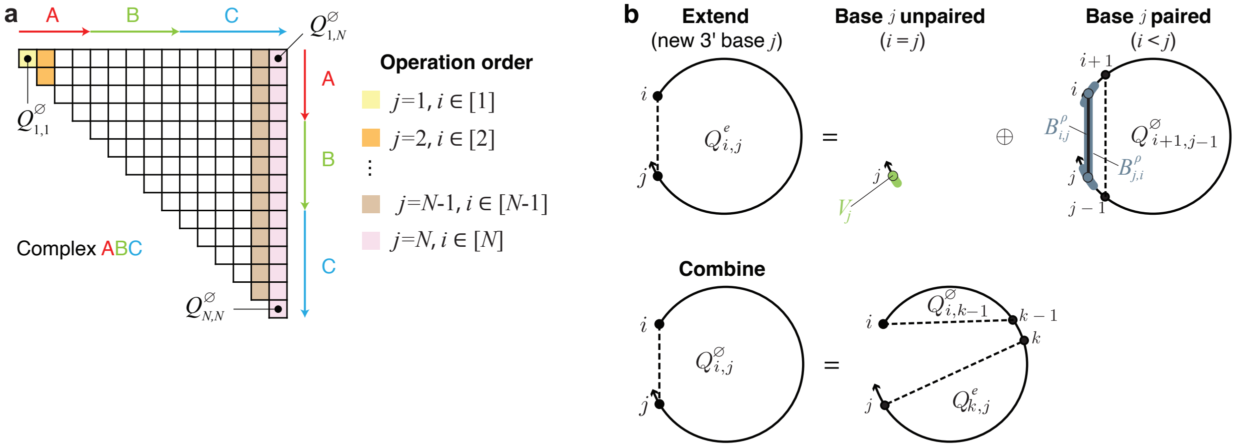

$$Q(e,i,j) \coloneqq \begin{cases} V_j, & i = j \\ \bigoplus_{\rho \in [\xi]} B^{\rho}_{i,j} \otimes \mathrm{Tr}_S(B^{\rho}_{j,i} \otimes Q^{\varnothing}_{i+1,j-1}), & i < j \end{cases}$$

$$Q(\varnothing,i,j) \coloneqq \bigoplus_{k \in [i:j]} Q^{\varnothing}_{i,k-1} \otimes Q^{e}_{k,j}$$

**Figure S1:** A dynamic programming algorithm for calculating the partition function of a complex based on the new tensor model using a single recursion (see the pseudocode of Algorithm 3.2). **(a)** Operation order. A forward sweep of 3′-most base $j$ from 1 to $N$, considering all subsequences $[i, j]$ for each $j$. **(b)** Recursion diagrams for $Q^{e}_{i,j}$ and $Q_{i,j}$. $Q^{e}_{i,j}$ represents the partition function contributions for subsequence $[i, j]$ where new 3′-most base $j$ is unpaired (if $i = j$) or is paired to base $i$ (if $i < j$). $Q^{\varnothing}_{i,j}$ respresents the partition function contributions for subsequence $[i, j]$. The $Q^{e}_{i,j}$ recursion performs the *extend* operation by extending the subsequence to a new 3′-most base $j$, and the $Q^{\varnothing}_{i,j}$ performs the *combine* operation by combining a newly computed $Q^{e}_{k,j}$ contribution with a previously computed $Q^{\varnothing}_{i,k-1}$ contribution. A solid straight line indicates a base pair and a dashed line demarcates a region without implying that the connected bases are paired. Shading corresponds to a transfer matrix ($V$) or tensor ($B$) incorporated at the current level of recursion. **(c)** Recursion equations for $Q^{e}_{i,j}$ and $Q^{\varnothing}_{i,j}$.

#### S5 Evaluation algebras for calculating each physical quantity

To calculate each physical quantity, the generic recursions of Section S4 are combined with a quantity-specific *evaluation algebra* (summarized in Table 3). Here, we provide additional details on the evaluation algebra abstraction, and on the definition of evaluation algebras for different physical quantities.

An evaluation algebra defines the mathematical form of the generic operators that appear in recursion equations. We define an evaluation algebra $\mathcal{A}$ that operates on transfer matrices with entries $(a \in [\gamma], b \in [\gamma])$ to have the following properties:

1. $\mathcal{D}$ is the domain of values in the evaluation algebra for each matrix entry $(a, b)$.
2. $\oplus$ is a matrix operation yielding a combination of alternative conditional ensembles for each matrix entry $(a, b)$. Thus, $U = X \oplus Y$ reflects the notion that "$U^{a,b}$ contains either conditional ensemble $X^{a,b}$ or conditional ensemble $Y^{a,b}$".
3. $\otimes$ is a matrix operation that yields a composition of conditional ensembles for each matrix entry $(a, b)$. Thus, $U = X \otimes Y$ reflects the notion that "$U^{a,b}$ contains both conditional ensemble $X^{a,c}$ and conditional ensemble $Y^{c,b}$ for some $c \in [\gamma]$".
4. $\mathbb{0}$ is a value in $\mathcal{D}$ that satisfies the concept of additive identity. Physically, $\mathbb{0}$ represents an impossible substructure (i.e., a structural element that is not in the complex ensemble).
5. $\mathbb{1}$ is a value in $\mathcal{D}$ that satisfies the concept of multiplicative identity. Physically, $\mathbb{1}$ represents a substructure with a free energy contribution of 0.
6. $W(g)$ is an operation that takes a (dimensionless) free energy $g$ to a value in $\mathcal{D}$. Physically, $W$ represents the weight on an individual substructure.
7. $Q(\lambda)$ is an operation that yields a value in $\mathcal{D}$ from a recursion element $\lambda$ in the set of all recursion elements $\Lambda$. $Q$ is used to give the previously-calculated result over a given conditional ensemble.

An evaluation algebra may be classified as an algebraic semiring equipped with two additional unary operators $W$ and $Q$. $\mathbb{I}$ is a matrix of dimension $\gamma$ with $\mathbb{1}$ on the diagonal and $\mathbb{0}$ off the diagonal. The weighted trace takes the form

$$\mathrm{Tr}_S(X) = \bigoplus_a (S \otimes X)^{a,a}. \tag{S24}$$

We now describe the definitions of these properties for evaluation algebras corresponding to different physical quantities, treating the calculation of scalar physical quantities in Section S5.1 and the calculation of quantities requiring structure generation in Section S5.2.

##### S5.1 Evaluation algebras for scalar outputs

###### S5.1.1 SumProduct: partition function

Within evaluation algebra $\mathcal{A}$ = SumProduct, the partition function of an ensemble, $\overline{Q}(\phi)$, is computed by taking the sum over products of Boltzmann factors.

$$\begin{aligned}
\mathcal{D} &= \mathbb{R}_{\geq 0} \\
X \oplus Y &= X^{a,b} + Y^{a,b}, \quad a, b \in [\gamma] \\
X \otimes Y &= \textstyle\sum_{c \in [\gamma]} X^{a,c} \cdot Y^{c,b}, \quad a, b \in [\gamma] \\
\mathbb{0} &= 0 \\
\mathbb{1} &= 1 \\
W(g) &= e^{-g} \\
Q(\lambda) &= Q^{r,a,b}_{i,j} \text{where } \lambda = (r, i, j, a, b) \text{ and } Q^r_{i,j} \text{ is the stored recursion matrix of type } r \in \{\varnothing, e\}.
\end{aligned} \tag{S25}$$

Each matrix entry in the algebra represents a Boltzmann factor, which is necessarily a non-negative real number. An impossible structure maps to a Boltzmann factor of 0, whereas a structure with a free energy of 0 maps to a Boltzmann factor of 1.

###### S5.1.2 COUNT: structure count

Within evaluation algebra $\mathcal{A}$ = COUNT, the size of the ensemble, $\overline{\Gamma}^{\shortparallel}(\phi)$, is computed by taking the sum over products of subensemble sizes.

$$\begin{aligned}
\mathcal{D} &= \mathbb{Z}_{\geq 0} \\
X \oplus Y &= X^{a,b} + Y^{a,b}, \quad a, b \in [\gamma] \\
X \otimes Y &= \textstyle\sum_{c \in [\gamma]} X^{a,c} \cdot Y^{c,b}, \quad a, b \in [\gamma] \\
\mathbb{0} &= 0 \\
\mathbb{1} &= 1 \\
W(g) &= 1 \\
Q(\lambda) &= Q_{i,j}^{r,a,b} \text{where } \lambda = (r, i, j, a, b) \text{ and } Q_{i,j}^{r} \text{ is the stored recursion matrix of type } r \in \{\varnothing, e\}.
\end{aligned} \tag{S26}$$

The only difference between COUNT and SUMPRODUCT is the definition of $W$. While the domain of COUNT is theoretically non-negative integers, it is still implemented using floating point types to avoid integer overflow.

###### S5.1.3 MINSUM: free energy of MFE subensemble state

Within evaluation algebra $\mathcal{A}$ = MINSUM, the free energy of the minimum free energy (MFE) subensemble state, $\overline{\Delta G}(\phi, s^{\shortparallel}_{\text{MFE}})$, is the minimum over sums of conditional ensemble free energies.

$$\begin{aligned}
\mathcal{D} &= \mathbb{R} \cup \{\infty\} \\
X \oplus Y &= \min(X^{a,b}, Y^{a,b}), \quad a, b \in [\gamma] \\
X \otimes Y &= \min_{c \in [\gamma]}(X^{a,c} + Y^{c,b}), \quad a, b \in [\gamma] \\
\mathbb{0} &= \infty \\
\mathbb{1} &= 0 \\
W(g) &= g \\
Q(\lambda) &= Q_{i,j}^{r,a,b} \text{where } \lambda = (r, i, j, a, b) \text{ and } Q_{i,j}^{r} \text{ is the stored recursion matrix of type } r \in \{\varnothing, e\}.
\end{aligned} \tag{S27}$$

An impossible structure that is not in the complex ensemble is assigned a free energy of $\infty$. We treat the primitives of MFE computation as $S$ (2.40), $Z$ (2.13), $\tilde{V}$ and $\tilde{B}$ [equation (2.41) or (2.42)], with operations between these matrices interpreted using the MINSUM evaluation algebra.

###### S5.1.4 $\partial$SUMPRODUCT: gradient of partition function

Consider evaluation of the gradient of the partition function with respect to a parameter

$$\frac{\partial \overline{Q}(\phi)}{\partial \theta} \tag{S28}$$

and note that the partition function can be used to calculate the complex free energy

$$\overline{\Delta G}(\phi) = \overline{\Delta H}(\phi) - T\overline{\Delta S}(\phi) = -\beta \log \overline{Q}(\phi), \tag{S29}$$

where $\beta \coloneqq k_B T$. Evaluation of the gradient of the partition function with respect to $\beta$ then enables calculation of the complex enthalpy

$$\overline{\Delta H}(\phi) = \frac{\partial \beta \overline{\Delta G}(\phi)}{\partial \beta} = \frac{\partial \overline{g}(\phi)}{\partial \beta}, \tag{S30}$$

and the complex entropy (e.g., for calculation of loop entropic cost for Figure 10)

$$\overline{\Delta S}(\phi) = -\frac{\partial \overline{\Delta G}(\phi)}{\partial T} = k_B \beta^2 \frac{\partial \beta^{-1} \overline{g}(\phi)}{\partial \beta}. \tag{S31}$$

More generally, evaluation of the gradient of the partition function with respect to a parameter enables calculation of the derivative of any equilibrium observable with respect to an input parameter $\theta$ (e.g., see the parameter regression of Section S9 without the use of finite differences).

The $\partial$SumProduct evaluation algebra uses automatic differentiation with the conventional rules of differentiation. Here, each element in the algebra is a pair of 1) the value of the quantity (subscript v) in the SumProduct evaluation algebra and 2) the derivative (subscript d) of that quantity with respect to a specified parameter $\theta$:

$$\begin{aligned}
\mathcal{D} &= \mathbb{R}_{\geq 0} \times \mathbb{R} \\
X \oplus Y &= \left(X_{\mathrm{v}}^{a,b} + Y_{\mathrm{v}}^{a,b},\ X_{\mathrm{d}}^{a,b} + Y_{\mathrm{d}}^{a,b}\right), \quad a,b \in [\gamma] \\
X \otimes Y &= \left(\textstyle\sum_{c\in[\gamma]} X_{\mathrm{v}}^{a,c} \cdot Y_{\mathrm{v}}^{c,b},\ \textstyle\sum_{c\in[\gamma]} X_{\mathrm{v}}^{a,c} \cdot Y_{\mathrm{d}}^{c,b} + X_{\mathrm{d}}^{a,c} \cdot Y_{\mathrm{v}}^{c,b}\right), \quad a,b \in [\gamma] \\
\mathbb{0} &= (0,0) \\
\mathbb{1} &= (1,0) \\
W(g) &= \left(e^{-g}, \frac{\partial e^{-g}}{\partial \theta}\right) \\
Q(\lambda) &= Q_{i,j}^{r,a,b} \text{where } \lambda = (r,i,j,a,b) \text{ and } Q_{i,j}^{r} \text{ is the stored recursion matrix of type } r \in \{\varnothing, e\}.
\end{aligned} \tag{S32}$$

#### S5.2 Evaluation algebras for structure generation

Structure generation conceptually yields specific secondary structures from a given weighting on the ensemble $\overline{\Gamma}^{\shortparallel}(\phi)$. In this case, because any given structure depends on only a sparse subset of recursion matrix elements, a backtracking operation order is in general more efficient than a forward pass iteration. Such an operation order jumps between recursion elements in an opposite direction to that in the forward pass. To enable such an approach, the recursion matrices in the forward pass must be computed beforehand (e.g., calculating the MFE before generating an ensemble of suboptimal structures or calculating the partition function before Boltzmann sampling structures).

Here, we correspondingly distinguish between a *forward* evaluation algebra and a *backtracking* evaluation algebra. Whereas a forward evaluation algebra like SumProduct operates on a subset of $\mathbb{R}$, we define a backtracking evaluation algebra to operate on a domain of conditional ensembles which may be queried for a set of dependent recursion elements. We next illustrate how a backtracking evaluation algebra may be defined with respect to the associated forward evaluation algebra.

##### S5.2.1 Generic approach to structure generation

We start with a consideration of the simplest structure generation evaluation algebras. To simplify the exposition, in Table 3 we defined evaluation algebras ArgRand to calculate a single randomly sampled structure and ArgMin to determine the MFE stacking state, $s^{\shortparallel}_{\mathrm{MFE}}$, assuming it was unique. For a given element $X^{a,b}$, each evaluation algebra was defined such that $X^{a,b}$ was a pair of scalar value $X_{\mathrm{v}}^{a,b}$ and recursion element set $X_{\lambda}^{a,b}$.

In Table 3, the operations on $X_{\mathrm{v}}$ and $Y_{\mathrm{v}}$ in ArgRand duplicate the operations of SumProduct, whereas the operations on $X_{\mathrm{v}}$ and $Y_{\mathrm{v}}$ in ArgMin duplicate the operations of MinSum. In ArgRand, $\oplus$ yields a random weighted choice via

$$\arg\,\mathrm{rand}(X,Y) \equiv \left(X_{\lambda}^{a,b} \text{ if } \mathcal{U}(0, X_{\mathrm{v}}^{a,b} + Y_{\mathrm{v}}^{a,b}) < X_{\mathrm{v}}^{a,b} \text{ else } Y_{\lambda}^{a,b}\right), \quad a,b \in [\gamma]. \tag{S33}$$

where $\mathcal{U}$ is the random uniform distribution function. In contrast, in ArgMin, $\oplus$ yields the conditional ensemble which is lower in free energy via

$$\arg\,\min(X,Y) \equiv \left(X_{\lambda}^{a,b} \text{ if } X_{\mathrm{v}}^{a,b} < Y_{\mathrm{v}}^{a,b} \text{ else } Y_{\lambda}^{a,b}\right), \quad a,b \in [\gamma]. \tag{S34}$$

Thus we can see that an intuitive approach for constructing a backtracking evaluation algebra is to augment a corresponding forward evaluation algebra with customized operations for structure attribution. We next describe the resulting definitions of ArgRand and ArgMin.

##### S5.2.2 ArgRand: Boltzmann sample single structure

Within evaluation algebra $\mathscr{A}$ = ArgRand, each element $X$ is a pair of partition function values $X_\mathrm{v}$ and sets of recursion elements $X_\lambda$.

$$\begin{aligned}
\mathcal{D} &= \mathbb{R}_{\geq 0} \times \mathcal{P}(\Lambda) \\
X \oplus Y &= \left(X_\mathrm{v}^{a,b} + Y_\mathrm{v}^{a,b}, \arg\ \mathrm{rand}(X_\mathrm{v}^{a,b}, Y_\mathrm{v}^{a,b})\right), \quad a, b \in [\gamma] \\
X \otimes Y &= \left(\sum_{c\in[\gamma]} X_\mathrm{v}^{a,c} Y_\mathrm{v}^{c,b}, \arg\ \mathrm{rand}_{c\in[\gamma]}(X_\mathrm{v}^{a,c} Y_\mathrm{v}^{c,b})\right), \quad a, b \in [\gamma] \\
\mathbb{0} &= (0, \varnothing) \\
\mathbb{1} &= (1, \varnothing) \\
W(g) &= (e^{-g}, \varnothing) \\
Q(\lambda) &= (Q_{\textsc{SumProduct}}(\lambda), \{\lambda\})
\end{aligned} \tag{S35}$$

Operations on the first element, $X_\mathrm{v}$, are defined using SumProduct. For each $a, b \in [\gamma]$, the second element, $X_\lambda^{a,b}$, is a set of recursion elements defining a restricted ensemble of conditional ensemble compositions. The set of all possible sets of recursion elements $\lambda$ is denoted as $\mathcal{P}(\Lambda)$. Any quantity that does not depend on the output of another recursion is thus assigned $X_\lambda^{a,b} = \varnothing$.

##### S5.2.3 ArgMin: MFE structure determination

Within evaluation algebra $\mathscr{A}$ = ArgMin, each element $X$ is a pair of free energy values $X_\mathrm{v}$ and set of recursion elements $X_\lambda$.

$$\begin{aligned}
\mathcal{D} &= \mathbb{R} \cup \{\infty\} \times \mathcal{P}(\Lambda) \\
X \oplus Y &= \left(\min(X_\mathrm{v}^{a,b}, Y_\mathrm{v}^{a,b}), \arg\ \min(X_\mathrm{v}^{a,b}, Y_\mathrm{v}^{a,b})\right) \quad a, b \in [\gamma] \\
X \otimes Y &= \left(\min_{c\in[\gamma]} X_\mathrm{v}^{a,c} + Y_\mathrm{v}^{c,b}, \arg\ \min_{c\in[\gamma]}(X_\mathrm{v}^{a,c} + Y_\mathrm{v}^{c,b})\right) \quad a, b \in [\gamma] \\
\mathbb{0} &= (\infty, \varnothing) \\
\mathbb{1} &= (0, \varnothing) \\
W(g) &= (g, \varnothing) \\
Q(\lambda) &= (Q_{\textsc{MinSum}}(\lambda), \{\lambda\})
\end{aligned} \tag{S36}$$

Operations on the first element, $X_\mathrm{v}$, are defined using MinSum. For each $a, b \in [\gamma]$, the second element, $X_\lambda^{a,b}$, is a set of recursion elements defining a restricted ensemble of conditional ensemble compositions. The set of all possible sets of recursion elements $\lambda$ is denoted as $\mathcal{P}(\Lambda)$. Any quantity that does not depend on the output of another recursion is thus assigned $X_\lambda^{a,b} = \varnothing$.

##### S5.2.4 Efficient structure generation via lazy evaluation

We derived more programmatically efficient evaluation algebras for generating an ensemble of $J$ Boltzmann-sampled structures, $\overline{\Gamma}_\mathrm{sample}(\phi; J)$, or for generating a ensemble of suboptimal structures, $\overline{\Gamma}_\mathrm{subopt}(\phi; \Delta G_\mathrm{gap})$. Note that MFE proxy structure(s) can be generated by setting $\Delta G_\mathrm{gap} = 0$. Here, we describe efficient evaluation algebras for ArgRandJ and ArgMinGap that build upon ArgRand and ArgMin.

The simpler but less efficient algebras ArgRand and ArgMin yield full representations of the chosen conditional ensembles, which are then enqueued by the respective operation order algorithms. Our more efficient algorithms work by backtracking through a given recursion element and enqueueing any combinations of recursion elements in conditional ensembles that match a given criterion. The matching evaluation algebras incorporate the enqueueing operation $\kappa$ directly such that each piece of a conditional ensemble is enqueued immediately as it is encountered. These evaluation algebras are similarly generic but operate lazily on recursion elements (obviating storage of intermediate structures which might not affect the final results).

We describe the ArgRandJ and ArgMinGap matrix evaluation algebras using a generic framework defined with respect to two corresponding *scalar* forward and backtracking evaluation algebras. We make use of the anonymous form of function notation $x \mapsto y$ to notate a function taking $x$ and returning $y$. Formally, we define the enqueueing operation $\kappa$ recursively as a function in $D_\kappa \equiv (\mathbb{R}, \mathcal{P}(\Lambda)) \mapsto D_\kappa$; effectively, it may be viewed as an iterator across each alternative conditional ensemble. Let $\mathcal{M}$ be the matrix backtracking evaluation algebra, $\mathcal{R}$ be the scalar backtracking evaluation algebra and $\mathcal{F}$ be the scalar forward evaluation algebra. Then we classify each expression in $\mathcal{R}$ as a closure within $D_\mathcal{R} \equiv D_\kappa \mapsto D_\kappa$ and denote a set of recursion elements as $\Lambda_i \in \mathcal{P}(\Lambda)$ below.

Within the scalar backtracking evaluation algebra $\mathcal{R}$, addition of $x$ and $y$ intuitively represents the successive iteration through multiple alternative structures $x$ and $y$. It may be defined formally as a higher-order function that accomplishes functional composition:

$$\oplus_\mathcal{R}(x, y) = \kappa \mapsto y(x(\kappa)). \tag{S37}$$

Matricization then yields the matrix addition operator in $\mathcal{M}$:

$$\oplus_\mathcal{M}(X, Y) = X^{a,b} \oplus_\mathcal{R} Y^{a,b}, \quad a, b \in [\gamma] \tag{S38}$$

Multiplication of $X$ and $Y$ represents independent combinations of conditional ensembles from $X$ and $Y$ being composed together – in effect, a lazily evaluated outer product of conditional ensembles within $X$ and $Y$. For scalars, it may be defined formally as the higher-order function:

$$\otimes_\mathcal{R}(x, y) = \kappa \mapsto x\big((z_1, \Lambda_1) \mapsto y\big((z_2, \Lambda_2) \mapsto \kappa\big(z_1 \otimes_\mathcal{F} z_2, \Lambda_1 \cup \Lambda_2\big)\big)\big). \tag{S39}$$

Matricization and application of the standard matrix multiplication rules then yield:

$$\otimes_\mathcal{M}(X, Y) = \bigoplus_{c \in [\gamma]}{}_\mathcal{R}\, X^{a,c} \otimes_\mathcal{R} Y^{c,b}. \tag{S40}$$

The properties of commutativity and associativity are preserved for $\oplus_\mathcal{R}$ and $\otimes_\mathcal{R}$ so long as $\kappa$ is independent of the order of evaluation (i.e., $\kappa(z_1, \Lambda_1)(z_2, \Lambda_2) = \kappa(z_2, \Lambda_2)(z_1, \Lambda_1)$), a property that is satisfied by all algorithms discussed here. The multiplicative identity corresponds to a zero free energy structure, which is not dependent on any recursion elements:

$$\mathbf{1}_\mathcal{M} = \kappa \mapsto \kappa(\mathbf{1}_\mathcal{F}, \varnothing). \tag{S41}$$

The additive identity is defined as the identity function, reflecting an impossible structure by returning the enqueueing function unchanged:

$$\mathbf{0}_\mathcal{M} = \kappa \mapsto \kappa. \tag{S42}$$

$W_\mathcal{M}$ brings a free energy parameter into the evaluation algebra domain, and is not dependent on any recursion elements:

$$W_\mathcal{M}(g) = \kappa \mapsto \kappa\left(W_\mathcal{F}(g), \varnothing\right). \tag{S43}$$

Finally, the recursion matrix operator yields the forward algebra value and a singleton of its associated recursion element:

$$Q_\mathcal{M}(\lambda) = \kappa \mapsto \kappa\left(Q_\mathcal{F}(\lambda), \{\lambda\}\right). \tag{S44}$$

Boltzmann sampling and suboptimal structure generation can then proceed by specification of the enqueueing function $\kappa$, which registers new recursion elements to backtrack, and application of the respective operation orders in Sections S6.6 and S6.7.

##### S5.2.5 ArgRandJ: simultaneously Boltzmann sample J structures

The implemented evaluation algebra $\mathcal{A}$ = ArgRandJ that uses higher order functions to accomplish lazy evaluation is a specialization of the generic structure generation algebra (Section S5.2.4) for the associated forward evaluation algebra SumProduct.

$$
\begin{aligned}
\mathcal{D} &= \mathcal{D}_\kappa \mapsto \mathcal{D}_\kappa \\
X \oplus Y &= X^{a,b} \oplus_{\mathcal{R}} Y^{a,b} \text{ where } x \oplus_{\mathcal{R}} y = \kappa \mapsto y(x(\kappa)), \quad a,b \in [\gamma] \\
X \otimes Y &= \bigoplus_{c\in[\gamma]}{}_{\mathcal{R}}\, X^{a,c} \otimes_{\mathcal{R}} Y^{c,b} \text{ where } x \otimes_{\mathcal{R}} y = \kappa \mapsto x\big((z_1,\Lambda_1) \mapsto y\big((z_2,\Lambda_2) \mapsto \kappa\big(z_1 \cdot z_2, \Lambda_1 \cup \Lambda_2\big)\big)\big), \quad a,b \in [\gamma] \\
\mathbb{0} &= \kappa \mapsto \kappa \\
\mathbb{1} &= \kappa \mapsto \kappa(1,\varnothing) \\
W(g) &= \kappa \mapsto \kappa\,(e^{-g},\varnothing) \\
Q(\lambda) &= \kappa \mapsto \kappa(Q_{\textsc{SumProduct}}(\lambda),\{\lambda\})
\end{aligned}
\tag{S45}
$$

##### S5.2.6 ArgMinGap: suboptimal structure generation

The implemented evaluation algebra $\mathcal{A}$ = ArgMinGap that uses higher order functions to accomplish lazy evaluation is a specialization of the generic structure generation algebra (Section S5.2.4) for the associated forward evaluation algebra MinSum.

$$
\begin{aligned}
\mathcal{D} &= \mathcal{D}_\kappa \mapsto \mathcal{D}_\kappa \\
X \oplus Y &= X^{a,b} \oplus_{\mathcal{R}} Y^{a,b} \text{ where } x \oplus_{\mathcal{R}} y = \kappa \mapsto y(x(\kappa)), \quad a,b \in [\gamma] \\
X \otimes Y &= \bigoplus_{c\in[\gamma]}{}_{\mathcal{R}}\, X^{a,c} \otimes_{\mathcal{R}} Y^{c,b} \text{ where } x \otimes_{\mathcal{R}} y = \kappa \mapsto x\big((z_1,\Lambda_1) \mapsto y\big((z_2,\Lambda_2) \mapsto \kappa\big(z_1 + z_2, \Lambda_1 \cup \Lambda_2\big)\big)\big), \quad a,b \in [\gamma] \\
\mathbb{0} &= \kappa \mapsto \kappa \\
\mathbb{1} &= \kappa \mapsto \kappa(0,\varnothing) \\
W(g) &= \kappa \mapsto \kappa(g,\varnothing) \\
Q(\lambda) &= \kappa \mapsto \kappa(Q_{\textsc{MinSum}}(\lambda),\{\lambda\})
\end{aligned}
\tag{S46}
$$

### S6 Operation orders for calculating each physical quantity

#### S6.1 Notation conventions for production code and pseudocode of Section S6

Conventions for presentation of models and algorithms in the main text are described in Section S4.1. For production code and the pseudocode of Section S6, we make the following modifications to these conventions.

**Summary of main text conventions: 1-indexed triangular recursion matrices $Q^e$ and $Q^\varnothing$.** With reference to Section S4, using main text conventions:

- **1-indexing.** The nucleotides in a complex of $N$ nucleotides are indexed $j \in [N]$ starting at $j = 1$.
- **Recursion elements for subsequence $[i, j]$.** Subsequence $[i, j]$ is represented by recursion elements $Q^e_{i,j}$ and $Q^\varnothing_{i,j}$, each a $\gamma \times \gamma$ matrix.
- **1-indexed triangular recursion matrices.** The triangular matrix $Q^e$ has columns $j \in [N]$ and rows $i \in [j]$; the triangular matrix $Q^\varnothing$ has columns $j \in [N]$ and rows $i \in [j]$ augmented by the base case $Q^\varnothing_{j,j-1}$ corresponding to empty subsequence $[j, j-1]$.

**Summary of production code conventions: shifted triangular recursion matrices $\hat{Q}^e$ and $\hat{Q}^\varnothing$ with 0-indexing stored as single square matrix $\hat{Q}$.** Using production code conventions:

- **0-indexing.** The nucleotides in a complex of $N$ nucleotides are indexed $j \in [0 : N-1]$ starting at $j = 0$.
- **Shifted recursion elements for subsequence $[i, j]$.** We shift the column index $j \to j+1$ so subsequence $[i, j]$ is represented by recursion elements $\hat{Q}^e_{i,j+1}$ and $\hat{Q}^\varnothing_{i,j+1}$, each a $\gamma \times \gamma$ matrix.
- **Shifted 0-indexed triangular recursion matrices.** The shifted triangular matrix $\hat{Q}^e$ with 0-indexing has columns $j \in [N]$ and rows $i \in [0 : j-1]$; the shifted triangular matrix $\hat{Q}^\varnothing$ has columns $j \in [N]$ and rows $i \in [0 : j-1]$ augmented by the base case $\hat{Q}^\varnothing_{j,j}$ corresponding to empty subsequence $[j, j-1]$.
- **Single square recursion matrix $\hat{Q}$.** The shifted 0-indexed triangular matrices can then be stored as a single $(N+1) \times (N+1)$ matrix $\hat{Q}$, with $\hat{Q}^\varnothing$ stored above the diagonal (with the base case on the diagonal) and $(\hat{Q}^e)^\top$ stored below the diagonal. For subsequence $[i, j]$ we then have:

```
for all j ∈ [0:N − 1] do                                   ▷ 3′-most base j
    Q̂_{j,j} ← Q̂^∅_{j,j}            ▷ On diagonal: base case for empty subsequence [j, j−1]
    for all i ∈ [0:j] do                                    ▷ 5′-most base i
        Q̂_{i,j+1} ← Q̂^∅_{i,j+1}                            ▷ Above diagonal: Q̂^∅
        Q̂_{j+1,i} ← (Q̂^e_{i,j+1})^⊤                ▷ Below diagonal: transpose of Q̂^e
```

#### S6.2 Operation order for partition function, number of subensemble states, MFE, and gradient of the partition function

For a complex with sequence $\phi$, the generic recursions of Figure S1 are combined with the operation order of Figure S2 to calculate either the partition function $\overline{Q}(\phi)$ (using the SumProduct evaluation algebra), the number of subensemble states $|\overline{\Gamma}^{\shortparallel}(\phi)|$ (using the Count evaluation algebra), or the free energy of the MFE stacking state $\overline{\Delta G}(\phi, s^{\shortparallel}_{\text{MFE}}(\phi))$ (using the MinSum evaluation algebra). The same operation order can be used to calculate the gradient of the partition function using the $\partial$SumProduct evaluation algebra, yielding both the partition function, $\overline{Q}(\phi)$, and its gradient, $\partial\overline{Q}(\phi)/\partial\theta$, with respect to a specified parameter, $\theta$. In the case of multiple parameters $(\theta_1, \theta_2, \dots)$, we avoid recomputation of partition function contributions by first computing the partition function using the SumProduct evaluation algebra and then computing the derivative for each parameter of interest using the $\partial$SumProduct evaluation algebra, looking up partition function intermediate results when possible to avoid recomputation.

- **Pseudocode with main text conventions:** Pseudocode detailing the operation order for calculating the partition function is shown using main text conventions in Algorithm S1 (operating on $Q^e$ and $Q^\varnothing$). Note that within column $j$ of $Q^\varnothing$, the entry for each row $i \in [j]$ can be calculated independently; since the complex partition function is calculated using the top right corner entry, $\text{Tr}_S(Q^\varnothing_{1,N})$, of column $N$, there is no need to calculate the other entries, $Q^\varnothing_{[2:N],N}$, in column $N$.
- **Pseudocode with production code and $\chi$ conventions:** Pseudocode using production code conventions is shown in Algorithm S2 (operating on $\hat{Q}$). In practice, the production code conventions are implemented by defining a function $\chi$ that encompasses the *extend* operation, $\hat{Q}^e$, and the *combine* operation base case $\hat{Q}^\varnothing_{j,j}$ (corresponding to empty subsequence $[j, j-1]$), shown in the pseudocode of Algorithm S3 (operating on $\hat{Q}$). Note that within column $j$ of $\hat{Q}$, the entry for each row $i \in [0:j]$ can be calculated independently; since the complex partition function is calculated using the top right corner entry, $\text{Tr}_S(\hat{Q}_{0,N})$, of column $N$, there is no need to calculate the other entries, $\hat{Q}_{[1:N-1],N}$, in column $N$.

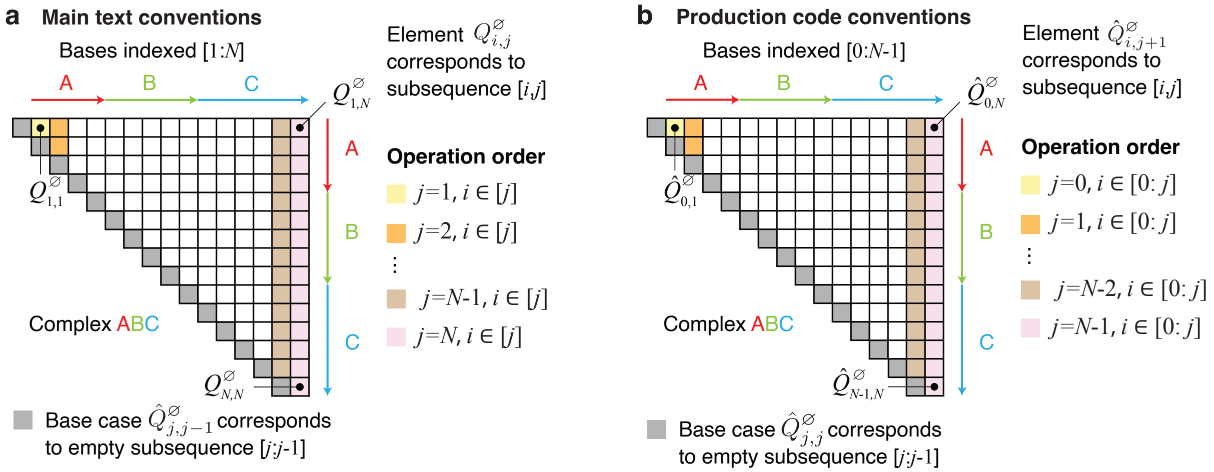

**Figure S2:** Operation order for calculation of the partition function, number of states, or MFE. (a) Main text conventions (Section S4.1; see the pseudocode of Algorithm S1). A forward sweep of 3′-most base $j$ from 1 to $N$, considering all subsequences $[i, j]$ for each $j$. For a given subsequence $[i, j]$, $Q^e_{i,j}$ is calculated (an *extend* operation) prior to calculating $Q^\varnothing_{i,j}$ (a *combine* operation); the base case $Q^\varnothing_{j,j-1}$ corresponds to empty subsequence $[j, j-1]$. (b) Production code conventions (Section S6.1; see the pseudocode of Algorithm S2). A forward sweep of 3′-most base $j$ from 0 to $N-1$, considering all subsequences $[i, j]$ for each $j$. For a given subsequence $[i, j]$, $\hat{Q}^e_{i,j+1}$ is calculated (an *extend* operation; with $(\hat{Q}^e_{i,j+1})^\top$stored below the diagonal as $\hat{Q}_{j+1,i}$) prior to calculating $\hat{Q}^\varnothing_{i,j+1}$ (a *combine* operation; stored above the diagonal as $\hat{Q}_{i,j+1}$); the base case $\hat{Q}^\varnothing_{j,j}$ (stored on the diagonal as $\hat{Q}_{j,j}$) corresponds to empty subsequence $[j, j-1]$.

**Algorithm S2** Optimized $O(N^3)$ algorithm to calculate the complex partition function $\overline{Q}(\phi)$ with reduced time-complexity prefactor using production code conventions (cf. main text conventions of Algorithm S1 and $\chi$ conventions of Algorithm S3).

```
function ComplexPartitionFunction(N, V, B)
    for j ∈ [0:N − 1] do                                        ▷ Loop over 3′-most base j
        Q̂_{j,j} ← I                                             ▷ Base case for empty subsequence [j, j−1]
        for all i ∈ [0:j−1] do                                  ▷ Loop over 5′-most base i
            Q̂_{j+1,i} ← (∑_{ρ∈ξ} B^ρ_{i,j} Tr_S[B^ρ_{j,i} Q̂_{i+1,j}])^⊤   ▷ Extend subsequence to j with base pair i · j
        Q̂_{j+1,j} ← (V_j)^⊤                                     ▷ Extend subsequence to j with unpaired base j
        for all i ∈ [0:j] do
            Q̂_{i,j+1} ← ∑_{k∈[i:j]} Q̂_{i,k} (Q̂_{j+1,k})^⊤       ▷ Combine operation
    return Tr_S(Q̂_{0,N})                                        ▷ Return partition function Q̄(φ)
```

**Algorithm S3** Optimized $O(N^3)$ algorithm to calculate the complex partition function $\overline{Q}(\phi)$ with reduced time-complexity prefactor using $\chi$ conventions (cf. main text conventions of Algorithm S1 and production code conventions of Algorithm S2).

```
function ComplexPartitionFunction(N, V, B)
    for j ∈ [0 : N] do               ▷ Loop over matrix columns including base case j=0; 3′-most base is j−1
        for all i ∈ [0 : j] do                                  ▷ Loop over 5′-most base i
            Q̂_{j,i} ← χ(V, B, i, j, Q̂)                          ▷ Extend operation
        for all i ∈ [0 : j − 1] do
            Q̂_{i,j} ← ∑_{k∈[i:j−1]} Q̂_{i,k} (Q̂_{j,k})^⊤         ▷ Combine operation
    return Tr_S(Q̂_{0,N})                                        ▷ Return partition function Q̄(φ)

function χ(V, B, i, j, Q̂)                      ▷ Extend operation: i is 5′-most base; j − 1 is 3′-most base
    if i = j then
        q ← I                                                   ▷ Base case for empty subsequence [j, j−1]
    else if i = j − 1 then
        q ← V_i                                 ▷ Extend subsequence to j−1 with unpaired base j−1
    else
        q ← ∑_{ρ∈ξ} B^ρ_{i,j−1} Tr_S(B^ρ_{j−1,i} Q̂_{i+1,j−1})   ▷ Extend subsequence to j−1 with base pair i · j−1
    return q^⊤                              ▷ Return transpose to be stored as Q̂_{j,i} on and below the diagonal
```

#### S6.3 Partition function implementation details

##### S6.3.1 Renormalization to prevent partition function overflow

Historically, it has been a significant challenge to calculate the partition function while preventing overflow[13, 19] as the size of the complex, $N \coloneqq |\phi|$, increases. Existing algorithms based on NN free energy models achieve scalable partition function calculations by employing an overflow-safe evaluation algebra that operates separately on the mantissa and exponent.[19] Here, we define a simpler renormalization approach for preventing partition function overflow that capitalizes on the operation order used for the new dynamic programs (Figures 7a and S1a), in which a new 3′-most base $j$ is introduced at each iteration.

**Renormalization using main text conventions.** Suppose that partition function contributions associated with base $j$ are weighted by a constant scalar, $x_j$. The partition function elements $Q^e_{i,j}$ and $Q^\varnothing_{i,j}$ for subsequence $[i, j]$ would then incorporate the scale factor $\prod_{k=i}^{j} x_k$, corresponding to a single factor of $x_k$ for each base $k$ in the subsequence. Using the dynamic program of Algorithm S1, since all subsequences $[i, j]$ incorporating 3′-most base $j$ are incorporated during a single iteration of the $Q^e_{i,j}$ recursion for $i \in [j]$, the renormalization for the first and last bases $i$ and $j$ can be applied for all such subsequences by explicitly multiplying $Q^e_{i,j}$ by $x_i$ and $x_j$ (illustrated in Figure S3a). For the intervening bases $[i+1, j-1]$, renormalization of $Q^e_{i,j}$ by the factor $\prod_{k=i+1}^{j-1} x_k$ is achieved implicitly via incorporation of the previously calculated partition function element $Q^\varnothing_{i+1,j-1}$ for subsequence $[i+1, j-1]$. Note that $Q^\varnothing_{i,j}$ is likewise normalized implicitly by the factor $\prod_{k=i}^{j} x_k$ via incorporation of $Q^e$ and $Q^\varnothing$ elements previously calculated for shorter subsequences.

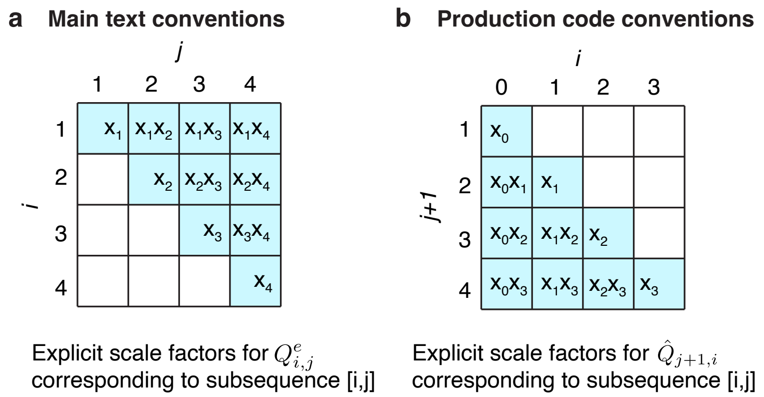

**Figure S3:** Explicitly applied partition function renormalization constants. (a) Main text conventions (Section S4.1; see the pseudocode of Algorithm S4): explicitly applied renormalization constant for element $Q^e_{i,j}$ corresponding to subsequence $[i, j]$. For example, the explicitly applied renormalization constant for element $Q^e_{1,3}$ corresponding to subsequence $[1, 3]$ is $x_1 x_3$. $x_k$ is the renormalization constant for base $k \in [N]$. (b) Production code conventions (Section S6.1; see the pseudocode of Algorithm S5): explicitly applied renormalization constant for element $\hat{Q}_{j+1,i}$ corresponding to subsequence $[i, j]$. For example, the explicitly applied renormalization constant for element $\hat{Q}_{3,0}$ corresponding to subsequence $[0, 2]$ is $x_0 x_2$. $x_k$ is the renormalization constant for base $k \in [0\,{:}\,N-1]$.

---

**Algorithm S4** Renormalization for prevention of partition function overflow using main text conventions (cf. production code conventions of Algorithm S5 and $\chi$ conventions of Algorithm S6)

---

**procedure** RENORMALIZE($Q^e, \vec{x}, j$) ▷ *$\vec{x}$ vector of renormalization constants: $x_k$ for base $k \in [N]$*
  **for all** $i \in [j-1]$ **do** ▷ *$i$ is 5′-most base; $j$ is 3′-most base*
    $Q^e_{i,j} \leftarrow Q^e_{i,j} \cdot x_i$ ▷ *Explicitly apply scale for 5′-most base $i$; elementwise multiplication*
  $y \leftarrow \max(Q^e_{[j],j})$ ▷ *Maximum over each scalar entry in $Q^e_{[j],j}$*
  **if** $y > 0$ **then**
    $x_j \leftarrow 1/y$
  **else**
    $x_j \leftarrow 1$
  $Q^e_{[j],j} \leftarrow Q^e_{[j],j} \cdot x_j$ ▷ *Explicitly apply scale for 3′-most base $j$; elementwise multiplication*

---

This renormalization approach is detailed in the pseudocode of Algorithm S4. $\vec{x}$ is a vector of renormalization constants, $x_k$, for $k \in [N]$. For subsequence $[i, j]$, first $Q^e_{i,j}$ is renormalized by the previously calculated renormalization factor, $x_i$, corresponding to 5′-most base $i$. Then the maximum of $Q^e_{i,j}$ for $i \in [j]$ is calculated (corresponding to all subsequences ending at $j$), inverted to define $x_j$, and used to renormalize $Q^e_{i,j}$ for all subsequences $[i, j]$. Note that scalar scale factors $x_i$ and $x_j$ are applied to each matrix entry $Q^{e,a,b}_{i,j}$ corresponding to $a \in [\gamma]$ and $b \in [\gamma]$.

The unnormalized value of any element may be calculated by multiplying by the reciprocal scale factor (e.g., $Q^{\varnothing}_{i,j} \prod_{k=i}^{j} x_k^{-1}$). To avoid overflow, the unnormalized complex free energy can be reported

$$\overline{g}(\phi) = -\log \mathrm{Tr}_S(Q^{\varnothing}_{1,N}) + \sum_{k=1}^{N} \log x_k \tag{S47}$$

instead of reporting the unnormalized complex partition function

$$\overline{Q}(\phi) = \mathrm{Tr}_S(Q^{\varnothing}_{1,N}) \prod_{k=1}^{N} x_k^{-1}. \tag{S48}$$

Suppose the previously-calculated elements of $Q^e$ are well-normalized (e.g., ≈1 in magnitude). The only way for overflow to occur in the next step of the normalized algorithm is for the change in the free energy of the complex associated with a new 3′-most base $j$ to be more favorable than roughly $g_{\max} = -k_B T \log(f_{\max})$, where $k_B$ is the Boltzmann constant, $T$ is temperature, and $f_{\max}$ is the maximum expressible floating point number. Using single-precision, $f_{\max} \approx 10^{38}$ and $g_{\max} \approx -50$ kcal/mol, while for double-precision $f_{\max} \approx 10^{308}$ and $g_{\max} \approx -500$ kcal/mol,

**Algorithm S5** Renormalization for prevention of partition function overflow using production code conventions (cf. main text conventions of Algorithm S4 and $\chi$ conventions of Algorithm S6)

```
procedure RENORMALIZE(Q̂, x⃗, j)        ▷ x⃗ vector of renormalization constants: x_k for base k ∈ [0:N−1]
    for all i ∈ [0:j−1] do                            ▷ i is 5′-most base; j is 3′-most base
        Q̂_{j+1,i} ← Q̂_{j+1,i} · x_i     ▷ Explicitly apply scale for 5′-most base i; elementwise multiplication
    y ← max(Q̂_{j+1,[0:j]})                       ▷ Maximum over each scalar entry in Q̂_{j+1,[0:j]}
    if y > 0 then
        x_j ← 1/y
    else
        x_j ← 1
    Q̂_{j+1,[0:j]} ← Q̂_{j+1,[0:j]} · x_j   ▷ Explicitly apply scale for 3′-most base j; elementwise multiplication
```

**Algorithm S6** Renormalization for prevention of partition function overflow using $\chi$ conventions (cf. main text conventions of Algorithm S4 and production code conventions of Algorithm S5).

```
procedure RENORMALIZE(Q̂, x⃗, j)        ▷ x⃗ vector of renormalization constants: x_k for base k ∈ [0:N−1]
    if j > 0 then
        for all i ∈ [0:j−2] do                          ▷ i is 5′-most base; j−1 is 3′-most base
            Q̂_{j,i} ← Q̂_{j,i} · x_i     ▷ Explicitly apply scale for 5′-most base i; elementwise multiplication
        y ← max(Q̂_{j,[0:j−1]})                     ▷ Maximum over each scalar entry in Q̂_{j,[0:j−1]}
        if y > 0 then
            x_{j−1} ← 1/y
        else
            x_{j−1} ← 1
    Q̂_{j,[0:j−1]} ← Q̂_{j,[0:j−1]} · x_{j−1}   ▷ Explicitly apply scale for 3′-most base j−1; elementwise multiplication
```

so overflow is prevented for all practical purposes using realistic free energy models with either single- or double-precision.

Note that the renormalization scales, $x_i$ for all $i \in [N]$, that are used in the forward pass to perform an overflow-safe calculation of the partition function using Algorithm S4 are also used to renormalize $Q^e$ recursion elements in the reverse pass to perform an overflow-safe calculation of the equilibrium pair probability matrix using Algorithm S10.

**Renormalization using production code and $\chi$ conventions.** Using production code conventions, the partition function elements for subsequence $[i,j]$, $\hat{Q}^e_{i,j+1}$ (with $(\hat{Q}^e_{i,j+1})^\top$ stored below the diagonal as $\hat{Q}_{j+1,i}$), and $\hat{Q}^\varnothing_{i,j+1}$ (stored above the diagonal as $\hat{Q}_{i,j+1}$), would then incorporate the scale factor $\prod_{k=i}^{j} x_k$, corresponding to a single factor of $x_k$ for each base $k$ in the subsequence. Renormalization is achieved by explicitly multiplying element $\hat{Q}^e_{i,j+1}$ (with $(\hat{Q}^e_{i,j+1})^\top$ stored below the diagonal as $\hat{Q}_{j+1,i}$) for subsequence $[i,j]$ by $x_i$ and $x_j$ (illustrated in Figure S3b), with all other renormalization achieved implicitly. Renormalization pseudocode using production code conventions is displayed in Algorithm S5 and using $\chi$ conventions in Algorithm S6. $\vec{x}$ is a vector of renormalization constants, $x_k$, for $k \in [0:N-1]$.

For subsequence $[i,j]$, the unnormalized value of $\hat{Q}^e_{i,j+1}$ (with $(\hat{Q}^e_{i,j+1})^\top$stored as $\hat{Q}_{j+1,i}$ below the diagonal) or of $\hat{Q}^\varnothing_{i,j+1}$ (stored as $\hat{Q}_{i,j+1}$ above the diagonal) may be calculated by multiplying by the reciprocal scale factor $\prod_{k=i}^{j} x_k^{-1}$ (e.g., $\hat{Q}_{0,N} \prod_{k=0}^{N-1} x_k^{-1}$ for the full sequence $[0,N-1]$). To avoid overflow, the unnormalized complex free energy can be reported

$$\overline{g}(\phi) = -\log \mathrm{Tr}_S(\hat{Q}_{0,N}) + \sum_{k=0}^{N-1} \log x_k \tag{S49}$$

instead of reporting the unnormalized complex partition function

$$\overline{Q}(\phi) = \mathrm{Tr}_S(\hat{Q}_{0,N}) \prod_{k=0}^{N-1} x_k^{-1}. \tag{S50}$$

##### S6.3.2 Partition function algorithm with renormalization and GPU acceleration

Here, we describe the production partition function algorithm, which incorporates renormalization to prevent overflow and optional GPU acceleration to reduce wall clock time.

**Partition function algorithm using main text conventions: renormalization and GPU acceleration.** Algorithm S7 iterates three operations:

- *Extend* the subsequence to a new 3′-most base $j$ by calculating $Q^e_{i,j}$.
  - Time complexity: $O(\xi\gamma^2 N^2)$.
  - Implementation: parallelization over CPU threads for each independent $i \in [j]$.
- *Renormalize* $Q^e$ to avoid overflow,
  - Time complexity: $O(\gamma^2 N^2)$.
  - Implementation: evaluate using CPU.
- *Combine* $Q^{\varnothing}_{i,k-1}$ and $Q^e_{k,j}$ to calculate the partition function for subsequence $[i, j]$.
  - Time complexity: $O(\gamma^3 N^3)$
  - Implementation: evaluate using CPU or GPU as a triangular matrix multiply BLAS routine as described below. For the GPU implementation, there is a data transfer overhead of $O(\gamma^2 N^2)$.

While existing dynamic programming algorithms for calculating the partition function over the ensemble of a complex exploited vectorized dot products for efficient CPU implementation,[19] here we go a step further and demonstrate that the *combine* operation, which is the sole component of the algorithm with $O(N^3)$ time complexity, can be implemented as a single matrix product, enabling exploitation of highly optimized matrix primitives (either BLAS routines for CPU implementation or cuBLAS routines for CUDA GPU implementation).

---

**Algorithm S7** Optimized $O(N^3)$ algorithm to calculate the complex partition function $\overline{Q}(\phi)$ with renormalization to avoid overflow and efficient linear algebra primitives with optional GPU acceleration using main text conventions (cf. production code conventions of Algorithm S8 and $\chi$ conventions of Algorithm S9).

---

**function** ComplexPartitionFunction$(N, V, B)$
  **for** $j \in [N]$ **do**
    $Q^{\varnothing}_{j,j-1} \leftarrow \mathbf{I}$ ▷ *Base case for empty subsequence* $[j, j-1]$
    **if** using GPU and $j > 2$ **then** get $Q^{\varnothing}_{[2:j-1],j-1}$ from GPU ▷ $O(\gamma^2 N)$ *data transfer*
    **for all** $i \in [j-1]$ **do**
      $Q^e_{i,j} \leftarrow \sum_{\rho\in\xi} B^{\rho}_{i,j} \mathrm{Tr}_S[B^{\rho}_{j,i} Q^{\varnothing}_{i+1,j-1}]$ ▷ *Extend operation: subsequence to j with base pair i · j*
    $Q^e_{j,j} \leftarrow V_j$ ▷ *Extend operation: subsequence to j with unpaired base j*
    Renormalize$(Q^e, \vec{x}, j)$ ▷ *Renormalize to avoid overflow using Algorithm S4*
    **if** using GPU **then** send $Q^{\varnothing}_{j,j-1}$ and $Q^e_{[j],j}$ to GPU ▷ $O(\gamma^2 N)$ *data transfer*
    $Q^{\varnothing}_{[j],j} \leftarrow$ Trmm$(Q^{\varnothing}_{[j],[0:j-1]}, Q^e_{[j],j})$ ▷ *Combine operation: multiply triangular* $Q^{\varnothing}$ *by vector* $Q^e$
  **return** $\mathrm{Tr}_S(Q^{\varnothing}_{1,N}) \prod_{k=1}^{N} x_k^{-1}$ ▷ *Return unnormalized partition function* $\overline{Q}(\phi)$

---

Consider the *combine* operation

$$Q^{\varnothing}_{i,j} = \sum_{k\in[i:j]} Q^{\varnothing}_{i,k-1} Q^e_{k,j} \tag{S51}$$

for subsequence $[i, j]$. Recall that each recursion element, $Q^{\varnothing}_{i,k-1}$ or $Q^e_{k,j}$, is a $\gamma \times \gamma$ matrix. Hence, this can be rewritten

$$\begin{aligned} Q^{\varnothing\,[a,b]}_{i,j} &= \sum_{k\in[i:j]} \sum_{c\in[\gamma]} Q^{\varnothing\,[a,c]}_{i,k-1} Q^{e\,[c,b]}_{k,j} \\ &= \sum_{k\in[j]} \sum_{c\in[\gamma]} Q^{\varnothing\,[a,c]}_{i,k-1} Q^{e\,[c,b]}_{k,j}\, \delta(k \geq i). \end{aligned} \tag{S52}$$

Let $Q^{\varnothing\prime}$ denote the raw matrix corresponding to $Q^{\varnothing}$ in memory such that $Q^{\varnothing\,[a,b]}_{i,j} = {Q^{\varnothing\prime}}_{\gamma i+a,\gamma j+b}$. Then (S52) may be rewritten

$$ {Q^{\varnothing\prime}}_{\gamma i+a,\gamma j+b} = \sum_{k\in[\gamma j]} {Q^{\varnothing\prime}}_{\gamma i+a,k-\gamma}\, {Q^{e\prime}}_{k,\gamma j+b}\, \delta(k \geq \gamma i), \tag{S53} $$

which can be efficiently implemented using optimized BLAS or cuBLAS routines[4]

$$ {Q^{\varnothing\prime}}_{\gamma[j]+a,\gamma j+b} = \textsc{Trmm}({Q^{\varnothing\prime}}_{\gamma[j]+a,[\gamma j]-\gamma}, {Q^{e\prime}}_{[\gamma j],\gamma j+b}) \tag{S54} $$

where $\textsc{Trmm}(A,B) := U(A)B$ with $U_{i,j}(X) := X_{i,j}\,\delta(j \geq i)$ corresponds to multiplication of an upper triangular matrix by a rectangular matrix. In this instance, the upper triangular matrix is $U({Q^{\varnothing\prime}}_{\gamma[j]+a,[\gamma j]-\gamma})$ and the rectangular matrix is the vector ${Q^{e\prime}}_{[\gamma j],\gamma j+b}$. In practice, $Q^e$ and $Q^{\varnothing}$ matrices are stored in blocks according to the layout of $S$, $V$, and $B$, yielding analogous $\textsc{Trmm}$ calls.

**Partition function algorithm using production code and $\chi$ conventions: renormalization and GPU acceleration.** Algorithm S8 uses production code conventions and Algorithm S9 uses $\chi$ conventions. Consider the *combine* operation

$$ \hat{Q}_{i,j+1} = \sum_{k\in[i:j]} \hat{Q}_{i,k} (\hat{Q}_{j+1,k})^\top \tag{S55} $$

for subsequence $[i,j]$. Recall that each recursion element, $\hat{Q}_{i,k}$ or $(\hat{Q}_{j+1,k})^\top$, is a $\gamma \times \gamma$ matrix. Hence, this can be rewritten

$$ \begin{aligned} Q^{[a,b]}_{i,j+1} &= \sum_{k\in[i:j]} \sum_{c\in[0:\gamma-1]} \hat{Q}^{[a,c]}_{i,k} (\hat{Q}_{j+1,k})^{\top\,[c,b]} \\ &= \sum_{k\in[0:j]} \sum_{c\in[0:\gamma-1]} \hat{Q}^{\varnothing\,[a,c]}_{i,k} (\hat{Q}_{j+1,k})^{\top\,[c,b]}\, \delta(k \geq i). \end{aligned} \tag{S56} $$

Let $\hat{Q}'$ denote the raw matrix corresponding to $\hat{Q}$ in memory such that $\hat{Q}^{[a,b]}_{i,j} = \hat{Q}'_{\gamma i+a,\gamma j+b}$. Then (S56) may be rewritten

$$ \hat{Q}'_{\gamma i+a,\gamma(j+1)+b} = \sum_{k\in[0:\gamma(j+1)-1]} \hat{Q}'_{\gamma i+a,k}\, \hat{Q}'_{\gamma(j+1)+b,k}\, \delta(k \geq \gamma i), \tag{S57} $$

which can be efficiently implemented using optimized BLAS routines[4]

$$ \hat{Q}'_{\gamma[0:j]+a,\gamma(j+1)+b} = \textsc{Trmm}(\hat{Q}'_{\gamma[0:j]+a,[0:\gamma(j+1)-1]}, (\hat{Q}'_{\gamma(j+1)+b,[0:\gamma(j+1)-1]})^\top). \tag{S58} $$

---

**Algorithm S8** Optimized $O(N^3)$ algorithm to calculate the complex partition function $\overline{Q}(\phi)$ with renormalization to avoid overflow and efficient linear algebra primitives with optional GPU acceleration using production code conventions (cf. main text conventions of Algorithm S7 and $\chi$ conventions of Algorithm S9).

---

**function** ComplexPartitionFunction($N, V, B$)
  **for** $j \in [0:N-1]$ **do**
    $\hat{Q}_{j,j} \leftarrow \mathbf{I}$ ▷ *Base case for empty subsequence* $[j, j-1]$
    **if** using GPU and $j > 1$ **then** get $\hat{Q}_{[1:j-1],j}$ from GPU ▷ $O(\gamma^2 N)$ *data transfer*
    **for all** $i \in [0:j-1]$ **do**
      $\hat{Q}_{j+1,i} \leftarrow (\sum_{\rho\in\xi} B^{\rho}_{i,j} \mathrm{Tr}_S[B^{\rho}_{j,i} \hat{Q}_{i+1,j}])^\top$ ▷ *Extend operation: subsequence to* $j$ *with base pair* $i \cdot j$
    $\hat{Q}_{j+1,j} \leftarrow (V_j)^\top$ ▷ *Extend operation: subsequence to* $j$ *with unpaired base* $j$
    Renormalize($\hat{Q}, \vec{x}, j$) ▷ *Renormalize to avoid overflow using Algorithm S5*
    **if** using GPU **then** send $\hat{Q}_{j,j}$ and $\hat{Q}_{j+1,[0:j]}$ to GPU ▷ $O(\gamma^2 N)$ *data transfer*
    $\hat{Q}_{[0:j],j+1} \leftarrow \textsc{Trmm}(\hat{Q}_{[0:j],[0:j]}, (\hat{Q}_{j+1,[0:j]})^\top)$ ▷ *Combine operation: multiply triangular matrix by vector*
  **return** $\mathrm{Tr}_S(\hat{Q}_{0,N}) \prod_{k=0}^{N-1} x_k^{-1}$ ▷ *Return unnormalized partition function* $\overline{Q}(\phi)$

---

**Algorithm S9** Optimized $O(N^3)$ algorithm to calculate the complex partition function $\overline{Q}(\phi)$ with renormalization to avoid overflow and efficient linear algebra primitives with optional GPU acceleration using $\chi$ conventions (cf. main text conventions of Algorithm S7 and production code conventions of Algorithm S8).

```
function ComplexPartitionFunction(N, V, B)
    for j ∈ [0 : N] do                         ▷ Loop over matrix columns including base case j=0; 3′-most base is j−1
        if using GPU and j > 1 then get Q̂_{[1:j−1],j−1} from GPU       ▷ O(γ²N) data transfer
        for all i ∈ [0 : j] do                                          ▷ Loop over 5′-most base i
            Q̂_{j,i} ← χ(V, B, i, j, Q̂)                                  ▷ Extend operation using χ of Algorithm S3
        Renormalize(Q̂, x⃗, j)                                  ▷ Renormalize to avoid overflow using Algorithm S6
        if using GPU then send Q̂_{j,[0:j]} to GPU                       ▷ O(γ²N) data transfer
        Q̂_{[0:j−1],j} ← Trmm(Q̂_{[0:j−1],[0:j−1]}, (Q̂_{j,[0:j−1]})^⊤)     ▷ Combine operation
    return Tr_S(Q̂_{0,N}) ∏_{k=0}^{N−1} x_k^{−1}          ▷ Return unnormalized partition function Q̄(φ)
```

#### S6.4 Backtrack-free operation order for equilibrium pair probability matrix

For a complex of $N$ nucleotides with sequence $\phi$, the equilibrium pair probability matrix, $\overline{P}(\phi)$, is a symmetric $N \times N$ matrix with off-diagonal entries, $\overline{P}_{i,j}(\phi) = \overline{P}_{j,i}(\phi)$, denoting the probability that base pair $i \cdot j$ forms at equilibrium over complex ensemble $\overline{\Gamma}(\phi)$, and diagonal entries, $\overline{P}_{i,i}(\phi)$, denoting the probability that base $i$ is unpaired at equilibrium over complex ensemble $\overline{\Gamma}(\phi)$. Each row sum is one.

Historically, using NN free energy models, equilibrium base-pairing probabilities for a single strand[14, 43] or a complex[13] were calculated using a dynamic program that backtracked through the matrix of subsequence partition functions. This backtracking process involved subtraction of intermediate partition function quantities, creating the risk of losing precision due to subtraction of large numbers differing by a small amount.

Subsequently, for NN free energy models, Fornace et al.[19] showed that equilibrium pair probabilities can be calculated without backtracking using a forward sweep of the (overflow-safe) partition function algorithm on a "doubled" sequence. To see how this works, consider a complex with strand ordering $\pi$ = AB and a total of $N$ nucleotides. As an intermediate result, the NN partition function algorithm calculates $Q^b_{i,j}$, the conditional partition function for the nucleotides internal to subsequence $[i, j]$ (inclusive), namely nucleotides $i, i+1, \ldots, j-1, j$, subject to the constraint that $i$ is paired to $j$. Similarly, consider the conditional partition function, $Q^{b_{\text{ext}}}_{i,j}$, for the nucleotides external to subsequence $[i, j]$ (inclusive), namely nucleotides $j, \ldots, N-1, 0, \ldots, i$ (Figure S4a; bottom). Because the structural ensemble $\overline{\Gamma}(\phi)$ excludes pseudoknots, the base pair $i \cdot j$ partitions the structural ensemble into non-interacting internal and external subensembles, so the partition function of all structures containing base pair $i \cdot j$ is the product $Q^b_{i,j} Q^{b_{\text{ext}}}_{i,j}$. As a result, the equilibrium probability of base pair $i \cdot j$ over ensemble $\overline{\Gamma}(\phi)$ may be expressed for $i < j$:

$$\overline{P}_{i,j}(\phi) = Q^b_{i,j}(\phi) Q^{b_{\text{ext}}}_{i,j}(\phi) / \overline{Q}(\phi). \qquad \text{(NN model)} \tag{S59}$$

Mathews et al.[40] employed this approach using new recursions to calculate the external conditional partition function $Q^{b_{\text{ext}}}_{i,j}$ for a single strand. Fornace et al.[19] addressed the general case of a complex of $L$ strands and show that $Q^{b_{\text{ext}}}_{i,j}$ can be calculated without new recursions by replicating the strands to form a "doubled" complex with sequence $\phi'$ (e.g., $\pi$ = ABAB) containing $2N$ nucleotides, and calculating $Q^b_{i,j}$ using the standard recursions for all subsequences of up to $N$ nucleotides. The external subsequence $j, \ldots, N, 1, \ldots, i$ for the original complex with sequence $\phi$ (Figure S4a; bottom) is simply the internal subsequence $j, \ldots, N+i$ for the doubled complex with sequence $\phi'$ (Figure S4b; bottom). Hence, the equilibrium pair probability matrix can be calculated without backtracking for $i < j$ using:

$$\overline{P}_{i,j}(\phi) = Q^b_{i,j}(\phi) Q^b_{j,N+i}(\phi') / \overline{Q}(\phi). \qquad \text{(NN model)} \tag{S60}$$

In Figure S4b, the pink matrix elements are cached from a previous complex partition function calculation of $Q^b_{i,j}(\phi)$ on sequence $\phi$ (Figure S4a) and the blue entries correspond to calculation of $Q^b_{j,N+i}(\phi')$ on doubled sequence $\phi'$.

Here, we adapt this backtrack-free approach to the new transfer matrix and tensor models. Expressed in terms of $\hat{Q}^e$ and $\hat{Q}^\varnothing$ using production code conventions (recursion elements for subsequence $[i, j]$ are $\hat{Q}^e_{i,j+1}$ and $\hat{Q}^\varnothing_{i,j+1}$; see Section S6.1), the analogous form of (S60) for $i < j$ is:

$$\overline{P}_{i,j}(\phi) = \text{Tr}_S(\hat{Q}^e_{i,j+1}(\phi) \hat{Q}^\varnothing_{j+1,N+i}(\phi')) / \overline{Q}(\phi). \tag{S61}$$

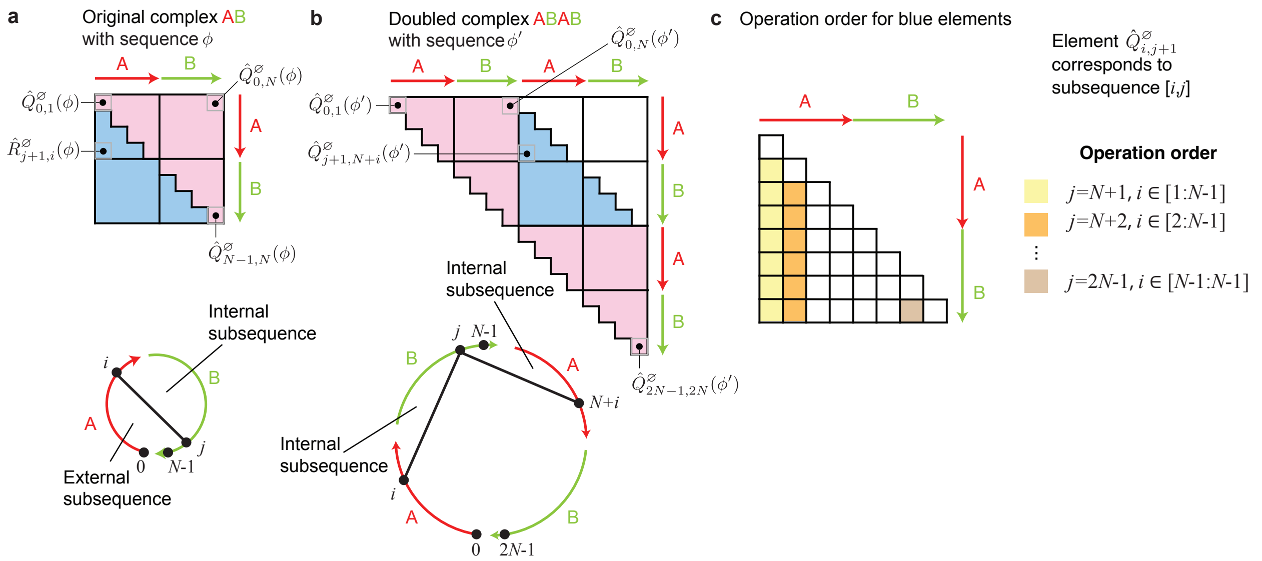

**Figure S4:** Backtrack-free operation order for calculation of equilibrium pair probability matrix. **(a)** Complex AB of $N$ nucleotides with sequence $\phi$. **(b)** "Doubled complex" ABAB of $2N$ nucleotides with sequence $\phi'$. Top: pink elements available from a previous partition function calculation for sequence $\phi$; blue elements calculated using cyclic indexing for sequence $\phi$ (panel a) or without cyclic indexing on doubled sequence $\phi'$ (panel b). Bottom: exterior subsequence created by base pair $i \cdot j$ for sequence $\phi$ with cyclic indexing (panel a) is equivalent to interior subsequence created by base pair $j \cdot N+i$ for doubled sequence $\phi'$ without cyclic indexing (panel b). **(c)** Forward sweep operation order for calculating blue elements; for a given exterior subsequence $j+1,\ldots,N-1,0,i-1$ using cyclic indexing (panel a), $\hat{R}^e_{i,j+1}$ is calculated (an *extend* operation; with $(\hat{R}^e_{i,j+1})^\top$stored on and above the diagonal as $\hat{R}_{j+1,i}$) prior to calculating $\hat{R}^{\varnothing}_{i,j+1}$ (a *combine* operation; stored below the diagonal as $\hat{R}_{i,j+1}$). Production code conventions (see Section S6.1).

In this expression, $\hat{Q}^e_{i,j+1}$ denotes the constrained partition function elements for interior subsequence $[i,j]$ of sequence $\phi$ including the contributions for base pair $i \cdot j$ (using a $B$ transfer matrix or tensor on each side of the base pair) and $\hat{Q}^{\varnothing}_{j+1,N+i}$ denotes the unconstrained partition function elements for interior subsequence $[j+1, N+i-1]$ of doubled sequence $\phi'$.

Note that when $i = j$, $\hat{Q}^e_{i,j+1}$ denotes the constrained partition function elements for interior subsequence $[i,j]$ of sequence $\phi$ including the contributions for unpaired base $i$ (using a $V$ transfer matrix) and $\hat{Q}^{\varnothing}_{j+1,N+i}$ denotes the unconstrained partition function elements for interior subsequence $[j+1, N+i-1]$ of doubled sequence $\phi'$. Hence, (S61) remains valid for $i = j$, and yields the probability that base $i$ is unpaired at equilibrium, $\overline{P}_{i,i}(\phi)$.

In practice, we avoid explicitly forming doubled sequence $\phi'$ by instead employing cyclic indexing on sequence $\phi$ (i.e., base index $N+i$ for $\phi'$ equivalent to base index $i$ for $\phi$). With cyclic indexing, (S61) may be restated for $i \le j$:

$$\overline{P}_{i,j}(\phi) = \mathrm{Tr}_S(\hat{Q}^e_{i,j+1}(\phi)\hat{R}^{\varnothing}_{j+1,i}(\phi))/\overline{Q}(\phi), \tag{S62}$$

where $\hat{R}^{\varnothing}_{j+1,i}(\phi)$ denotes the unconstrained partition function elements for exterior subsequence $j+1,\ldots,N-1,0,\ldots,i-1$ of sequence $\phi$. The cyclic-indexed partition function matrices on $\phi$ are defined in terms of standard-indexed matrices on $\phi'$ as follows:

$$\hat{R}^{\varnothing}_{j,i}(\phi) \coloneqq \hat{Q}^{\varnothing}_{j,N+i}(\phi'), \tag{S63}$$
$$\hat{R}^{e}_{j,i}(\phi) \coloneqq \hat{Q}^{e}_{j,N+i}(\phi'). \tag{S64}$$

$\hat{Q}^{\varnothing}$ and $\hat{Q}^e$ are available for the pink elements in Figure S4a from a precursor partition function calculation using the forward sweep operation order of Figure S2b. $\hat{R}^{\varnothing}$ and $\hat{R}^e$ are calculated for the blue elements in Figure S4a using the forward sweep operation order of Figure S4c.

In production code, we store $\hat{R}^{\varnothing}$ and $\hat{R}^e$ in a single $N \times N$ matrix $\hat{R}$, with $\hat{R}^{\varnothing}$ stored below the diagonal (as for the blue elements in Figure S4a) and $(\hat{R}^e)^\top$ stored on and above diagonal:

$$\hat{R}_{i,j}(\phi) := \begin{cases} \hat{R}^{\varnothing}_{i,j}(\phi), & i > j, \\ (\hat{R}^{e}_{j,i}(\phi))^\top, & i \le j. \end{cases} \tag{S65}$$

With this convention, (S62) may be restated for $i \le j$:

$$\overline{P}_{i,j}(\phi) = \mathrm{Tr}_S((\hat{Q}_{j+1,i}(\phi))^\top \hat{R}_{j+1,i}(\phi))/\overline{Q}(\phi). \tag{S66}$$

We exploit the symmetry of the pair probability matrix by calculating $\overline{P}_{i,j}(\phi)$ for $i \le j$ and setting $\overline{P}_{j,i}(\phi) = \overline{P}_{i,j}(\phi)$ for $i > j$. Pseudocode for calculation of the equilibrium pair probability matrix is shown in Algorithm S10.

Note that the use of transfer matrices $V$ and $B$ makes it possible to prohibit base $i$ from being unpaired ($V_i = 0$) or to prohibit base pair $i \cdot j$ from forming ($B_{i,j} = 0$). The combination of such manipulations enables an explicit constraint that base pair $i \cdot j$ forms, which is convenient during hierarchical ensemble decomposition in sequence design,[77,82] enabling the definition of conditional subensembles on either side of a clamped base pair $i \cdot j$.

**Algorithm S10** Backtrack-free $O(N^3)$ algorithm to calculate the equilibrium pair probability matrix $\overline{P}(\phi)$.

```
function ComputeExternalElements(Q̂, N, V, B)
    for i ∈ [0:N − 1] do
        for all j ∈ [i:N − 1] do
            if mod(j + 1, N) ≤ mod(i − 1, N) then                ▷ Cyclic indexing
                z ← Q̂
            else
                z ← R̂
            R̂_{i,j} ← χ(V, B, j, N + i, z)                    ▷ Extend operation using χ of Algorithm S3
        for all j ∈ [i + 1:N − 1] do
            R̂_{j,i} ← Σ_{k∈[j:N−1]} Q̂_{j,k}(R̂_{i,k})^⊤ + Σ_{k∈[0:i−1]} R̂_{j,k}Q̂^⊤_{i,k}     ▷ Combine operation
    return R̂

function PairProbabilities(Q̂, N, V, B)
    R̂ ← ComputeExternalElements(Q̂, N, V, B)
    for all j ∈ [0:N − 1] do        ▷ Loop over matrix columns including base case j = 0; 3′-most base is j − 1
        for all i ∈ [0:j] do                                  ▷ Loop over 5′-most base i
            P̄_{i,j} ← Tr_S((Q̂_{j+1,i})^⊤ R̂_{j+1,i})/Tr_S(Q̂_{0,N})
            P̄_{j,i} ← P̄_{i,j}                                 ▷ Symmetric probability matrix
    return P̄
```

#### S6.5 Generic approach to backtracking operation orders for structure generation

As described in Fornace et al.,[19] after calculating the complex partition function $\overline{Q}(\phi)$ with a forward sweep operation order, a structure $s_{\text{sample}}$ can be Boltzmann-sampled from the structural ensemble $\overline{\Gamma}(\phi)$ by backtracking through the matrix $\hat{Q}$ of subsequence partition function elements. Likewise, after calculating the free energy (S14) of the MFE subensemble state with a forward sweep, the corresponding MFE subensemble state $s^{\shortparallel}_{\text{MFE}} \in \overline{\Gamma}^{\shortparallel}(\phi)$ can be determined by backtracking through the matrix of subsequence MFE elements, which in turn determines the corresponding MFE proxy structure (S15).

For either structure sampling or MFE proxy structure determination, backtracking starts from the recursion for $\hat{Q}_{0,N}$, representing partition function information in the case of structure sampling and MFE information in the case of MFE proxy structure determination. In either case, backtracking is used to "choose" between competing recursion elements when a $\oplus$ operator is encountered and to "join" compatible recursions elements when a $\otimes$ operator is encountered; the mathematical implementations of these operators are described by quantity-specific evaluation algebras (see Section S5.2). For sampling, $\oplus$ corresponds to randomly choosing between competing (Boltzmann-weighted) recursion elements, while for MFE proxy structure determination, $\oplus$ corresponds to choosing the MFE of competing recursion elements. For both structure sampling and MFE structure determination, $\otimes$ corresponds to the set union $\cup$ of compatible recursion elements.

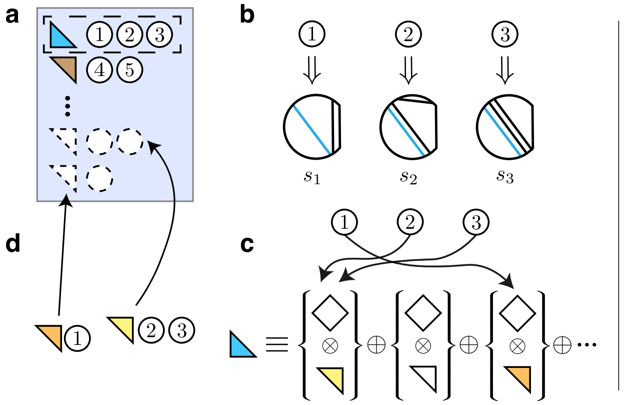

**Figure S5:** Depiction of simultaneous Boltzmann sampling algorithm. **(a)** A priority queue of remaining recursion elements with matching structure indices is maintained throughout the algorithm. Held recursion elements $E_{i,j}^{a,b}$ either satisfy $i < j$ (upper triangle; $\hat{Q}^{\varnothing}$ element) or $i > j$ (lower triangle; $\hat{Q}^{e}$ element). Until the queue is empty, a recursion element and list of structures (circled numbers) are popped from the list (blue lower triangle). **(b)** In this case, $i > j$ indicates that the derived structure contains base pair $j \cdot i-1$ or unpaired base $j$ (in the special case $i = j+1$). That base pair or unpaired base is assigned in all matching structures (here, $s_1, s_2, s_3$). **(c)** Since $i > j$, $\hat{Q}_{i,j}^{a,b}$ corresponds to a $\hat{Q}^{e}$ element ("extend" operation) formed from a sum over products of pairs comprising a diamond (corresponding to a $B$ matrix containing model parameters) and an upper triangle (a $\hat{Q}^{\varnothing}$ element). Each structure is randomly assigned to a given diamond/upper triangle pair with probabilities proportional to the weight of each pair. Here, structures $s_2$ and $s_3$ are assigned to the yellow triangle, while $s_1$ is assigned to the orange triangle. **(d)** For each chosen recursion element, if $i \neq j$, it is added to the queue with the list of matching structure(s); if the recursion element is already in the queue, the structures are appended to the respective list. **(e)** In other cases, the popped element satisfies $i < j$ (brown upper triangle). **(f)** In such cases, $\hat{Q}_{i,j}^{a,b}$ corresponds to a $\hat{Q}^{\varnothing}$ element ("combine" operation) formed from a sum over products of pairs comprising an upper triangle ($\hat{Q}^{\varnothing}$ element) and a lower triangle ($\hat{Q}^{e}$ element). Each structure is randomly assigned to a given upper triangle/lower triangle pair with probabilities proportional to the weight of each pair. Here, structure $s_4$ is assigned to the pink and gray triangles, while $s_5$ is assigned to the green and yellow triangles. **(g)** Each chosen recursion element is added to the priority queue with the matched structure(s); if the recursion element is already in the queue, the structures are appended to the respective list.

While the pair probability matrix $\overline{P}(\phi)$ provides the equilibrium probability of each base pair over the complex ensemble, it does not reveal correlation information between different base pairs. By sampling a set of $J$ secondary structures and averaging or clustering over this set, it is possible to address questions like "what is the probability that a set of adjacent bases are simultaneously unpaired?"[12] or "is the free energy landscape dominated by multiple deep basins each defined by a set of related secondary structures?"[11] Motivated by the central use case where a set of $J$ structures is needed for averaging or clustering, we develop a simultaneous sampling approach that samples $J$ structures all at once ($O(JN^2)$ with exact treatment of interior loops).[19] A given recursion element may contribute to a large number of sampled structures (e.g., if there is a deep well on the free energy landscape), so we perform backtracking using a priority queue data structure that reduces computational effort by ensuring that all samples of any given recursion element are performed during a single visit to that recursion element (see Section S6.6).

The MFE proxy structure algorithm can be generalized to calculate the set of suboptimal proxy structures $\overline{\Gamma}(\phi; \Delta G_{\text{gap}})$ within a specified free energy gap $\Delta G_{\text{gap}} \geq 0$ of the MFE subensemble state using generalized evaluation operators for $\oplus$ and $\otimes$ (see Section S5.2.6). In practice, we implement this more general algorithm and then apply it with $\Delta G_{\text{gap}} = 0$ if the MFE proxy structure is requested. The number of suboptimal proxy structures can grow rapidly with $\Delta G_{\text{gap}}$ and $N$ so we perform backtracking using a stack data structure that reduces memory usage by generating complete structures at the earliest opportunity, enabling these structures to be emitted in a streaming fashion while additional structures are determined (see Section S6.7).

#### S6.6 Operation order for Boltzmann-sampled secondary structure generation

The goal of simultaneous Boltzmann sampling is to generate $J$ secondary structures from the Boltzmann ensemble:

$$\overline{\Gamma}_{\text{sample}}(\phi; J) \in \overline{\Gamma}(\phi), \tag{S67}$$

each independently sampled with probability $\overline{p}(\phi, s)$ given by (S12).

Fig. S5 depicts the operation order for simultaneous Boltzmann sampling of $J$ structures, while Algorithm S11 provides pseudocode. A priority queue of recursion elements is maintained throughout the algorithm. The operation order works via a backtracking principle, in which, after running a forward pass to compute the partition function value, recursions are randomly sampled and inspected one by one to find the associated base pairs. In this backtracking algorithm, $\hat{Q}_{i,j}^{a,b}$ is the (scalar) partition function of a given subsequence and subensemble state, while $E_{i,j}^{a,b}$ is the recursion element which "names" that quantity, defined as the tuple $E_{i,j}^{a,b} := (i, j, a, b)$. The priority queue maps from each recursion element $E_{i,j}^{a,b}$ to the indices of all sampled structures containing that recursion element; the queue is defined via a partial ordering on recursion elements using comparison key $\delta(j \geq i) - 2|i - j|$. Such an order means that the popped element is the one with the largest extent $|i - j|$ and with elements with $i \leq j$ coming after those with $i > j$ in the case of ties. This ensures that each recursion element is only processed once.

The priority queue is initialized by considering the complex partition function $\overline{Q}(\phi) = \mathrm{Tr}_S[\hat{Q}_{0,N}]$. Since

$$\mathrm{Tr}_S[\hat{Q}_{0,N}] = \sum_{a,b} S^{b,a} \hat{Q}_{0,N}^{a,b}, \tag{S68}$$

initialization takes place by sampling recursion elements $\{E_{0,N}^{a,b}\}$ with probability proportional to $S^{b,a}\hat{Q}_{0,N}^{a,b}$ for each requested structure index, and the backtracking procedure is repeated until the queue is empty. The main speedup in the simultaneous sampling algorithm is that each recursion must be backtracked at most once (see Figure S5cf), while each might be backtracked multiple times in repeated sequential sampling.

In Algorithm S11, SampleWeights is an $O(n)$ routine for generating $n$ pseudorandom numbers uniformly from interval [0:$T$]. In contrast to sampling algorithm of Fornace et al.,[19] which sampled $n$ numbers followed by an $O(n \log n)$ sort, here we derive an $O(n)$ routine by exploiting the joint probability distribution of the maximum value of $n$ uniformly sampled numbers in [0:1], which follows $p(x_1, \cdots, x_n) = \prod_{i=1}^{n-1} \delta(x_i \leq x_{i+1})$. The maximum value $x_n$ thus follows the probability distribution $p(x_n) = \int_0^1 \cdots \int_0^1 dx_1 \cdots dx_{n-1} = n x_n^{n-1}$. Inverting the cumulative probability distribution yields that $x_n$ may be generated according to $y^{1/n} : y \sim \mathcal{U}(0,1)$ where $\mathcal{U}$ being the uniform distribution. The procedure is then iterated until no elements remain to be generated. Results are multiplied by a constant factor of $T$ to sample from [0:$T$] instead of [0:1].

When an element $E_{i,j}^{a,b}$ is popped, the condition $j = i - 1$ indicates that base $j$ is unpaired in the corresponding structures, while the condition $j < i - 1$ indicates that bases $j$ and $i - 1$ are paired in the corresponding structures. In this context, it is therefore helpful to interpret an unpaired base to be a base that (notationally speaking) "pairs to itself." As each possible contribution to the popped element is encountered, the result is accumulated in a sum variable $t$. Contributions are matched to each associated structure of index $i$ by the condition that $t \leq w_i \leq t'$ (i.e., the prefix sum crosses $w$ with the addition of that contribution). This matching may be efficiently performed in simultaneous sampling: since the structure weights $\vec{w}$ are pre-sorted, the position of the new prefix sum $t$ in $\vec{w}$ may be found via binary search with a cost $O(\log(|\vec{w}|))$. Structure weights which have been previously matched may be discarded from future consideration; when there are no remaining structure weights, the procedure may terminate early. When a structure weight is matched, the contributing recursion element is inserted into the priority queue $\mathcal{M}$, and the structure index is appended to the mapped item for that element. During backtracking of a recursion for a $\hat{Q}^{\varnothing}$ element, we search 5′-most opening base pair positions $k$ in descending order. While the order does not affect correctness, local base pairs are common in many sequences, in which performing the scan in this order will short-circuit sooner without considering all possibilities.

To Boltzmann-sample a single secondary structure, the time complexity is $O(N^2\gamma + N\xi\gamma^2)$; there are a maximum of $N$ upper triangle ($\hat{Q}^{\varnothing}$) recursion elements to backtrack, each with a linear scan over $O(N\gamma)$ elements, and a maximum of $N$ lower triangle ($\hat{Q}^{e}$) recursion elements to backtrack, each with a scan of $O(\xi\gamma^2)$ elements. Note that summation over product $SB$ can be pre-contracted in Algorithm S11 (see Comment) so that it does not impact the algorithm complexity. Since the simultaneous algorithm is strictly faster than repeated sequential sampling, the time complexity with respect to complex size $N$ is thus upper bounded by $O(JN^2)$ for $J$ samples. In practice, as with our previous sampling algorithm,[19] the observed empirical complexity is expected to be lower.

**Algorithm S11** Simultaneous Boltzmann sampling

**function** SampleWeights($\mathcal{I}, T$)

▷ *Associate random weights in [0:T] to each element in index sequence $\mathcal{I}$ and return the resultant pairs in the order in which those weights are ascending* ◁

$\mathcal{I}' \leftarrow$ UniformlyShuffled($\mathcal{I}$)

$T' \leftarrow T$

**for** $n = |\mathcal{I}|, \ldots, 1$ **do**

$T' \leftarrow T' \cdot (\text{UniformRandom}(0, 1))^{1/n}$

$w_{n-1} \leftarrow (T', \mathcal{I}'_{n-1})$ ▷ *Each returned element has a weight and an index*

**return** $\vec{w}$

**function** AdjustWeights($\vec{w}, t, \mathcal{M}, \vec{e}$)

▷ *Consider prefix sum $t$ formed from recursion elements $\vec{e}$, modify $\vec{w}$ and $\mathcal{M}$ accordingly* ◁

**if** $|\vec{w}| > 0$ and FirstWeight($\vec{w}$) $< t$ **then**

$(\vec{u}, \vec{w}) \leftarrow$ UpperBound($\vec{w}, t$) ▷ *Partition elements (already sorted) with weights $\leq t$ and $> t$*

**for all** $E^{a,b}_{i,j} \in \vec{e}$ **do**

▷ *Insert recursion element into $\mathcal{M}$ if absent, concatenate indices of $\vec{u}$ to its mapped index list* ◁

InsertAndConcatenate($\mathcal{M}, E^{a,b}_{i,j}, \vec{u}$)

**return** $|\vec{w}| = 0$ ▷ *Return whether backtracking of this element is finished*

**procedure** SampleElement($t, \mathcal{M}, \hat{Q}, E^{a,b}_{i,j}, \vec{w}$) ▷ *Backtrack through element $E^{a,b}_{i,j}$ with weights $\vec{w}$*

**if** $i < j$ **then** ▷ *Backtrack through "combine" recursion corresponding to $\phi_{[i:j-1]}$*

**for all** $k = j-1, \ldots, i,\ c \in [0{:}\gamma-1]$ **do** ▷ *Descending order of $k$ is often more efficient (see text for details)*

$t \leftarrow t + \hat{Q}^{a,c}_{i,k} \cdot \hat{Q}^{b,c}_{j,k}$

**if** AdjustWeights($\vec{w}, t, \mathcal{M}, \{E^{a,c}_{i,k}, E^{b,c}_{j,k}\}$) **then return**

**else** ▷ *Backtrack through "extend" recursion corresponding to $\phi_{[j:i-1]}$ with base pair $j \cdot i-1$*

**for all** $\rho \in [1{:}\xi-1],\ c \in [0{:}\gamma-1],\ d \in [0{:}\gamma-1]$ **do**

$t \leftarrow t + \sum_{e \in [0:\gamma-1]} B^{\rho,a,b}_{i,j} S^{d,e} B^{\rho,e,c}_{j,i} \hat{Q}^{c,d}_{j+1,i-1}$ ▷ *Precontraction possible for sum over SB product*

**if** AdjustWeights($\vec{w}, t, \mathcal{M}, \{E^{c,d}_{j+1,i-1}\}$) **then return**

**function** Sample($J, \hat{Q}, N$) ▷ *$J$: the number of samples desired*

$\mathcal{M} \leftarrow$ EmptyPriorityQueue() ▷ *Map from recursion element to the indices of associated structures*

**for all** $k \in [0{:}J-1]$ **do**

$s_k \leftarrow$ UnpairedStructure($N$)

▷ *Sample from the unconstrained partition function $\text{Tr}_S[\hat{Q}_{0,N}]$* ◁

$w \leftarrow$ SampleWeights($[0{:}J-1], \text{Tr}_S[\hat{Q}_{0,N}]$)

$t \leftarrow 0$

**for all** $a \in [0{:}\gamma-1],\ b \in [0{:}\gamma-1]$ **do**

$t \leftarrow t + S_{b,a} \hat{Q}^{a,b}_{0,N}$

**if** AdjustWeights($\vec{w}, t, \mathcal{M}, \{E^{a,b}_{0,N}\}$) **then break**

▷ *Sampling remaining elements until none remain* ◁

**while** $|\mathcal{M}| > 0$ **do**

$(E^{a,b}_{i,j}, \mathcal{I}) \leftarrow$ Pop($\mathcal{M}$)

**if** $i > j+1$ **then**

**for all** $k \in \mathcal{I}$ **do** ▷ *Add base pairs in all matching structures $\mathcal{I}$*

AddBasePair($s_k, j, i-1$)

**if** $j > i+1$ or $i > j+3$ **then** ▷ *Ignore intervals with no possible nested pairs*

SampleElement($0, \mathcal{M}, \hat{Q}, E^{a,b}_{i,j}$, SampleWeights($\mathcal{I}, \hat{Q}^{a,b}_{i,j}$))

**return** $\vec{s}$

#### S6.7 Operation orders for MFE and suboptimal proxy structure generation

As a starting point, Figure S6 outlines an operation order to return the first-found MFE subensemble state. Note that there can be multiple MFE subensemble states, so it is simplest to first consider an algorithm that returns only one of them. The operation order works via a backtracking principle, in which, after running a forward pass to compute the free energy of the MFE subensemble state given by (S14), recursions are inspected one by one to find the base pairs associated with that MFE value. In this backtracking algorithm, we adopt the notation that $\hat{Q}^{a,b}_{i,j}$ is the (scalar) MFE of a given subsequence and subensemble state, while $E^{a,b}_{i,j}$ is the recursion element which "names" that quantity, defined as the tuple $E^{a,b}_{i,j} \coloneqq (i, j, a, b)$.

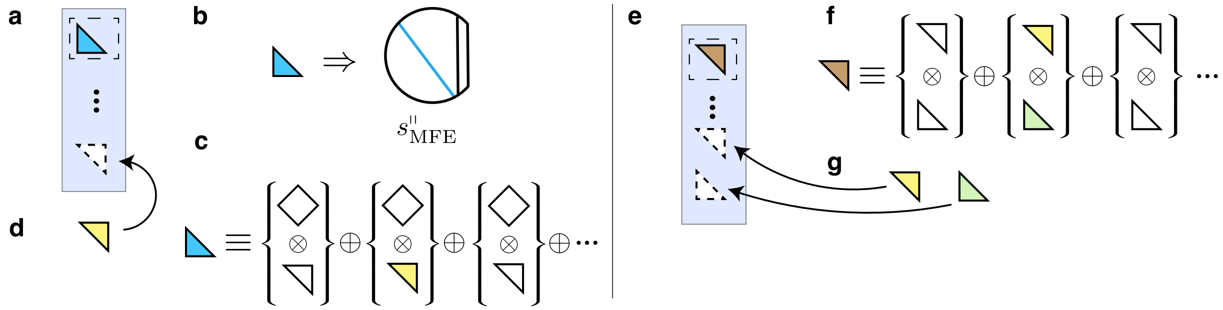

**Figure S6:** Depiction of first-found MFE subensemble state generation algorithm. **(a)** A set of remaining recursion elements is maintained throughout the algorithm. Held recursion elements $E^{a,b}_{i,j}$ either satisfy $i < j$ (upper triangle; $\hat{Q}^{\varnothing}$ element) or $i > j$ (lower triangle; $\hat{Q}^{e}$ element). The set of recursion elements is initialized to $\{E^{a,b}_{0,N}\}$ with $a, b = \arg\min_{a',b'}(\hat{Q}^{a',b'}_{0,N} + S^{b',a'})$. Until the set is empty, a recursion element is popped from the set (blue lower triangle). **(b)** In this case, $i > j$ indicates that the derived structure contains base pair $j \cdot i-1$ or unpaired base $j$ (in the special case $i = j+1$). That base pair or unpaired base is assigned in the output structure $s_{\mathrm{MFE}'}$. **(c)** Since $i > j$, $\hat{Q}^{a,b}_{i,j}$ corresponds to a $\hat{Q}^{e}$ element ("extend" operation) formed from a minimum over sums of pairs comprising a diamond (corresponding to a $B$ matrix containing model parameters) and an upper triangle (a $\hat{Q}^{\varnothing}$ element). The first such pair $E^{c,d}_{j+1,i-1}$ to match the minimum value $\hat{Q}^{a,b}_{i,j}$ is chosen (yellow triangle; the diamond may be ignored, as it corresponds to a $V$, $B$, or $S$ parameter that need not be backtracked). **(d)** The chosen upper triangle is added to the set of remaining elements (unless $i = j$, the empty recursion). **(e)** In other cases, the popped element satisfies $i < j$ (brown upper triangle). **(f)** In such cases, the value $\hat{Q}^{a,b}_{i,j}$ corresponds to a $\hat{Q}^{\varnothing}$ element ("combine" operation)formed from a minimum over sums of pairs comprising an upper triangle ($\hat{Q}^{\varnothing}$ element) and a lower triangle ($\hat{Q}^{e}$ element). The first such pair to match the minimum value $\hat{Q}^{a,b}_{i,j}$ is chosen (yellow and green triangles). **(g)** Each of these two chosen triangles $E^{a,c}_{i,k}$ (yellow) and $E^{a,c}_{j,k}$ (green) is added to the set of remaining recursion elements.

The suboptimal structure generation algorithm returns the set of all suboptimal proxy structures, $\overline{\Gamma}_{\mathrm{subopt}}(\phi; \Delta G_{\mathrm{gap}})$, given by (S18) that have subensemble states within a specified $\Delta G_{\mathrm{gap}} \geq 0$ of the MFE subensemble state. Our algorithm is described in terms of dimensionless energy gap, $g_{\mathrm{gap}}$, where $\Delta G_{\mathrm{gap}} \coloneqq g_{\mathrm{gap}} k_B T$.

Efficient suboptimal structure generation follows a related approach to the one for Boltzmann sampling. Our implementation closely follows Fornace et al.[19] with modifications due to the use of new recursions and the matrix formulation. Another change with the current algorithm is the explicit handling of the first-found MFE stacking state in each backtracking iteration; with this approach, the prior map from partial structures to unrecursed recursion elements is replaced by a simple list (rather than ordered map) for each structure of the component base pairs and respective unrecursed recursion elements. During the backtracking of a single recursion element, each matching structure is modified in place to reflect the first-found MFE contribution, while every other matching contribution is handled by appending a new partial structure. This modification makes it trivial to modify the algorithm to return solely the first-found MFE stacking state.

Figure S7 summarizes the suboptimal structure generation operation order. Building on the approach for simultaneous Boltzmann sampling (Algorithm S5), both a priority queue of recursion elements with matching structure indices and a reverse map from structure indices to their matched recursion elements are maintained throughout the algorithm. Again, the operation order works via a backtracking principle, in which, after running a forward pass to compute the minimum free energy value, recursions are inspected one by one to identify all combinations of recursion elements compatible with each output structure. The priority queue and map are initialized by associating a unique structure index to each recursion element $E^{a,b}_{0,N}$ satisfying $S^{b,a}\hat{Q}^{a,b}_{0,N} \leq \overline{g}_{\mathrm{MFE}}(\phi) + g_{\mathrm{gap}}$, and the backtracking

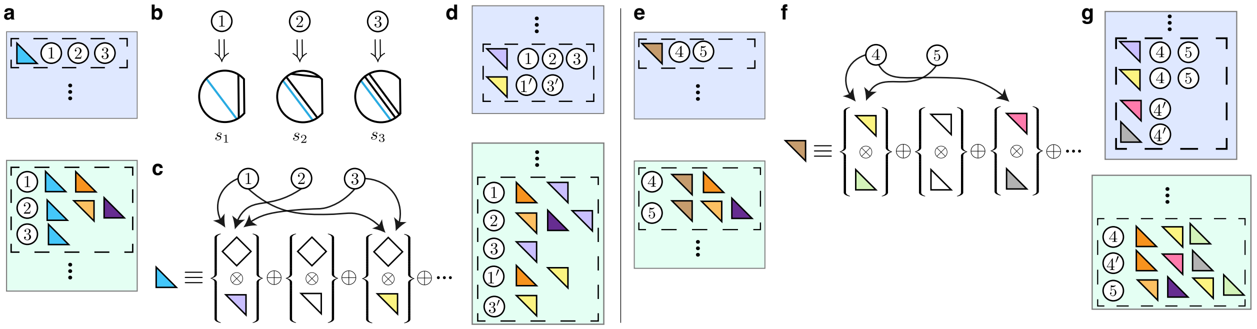

**Figure S7:** Depiction of suboptimal structure generation algorithm. **(a)** A priority queue of remaining recursion elements with matching structure indices is maintained throughout the algorithm (blue), as is a map from each structure to its matching remaining recursion elements (green). Held recursion elements $E^{a,b}_{i,j}$ either satisfy $i < j$ (upper triangle; $\hat{Q}^{\varnothing}$ element) or $i > j$ (lower triangle; $\hat{Q}^{e}$ element). Until the queue is empty, a recursion element and list of structures (circled numbers) popped from the list (blue lower triangle). **(b)** In this case, $i > j$ indicates that the derived structure contains base pair $j \cdot i-1$ or unpaired base $j$ (in the special case $i = j+1$). That base pair or unpaired base is assigned in all matching structures (here, $s_1, s_2, s_3$). **(c)** Since $i > j$, $\hat{Q}^{a,b}_{i,j}$ corresponds to a $\hat{Q}^{e}$ element ("extend" operation) formed from a minimum over sums of pairs comprising a diamond (corresponding to a $B$ matrix containing model parameters) and an upper triangle (a $\hat{Q}^{\varnothing}$ element). Each structure is assigned to every recursion element incurring an energy cost such that the modified energy of that structure is still below the user-chosen energy gap. Here, structure $s_2$ is assigned to the purple triangle, while $s_1$ and $s_3$ are assigned to both the purple and yellow triangles. A structure is duplicated for each additional recursion element beyond the first matched one (yielding $s_{1'}$ and $s_{3'}$ here). **(d)** For each chosen recursion element, if $i \neq j$, it is added to the queue (blue) with the list of matching structure(s); if the recursion element is already in the queue, the structures are appended to the respective list. Each structure is also added back into the map (green) with a list of its matching recursion elements. **(e)** In other cases, the popped element satisfies $i < j$ (brown upper triangle). **(f)** In such cases, $Q^{a,b}_{i,j}$ corresponds to a $\hat{Q}^{\varnothing}$ element ("combine" operation) formed from a minimum over sums of pairs of comprising an upper triangle ($\hat{Q}^{\varnothing}$ element) and a lower triange ($\hat{Q}^{e}$ element). Each structure is assigned to every recursion element incurring an energy cost such that the modified energy of that structure is still below the user-chosen energy gap. Here, structure $s_4$ is assigned to the green, yellow, pink, and gray triangles, while $s_5$ is assigned to the green and yellow triangles. Again, a structure is duplicated for each matching recursion element beyond the first one (yielding $s_{4'}$ here). **(g)** Again, each chosen recursion element is added to the priority queue (blue) with the matched structure(s), and each structure is added to the map (green) with a list of its matching recursion elements.

procedure is repeated until the queue is empty. Again, each recursion must be backtracked at most once (Figure S7cf), providing empirical cost savings.

For a single recursion element, backtracking works by the following procedure. For each possible alternative substructure to the given recursion element: 1) if that substructure is the first-found MFE substructure, save the respective recursion indices in $\vec{e}_{\text{MFE}}$, or 2) for every combination of that substructure and partial structure that can support a free energy within the specified user gap, make a new output structure slot by a) copying the respective partial structure and b) adding the new substructure recursion elements. At all times in the algorithm, we maintain: 1) a map $\mathcal{M}$ from recursion elements to the indices of matching partial structures $\mathcal{I}$, and 2) a list $\vec{s}$ for each partial structure containing a) the base pairs in $s$, b) the free energy difference between $s$ and the MFE subensemble state, and c) the unrecursed recursion elements discovered for $s$.

Algorithm S12 employs helper functions. AppendStructureIndices takes the recursion elements for the best structure, then adds those elements to that structure and that structure to those elements. LowerBound uses binary search to return all structure indices from input $\mathcal{I}$ for which the given contribution may be allowed while not exceeding the user-specified free energy gap. InsertAndAppend($\mathcal{M}, h, \iota$): 1) inserts a recursion element $h$ into queue $\mathcal{M}$ if it does not exist, and 2) appends structure index $\iota$ to the mapped structure indices for that element. Finally, AppendSegments($s, \vec{e}$) adds each of the recursion elements in $\vec{e}$ to the structure $s$.

It is difficult to characterize the time complexity of suboptimal structure generation for the general case of an arbitrary free energy gap and sequence as the number of generated structures is potentially exponential in $N$. In the special case with $g_{\text{gap}} = 0$ for a complex with a unique $s^{\shortparallel}_{\text{MFE}}$, the time complexity of Algorithm S12 is $O(N^2\gamma + N\xi\gamma^2)$, as in the case of sampling a single structure (Section S6.6). Note that the minimum over sum $S + B$ can be pre-contracted in Algorithm S12 (see Comment) so that it does not impact the algorithm complexity.

**Algorithm S12** Suboptimal structure generation

```
procedure Adjust(e⃗_MFE, M, s⃗, g_gap, t, T, I, e⃗)
    ▷ Arguments: first-discovered optimal structure, queue, structure list, user gap, alternative value, optimal value, structure indices, recursion indices ◁
    if t = T and e⃗_MFE = ∅ then                ▷ Save the first-discovered optimal structure for later handling
        e⃗_MFE ← e⃗
    else if t + g_gap < T then                  ▷ Create new partial structures for every other feasible combination
        for i ∈ [0:LowerBound(I, g_gap + t − T) − 1] do
            (r, d, y⃗) ← s_{I_i}                  ▷ Structure, structure gap, recursion indices
            AppendStructure(s⃗, (r, d + T − t, e⃗ ∪ y⃗))   ▷ Create a new structure combining s_{I_i} with e⃗
            for h ∈ e⃗ ∪ y⃗ do
                InsertAndAppend(M, h, |s⃗| − 1)    ▷ Update M with the new partial structure

procedure SuboptElement(M, s⃗, E^{a,b}_{i,j}, I)
    e⃗_MFE ← ∅                                   ▷ Backtrack through contributions to a given element
    if i < j then
        for all k ∈ [i:j − 1], c ∈ [0:γ − 1] do
            Adjust(e⃗_MFE, M, s⃗, g_gap, Q̂^{a,c}_{i,k} + Q̂^{b,c}_{j,k}, Q̂^{a,b}_{i,j}, I, {E^{a,c}_{i,k}, E^{b,c}_{j,k}})
    else
        for all ρ ∈ [1:ξ − 1], c ∈ [0:γ − 1], d ∈ [0:γ − 1] do
            t ← min_{e∈[0:γ−1]} B^{ρ,a,b} + S^{d,e} + B^{ρ,e,c} + Q̂^{c,d}_{j+1,i−1}   ▷ Precontraction possible for minimum over sum S + B
            Adjust(e⃗_MFE, M, s⃗, g_gap, t, Q̂^{a,b}_{i,j}, I, {E^{c,d}_{j+1,i−1}})
    for all b ∈ e⃗_MFE do                         ▷ Handle the first-found optimal structure separately
        AppendStructureIndices(M, b, I)
    for all k ∈ I do
        AppendSegments(s_k, e⃗_MFE)

function Subopt(g_gap, Q̂, N)
    M ← EmptyOrderedMap()                       ▷ Map from recursion elements to associated structure indices
    s⃗ ← {(UnpairedStructure(N), 0, ∅)}          ▷ Initialize partial structures with a single empty structure
    ▷ Recurse through the trace Tr_S[Q̂_{0,N}] ◁
    e⃗_MFE ← ∅                                   ▷ Recursion indices of the first-discovered optimal structure
    for all a ∈ [0:γ − 1], b ∈ [0:γ − 1] do
        Adjust(e⃗_MFE, M, s⃗, g_gap, S^{b,a} + Q̂^{a,b}_{0,N}, Tr_S[Q̂_{0,N}], {0}, {E^{a,b}_{0,N}})
    for all b ∈ e⃗_MFE do
        AppendStructureIndices(M, b, I)
    AppendSegments(s_0, e⃗_MFE)
    ▷ Backtrack through recursion elements until none remain ◁
    while |M| > 0 do
        (E^{a,b}_{i,j}, I) ← Pop(M)
        if i > j + 1 then
            for all k ∈ I do                     ▷ Add base pairs in all matching structures I
                s_k ← AddBasePair(s_k, j, i − 1)
        for all k ∈ I do
            EraseElement(s_k, E^{a,b}_{i,j})      ▷ Remove recursion element from the structures which contained it
        if j > i + 1 or i > j + 3 then           ▷ Ignore intervals with no possible nested pairs
            SortByGap(I, s⃗)                     ▷ Sort structure indices by the gap of the respective structure
            SuboptElement(M, E^{a,b}_{i,j}, I)
    return s⃗
```

Note that the MFE algorithm does not return the free energy of the MFE secondary structure $s_{\text{MFE}}$ but rather the free energy of the MFE subensemble state $s^{\shortparallel}_{\text{MFE}}$ given by (S14). The backtracking dynamic program then returns the MFE proxy structure $s_{\text{MFE}'}$ given by (S15) that contains $s^{\shortparallel}_{\text{MFE}}$ within its subensemble.

By definition, an MFE secondary structure has the highest equilibrium probability, $\overline{p}(\phi, s_{\text{MFE}})$, in structural ensemble $\overline{\Gamma}(\phi)$. However, $\overline{p}(\phi, s_{\text{MFE}})$ can nonetheless be arbitrarily small due to competition from other structures in $\overline{\Gamma}(\phi)$. For ensembles where $\overline{p}(\phi, s_{\text{MFE}})$ is non-negligible, an attractive alternative to the deterministic approach is to use Boltzmann sampling to discover the MFE secondary structure.[19] One advantage of the random approach is that it determines MFE status based on secondary structure $s$ rather than subensemble state $s^{\shortparallel} \in s$.

Sampling is performed for ensemble $\overline{\Gamma}(\phi)$ treating all strands as distinct. Suppose that the identity of $s_{\text{MFE}}$ is unknown, as is its free energy $\overline{\Delta G}(\phi, s_{\text{MFE}})$ and its equilibrium probability $\overline{p}(\phi, s_{\text{MFE}})$. The probability, $p_{\text{fail}}$, that a sample of $J$ structures does not include a structure $s_{\text{MFE}}$ that has probability $\overline{p}(\phi, s_{\text{MFE}}) \geq p_{\text{min}}$ is

$$p_{\text{fail}} \leq (1 - p_{\text{min}})^J. \tag{S69}$$

Inverting this relationship, for a given $p_{\text{min}}$, we can calculate the number of samples, $J$, required to assure a failure probability no higher than $p_{\text{fail}}$:

$$J \geq \frac{\log p_{\text{fail}}}{\log(1 - p_{\text{min}})} \approx \frac{\log p_{\text{fail}}}{-p_{\text{min}}} \tag{S70}$$

Because the dependence of $J$ on $p_{\text{fail}}$ is logarithmic, it is inexpensive to reduce $p_{\text{fail}}$ for fixed $p_{\text{min}}$. For example, for $p_{\text{min}} = 0.01$, we have $J \geq 688$ for $p_{\text{fail}} = 10^{-3}$ and $J \geq 2750$ for $p_{\text{fail}} = 10^{-12}$. However, the required number of samples is sensitive to the value of $p_{\text{min}}$ (the assumed lower bound in the MFE probability). For example, holding $p_{\text{fail}} = 10^{-12}$ fixed, we require $J \geq 27,618$ samples for $p_{\text{min}} = 0.001$ and $J \geq 276,297$ samples for $p_{\text{min}} = 0.0001$. While that number of samples remains affordable using the simultaneous sampling algorithm (Algorithm S11), if the MFE probability becomes vanishingly small, the required number of samples would grow too large to be practical. On the other hand, if the MFE probability is vanishingly small, the MFE structure may not provide a useful summary of the equilibrium base-pairing properties of the ensemble (in which case the equilibrium base-pairing probability matrix $P(\phi)$ will continue to provide such a summary).

After sampling $J$ structures using the new simultaneous sampling method, let $p_{\text{MFE}^*}$ denote the highest probability of the sampled structures, and let $s_{\text{MFE}^*}$ denote the MFE proxy structure determined by random sampling. The probability that there exists an (undiscovered) MFE structure with $\overline{p}_{\text{MFE}} \geq \overline{p}_{\text{MFE}^*}$ is bounded by

$$p_{\text{fail}} \leq [1 - \overline{p}(\phi, s_{\text{MFE}^*})]^J. \tag{S71}$$

Hence, after sampling $J$ structures, it is straightforward to calculate the probability that the true MFE structure was not identified. If desired, additional samples can be performed to increase $J$ (potentially identifying a higher $p_{\text{MFE}^*}$) and further reduce the failure rate.

### S7 Logarithmic scaling of the entropic cost of loop closure

#### S7.1 Fits of logarithmic functional forms

To examine approximation of logarithmic scaling of the entropic cost of loop closure using transfer matrices, fits of $g^{\nu}_{\mathrm{unweighted}}(n)$ using an unweighted trace, or $g^{\nu}_{\mathrm{weighted}}(n)$ using a weighted trace, to the Jacobson-Stockmayer theoretical result, $g_{\mathrm{theory}}(n)$, were performed by minimizing the fitting error:

$$\min_{\vec{a},\vec{b}} \sum_{n=1}^{50} [g^{\nu}_{\mathrm{(un)weighted}}(n) - g_{\mathrm{theory}}(n)]^2, \tag{S72}$$

over loops with $n = 1, \ldots, 50$ nucleotides for models with $\nu = 1, \ldots, 5$ and $\kappa = 1.75$ using the `NonlinearModelFit` nonlinear least squares routine in Mathematica. Fitting errors are displayed in Table S2 for the fits displayed in Fig. 6 (note that the fitted curves are displayed beyond the fitting region $1 \leq n \leq 50$). A large fitting error is observed using $\nu = 1$, for which the unweighted and weighted formulations are equivalent. The fitting error decreases slowly upon the addition of more terms to the unweighted formulation, but decreases rapidly with the addition of each new term to the weighted formulation. The regressed parameters for models with $\nu = 1, \ldots, 5$ are displayed in Table S3 (unweighted trace) and Table S4 (weighted trace).

| | RMSE | |
|---|---|---|
| $\nu$ | Unweighted | Weighted |
| 1 | 0.642568 | 0.642568 |
| 2 | 0.547698 | 0.157721 |
| 3 | 0.488755 | 0.0351679 |
| 4 | 0.445939 | 0.00600311 |
| 5 | 0.412417 | 0.000798229 |

**Table S2:** Fitting errors for approximate logarithmic entropic scaling using an unweighted or weighted trace (cf. Fig. 6). Root mean square errors for (S72) for models with $\nu = 1, \ldots, 5$.

| $\nu$ | $a_0$ | $b_1$ | $b_2$ | $b_3$ | $b_4$ | $b_5$ |
|---|---|---|---|---|---|---|
| 1 | 2.72292 | −0.0970119 | | | | |
| 2 | 2.97828 | −0.290621 | −0.089947 | | | |
| 3 | 3.12939 | −0.307835 | −0.307835 | −0.0857241 | | |
| 4 | 3.23651 | −0.321259 | −0.321259 | −0.321259 | −0.0827154 | |
| 5 | 3.3191 | −0.332462 | −0.332462 | −0.332462 | −0.332462 | −0.0803884 |

**Table S3:** Regressed parameters for approximate logarithmic entropic scaling using an unweighted trace (cf. Figure 6a). Fits of $g^{\nu}_{\mathrm{unweighted}}(n)$ to $g_{\mathrm{theory}}$ for $\nu = 1, \ldots, 5$.

| $\nu$ | $a_1$ | $a_2$ | $a_3$ | $a_4$ | $a_5$ | $b_1$ | $b_2$ | $b_3$ | $b_4$ | $b_5$ |
|---|---|---|---|---|---|---|---|---|---|---|
| 1 | −2.72292 | | | | | −0.0970119 | | | | |
| 2 | −0.120923 | −3.97237 | | | | −0.533379 | −0.0615537 | | | |
| 3 | 1.02061 | −1.95306 | −4.58518 | | | −1.22731 | −0.2449 | −0.0462829 | | |
| 4 | 1.50671 | −0.833561 | −2.87381 | −5.03413 | | −1.87936 | −0.536352 | −0.162895 | −0.0367551 | |
| 5 | 1.72664 | −0.225072 | −1.87894 | −3.47942 | −5.41296 | −2.37743 | −0.860978 | −0.335985 | −0.122553 | −0.0299985 |

**Table S4:** Regressed parameters for approximate logarithmic entropic scaling using a weighted trace (cf. Figure 6b). Fits of $g^{\nu}_{\mathrm{weighted}}(n)$ to $g_{\mathrm{theory}}$ for $\nu = 1, \ldots, 5$.

#### S7.2 Scaling of entropic cost with loop size for transfer matrix models

Jacobson-Stockmayer theory states that the entropic cost of loop closure scales logarithmically with loop size (2.30). Here, we calculate the predicted entropic cost of loop closure as a function of loop size for transfer matrix models to enable comparison to the theoretical result.

Loop test sets were constructed for random RNA loop sequences with 1 closing base pair (hairpin loops), 2 closing base pairs (interior loops), and 3 closing base pairs (multiloops), representing all three categories of non-exterior loops (i.e., all categories of loops containing no nick). Within a loop, each segment of unpaired bases between closing base pairs is the same length. For example, in an interior loop or multiloop, the unpaired segments between closing base pairs are selected from $\{0, 1, 2, \dots\}$ nt. For a hairpin loop, the segment of unpaired bases is selected from $\{3, 4, 5, \dots\}$ nt (due to steric hindrance).

For a given loop, we wish to calculate the entropic cost of closure, which is obtained subtracting the entropy of a reference unpaired state (secondary structure $s_{\text{unpaired}}$), from the entropy of the closed state (secondary structure $s_{\text{closed}}$):

$$\Delta S_{\text{closure}}(\phi) = \Delta S(\phi, s_{\text{closed}}) - \Delta S(\phi, s_{\text{unpaired}}) \tag{S73}$$

These entropies are calculated by taking partial derivatives of the corresponding free energies as follows:

$$\begin{aligned}\Delta S_{\text{closure}}(\phi) &= -\frac{\partial}{\partial T}[\Delta G(\phi, s_{\text{closed}}) - \Delta G(\phi, s_{\text{unpaired}})] \\ &= k_B \left[\beta \frac{\partial}{\partial \beta} - 1\right][g(\phi, s_{\text{closed}}) - g(\phi, s_{\text{unpaired}})],\end{aligned} \tag{S74}$$

using the $\partial$SumProduct evaluation algebra to compute derivatives analytically, and evaluating the resulting expressions at reference temperature $T_{37} \coloneqq 310.15$ K.

We then compare the $-\Delta S_{\text{closure}}(\phi)$ calculated using the transfer matrix model against the theoretical entropy result

$$\Delta S_{\text{theory}}(n) = -k_B \kappa \log(n) \tag{S75}$$

with $\kappa = 1.75$ and loop size $n$ defined as the number of transfer matrices in the loop transfer matrix network (i.e., number of unpaired bases + base pairs). Note that (2.30) displays the (unitless) free energy, $g_{\text{theory}}(n) = -\Delta S_{\text{theory}}(n)/k_B$, associated with this entropic cost.

To compare modeling to theory, we consider transfer matrix models of dimension $\nu, \mu, \xi = (2, 2, 1)$ regressed from literature melt data (Section S9.1). Figure 10 displays results for RNA multiloops using transfer matrix models with either the fixed-weight or floating-weight formulation. Figures S8 (RNA) and S9 (DNA) display results for hairpin loops, interior loops, and multiloops using transfer matrix models with either the fixed-weight, floating-weight, or floating block formulations.

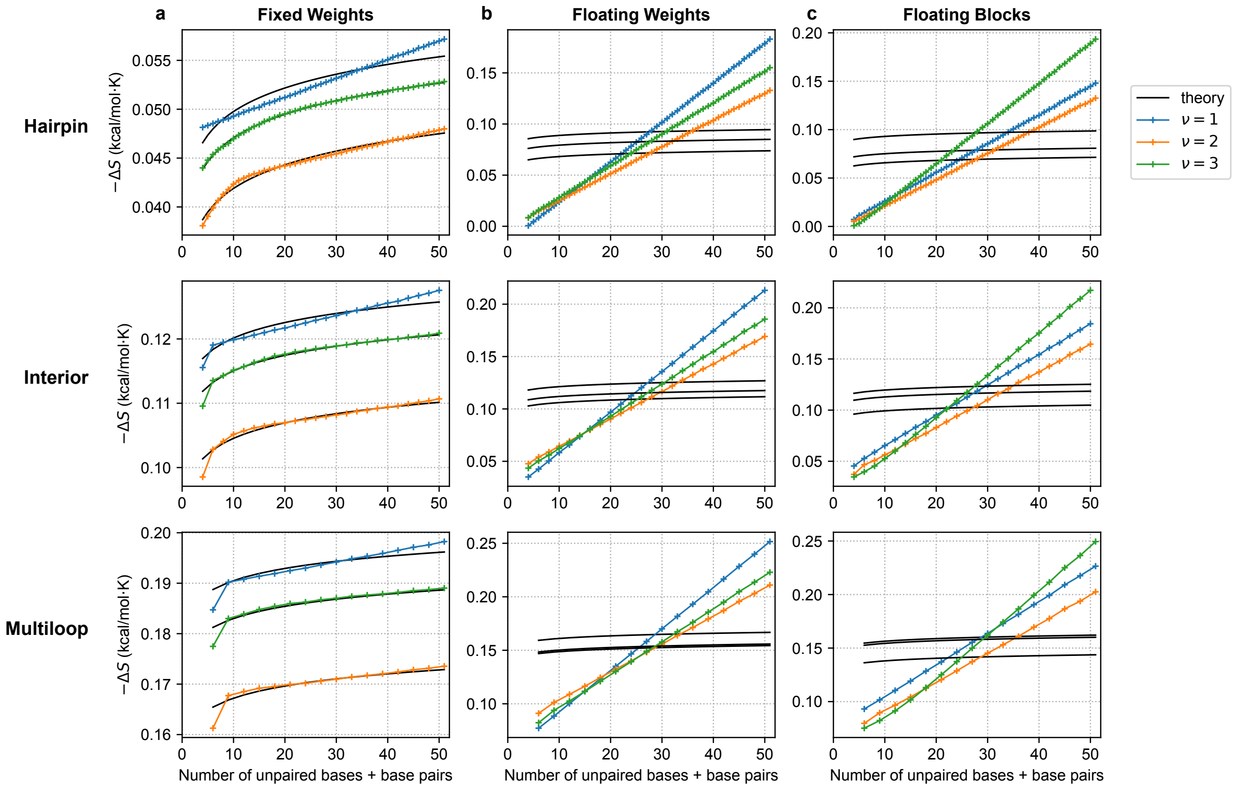

**Figure S8:** Assessment of logarithmic scaling of entropic cost with loop size for RNA multiloops using transfer matrix models (cf. Figure 10 with: a) fixed-weight formulation, b) floating-weight formulation, c) floating-block formulation. Means over 10,000 random sequences for each combination of loop type and size. Theory depicted for $\kappa = 1.75$. Fit of theory curve to transfer matrix results performed for each $\nu$ by minimizing $\min_{g_{\text{init}}} \sum_n (\Delta S^{\nu}_{\text{closure}}(n) - \Delta S_{\text{theory}}(n; g_{\text{init}}))^2$ over loop size $n$ with respect to free theory parameter $g_{\text{init}}$. Transfer matrix model: $\nu, \mu, \xi = (2, 2, 1)$. Conditions: RNA, 37 °C, 1.0 M $Na^+$.

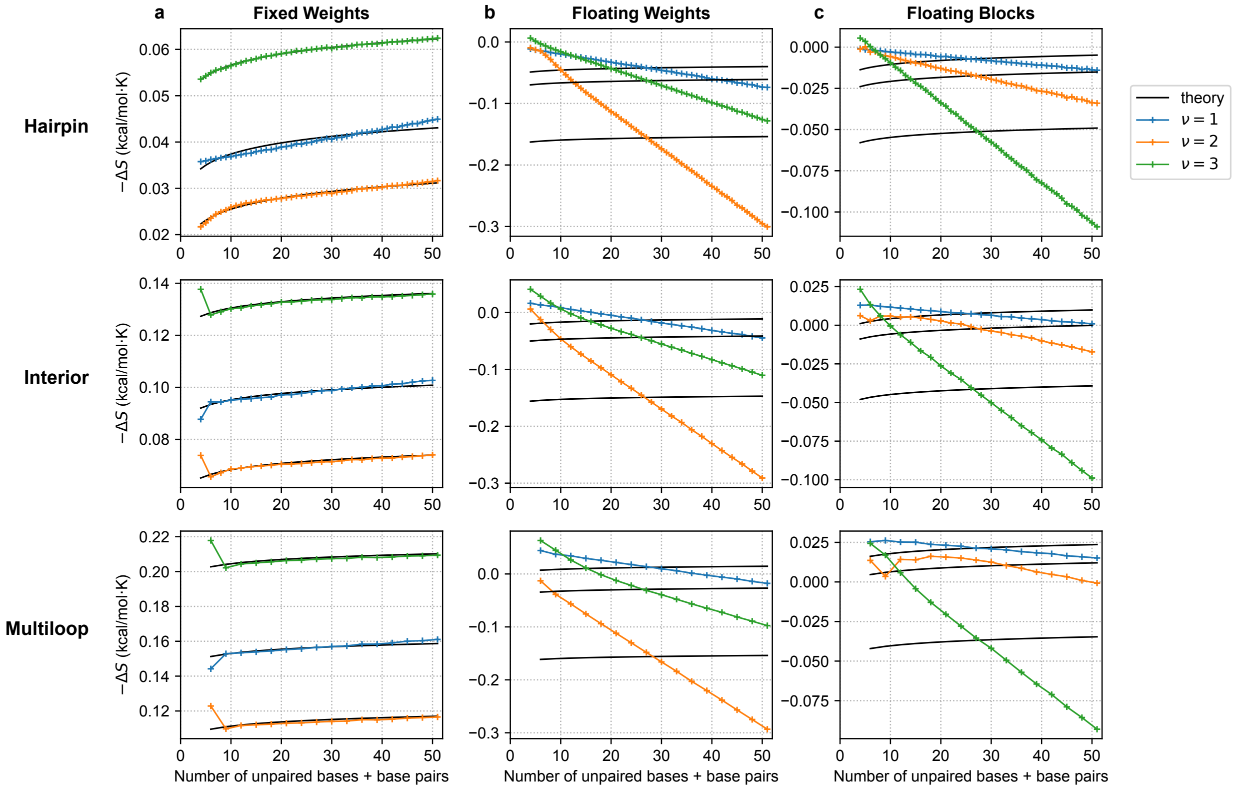

**Figure S9:** Assessment of logarithmic scaling of entropic cost with loop size for DNA multiloops using transfer matrix models (cf. Figure 10) with: a) fixed-weight formulation, b) floating-weight formulation, c) floating-block formulation. Means over 10,000 random sequences for each combination of loop type and size. Theory depicted for $\kappa = 1.75$. Fit of theory curve to transfer matrix results performed for each $\nu$ by minimizing $\min_{g_{\text{init}}} \sum_n (\Delta S^{\nu}_{\text{closure}}(n) - \Delta S_{\text{theory}}(n; g_{\text{init}}))^2$ over loop size $n$ with respect to free theory parameter $g_{\text{init}}$. Transfer matrix model: $\nu, \mu, \xi = (2, 2, 1)$. Conditions: DNA, 37 °C, 1 M $Na^+$.

### S8 Representational capacity of transfer matrix models to reproduce reference NN free energy models

#### S8.1 The box ensemble

To provide a basis for measuring the representational capacity of transfer matrix models to reproduce reference NN free energy models, we consider the ensemble of a fixed volume (or "box") containing an arbitrary number of strand species that interact to form complexes in a dilute solution. Here, we define the *box ensemble* by adapting the formulation of Dirks et al.[13]

Consider a box containing $M_{\text{sol}}$ solvent molecules and the set of strands $\Psi^0$, where each strand is distinct. Suppose the strands can interact to form the set of complexes $\Psi$. For each complex $j \in \Psi$, let $\phi_j$ denote the sequence and $Q(\phi_j)$ the partition function. In general, due to conservation of mass, the strands in $\Psi^0$ can interact to form only a subset of the complexes in $\Psi$ at any given time, with each such subset corresponding to a *complex composition* of the box. For example, if $\Psi^0 = \{A, B\}$ and $\Psi = \{A, B, AB\}$, the box can have the complex composition $c_{\text{box}} = \{A, B\}$ (corresponding to two monomers) or the composition $c_{\text{box}} = \{AB\}$ (corresponding to one dimer). Let $\Lambda$ denote the set of complex compositions that can form in the box subject to conservation of mass. Let $\phi_{\text{box}}$ denote all the sequence information in the box. The number of solvent molecules may be expressed

$$M_{\text{sol}} = [\text{H}_2\text{O}]\, V N_A, \tag{S76}$$

where $N_{\text{A}}$ is Avogadro's number, $[\text{H}_2\text{O}]$ is the molarity of water, and $V$ is the volume of the box.

The partition function of the box takes the form:[13]

$$Q(\phi_{\text{box}}) = Q_{\text{ref}} \sum_{c_{\text{box}} \in \Lambda} \Big[ M_{\text{sol}}^{|c_{\text{box}}|} \prod_{j \in c_{\text{box}}} Q(\phi_j) \Big], \tag{S77}$$

where $Q_{\text{ref}}$ may be chosen to set the reference state, $g_{\text{ref}} := -\log Q_{\text{ref}}$, of the free energy of the box:

$$\overline{g}(\phi_{\text{box}}) := -\log Q(\phi_{\text{box}}) = g_{\text{ref}} - \log \sum_{c_{\text{box}} \in \Lambda} \Big[ M_{\text{sol}}^{|c_{\text{box}}|} \prod_{j \in c_{\text{box}}} Q(\phi_j) \Big]. \tag{S78}$$

In (S77), each term in the sum represents the partition function for one composition $c_{\text{box}} \in \Lambda$. By definition, each complex partition function, $Q(\phi_j)$, is the sum of Boltzmann factors over alternative secondary structures $s_j$ for complex $j$, so substituting:

$$Q(\phi_{\text{box}}) = Q_{\text{ref}} \sum_{c_{\text{box}} \in \Lambda} \Big[ M_{\text{sol}}^{|c_{\text{box}}|} \prod_{j \in c_{\text{box}}} \sum_{s_j \in \overline{\Gamma}(\phi_j)} e^{-\overline{g}(\phi_j, s_j)} \Big]. \tag{S79}$$

For each composition $c_{\text{box}} \in \Lambda$, there are $|c_{\text{box}}|$ powers of $M_{\text{sol}}$, corresponding to an entropy of mixing term. Further, the product of complex partition functions causes the free energies in the exponentials of the Boltzmann factors for different complexes to add, leading to one partition function contribution for each secondary structure of the box, $s_{\text{box}}$, consistent with composition $c_{\text{box}}$. Let $s_j \in s_{\text{box}}$ denote the set of complex secondary structures constituent within $s_{\text{box}}$, one for each complex $j \in c_{\text{box}}$. Let $\overline{\Gamma}(\phi_{\text{box}})$ denote the set of all secondary structures of the box. By definition, the partition function of the box can be written as a Boltzmann-weighted sum over all the secondary structures that the box can adopt:

$$Q(\phi_{\text{box}}) = \sum_{s_{\text{box}} \in \overline{\Gamma}(\phi_{\text{box}})} e^{-\overline{g}(\phi_{\text{box}}, s_{\text{box}})}. \tag{S80}$$

Comparing (S80) to (S79), we then have for each $s_{\text{box}} \in \overline{\Gamma}(\phi_{\text{box}})$, the box secondary structure free energy:

$$\overline{g}(\phi_{\text{box}}, s_{\text{box}}) = g_{\text{ref}} - |c_{\text{box}}| \log M_{\text{sol}} + \sum_{s_j \in s_{\text{box}}} \overline{g}(\phi_j, s_j), \tag{S81}$$

The equilibrium probability of secondary structure $s_{\text{box}}$ is then

$$p(\phi_{\text{box}}, s_{\text{box}}) = e^{-\overline{g}(\phi_{\text{box}}, s_{\text{box}})} / Q(\phi_{\text{box}}). \tag{S82}$$

#### S8.2 Kullback-Leibler divergence

The Kullback-Leibler divergence

$$\mathcal{D}_{\mathrm{KL}}(U\,||\,V) \coloneqq \sum_{x\in\mathcal{X}} U(x)\log\frac{U(x)}{V(x)} \tag{S83}$$

$$= \mathbb{E}_{x\sim U}\left[\log\frac{U(x)}{V(x)}\right] = \mathbb{E}_{x\sim U}\left[\log U(x) - \log V(x)\right] \tag{S84}$$

measures the distance between the true probability distribution $U(x)$ and an approximating probability distribution $V(x)$.[8] $\mathcal{D}_{\mathrm{KL}}$ is non-negative and equals zero if and only if $U(x) = V(x)$.[8]

To assess the representational capacity of transfer matrix models (which employ a single functional form [trace of a matrix product] for all loop types and sizes) to reproduce NN reference free energy models (which employ different functional forms and lookup tables for different loop types and sizes), we define $\mathcal{D}_{\mathrm{KL}}$ in terms of secondary structure equilibrium probability distributions letting $U(x)$ correspond to the reference NN model and $V(x)$ to a transfer matrix model. As there are an infinite number of sequences and secondary structures over which these probability distributions could be defined, it is convenient to adopt form (S84), expressed as an expectation over: 1) box sequences, $\phi_{\mathrm{box}}$, sampled from a representative distribution $\Phi_{\mathrm{box}}$, and 2) secondary structures, $s_{\mathrm{box}}$, sampled from the Boltzmann distribution, $p^{\mathrm{lit}}(\phi_{\mathrm{box}})$, according to a literature NN reference model:

$$\begin{aligned}\mathcal{D}_{\mathrm{KL}}(\vec{g}_{37}) &\coloneqq \mathbb{E}_{\phi_{\mathrm{box}}\sim\Phi_{\mathrm{box}}}\left[\mathbb{E}_{s_{\mathrm{box}}\sim p^{\mathrm{lit}}(\phi_{\mathrm{box}})}\left[\log p^{\mathrm{lit}}(\phi_{\mathrm{box}}, s_{\mathrm{box}}) - \log p(\phi_{\mathrm{box}}, s_{\mathrm{box}}; \vec{g}_{37})\right]\right],\\ &= \mathbb{E}_{\phi_{\mathrm{box}}\sim\Phi_{\mathrm{box}}}\left[\overline{g}^{\mathrm{lit}}(\phi_{\mathrm{box}}) - \overline{g}(\phi_{\mathrm{box}};\vec{g}_{37}) + \mathbb{E}_{s_{\mathrm{box}}\sim p^{\mathrm{lit}}(\phi_{\mathrm{box}})}\left[\overline{g}(\phi_{\mathrm{box}}, s_{\mathrm{box}};\vec{g}_{37}) - \overline{g}^{\mathrm{lit}}(\phi_{\mathrm{box}}, s_{\mathrm{box}})\right]\right].\end{aligned} \tag{S85}$$

Here, $\vec{g}_{37}$ denotes the parameter set for the transfer matrix model (at reference temperature $T_{37} \coloneqq 310.15$ K), with values regressed via the minimization:

$$\min_{\vec{g}_{37}} \mathcal{D}_{\mathrm{KL}}(\vec{g}_{37}) = |\Psi_{\mathrm{boxes}}|^{-1} \sum_{\phi_{\mathrm{box}}\in\Psi_{\mathrm{boxes}}} \left\{\overline{g}^{\mathrm{lit}}_{\mathrm{box}}(\phi_{\mathrm{box}}) - \overline{g}(\phi_{\mathrm{box}};\vec{g}_{37}) + J^{-1}\sum_{s_{\mathrm{box}}\in\overline{\Gamma}_{\mathrm{sample}}(\phi_{\mathrm{box}};J)}\left[\overline{g}(\phi_{\mathrm{box}}, s_{\mathrm{box}};\vec{g}_{37}) - \overline{g}^{\mathrm{lit}}(\phi_{\mathrm{box}}, s_{\mathrm{box}})\right]\right\}, \tag{S86}$$

where $\Psi_{\mathrm{boxes}}$ denotes a set of $|\Psi_{\mathrm{boxes}}|$ box ensembles sampled from $\Phi_{\mathrm{box}}$ and $\overline{\Gamma}_{\mathrm{sample}}(\phi_{\mathrm{box}};J)$ represents a set of $J$ structures Boltzmann-sampled from box ensemble $\overline{\Gamma}(\phi_{\mathrm{box}})$ using the literature NN reference model. Calculation of $\mathcal{D}_{\mathrm{KL}}(\vec{g}_{37})$ for a given parameter set, $\vec{g}_{37}$, requires calculation of:

- Using the literature NN reference model and dynamic programs:[18]
  - The box free energy, $\overline{g}^{\mathrm{lit}}(\phi_{\mathrm{box}})$, for each box ensemble $\phi_{\mathrm{box}} \in \Psi_{\mathrm{boxes}}$.
  - The set of $J$ Boltzmann-sampled box structures, $\overline{\Gamma}_{\mathrm{sample}}(\phi_{\mathrm{box}};J)$, for each box ensemble $\phi_{\mathrm{box}} \in \Psi_{\mathrm{boxes}}$.
  - The box secondary structure free energy, $\overline{g}^{\mathrm{lit}}(\phi_{\mathrm{box}}, s_{\mathrm{box}})$, for each box secondary structure $s_{\mathrm{box}} \in \overline{\Gamma}_{\mathrm{sample}}(\phi_{\mathrm{box}};J)$ for each box ensemble $\phi_{\mathrm{box}} \in \Psi_{\mathrm{boxes}}$.
- Using the transfer matrix model and dynamic programs:
  - The box free energy, $\overline{g}(\phi_{\mathrm{box}};\vec{g}_{37})$, for each box ensemble $\phi_{\mathrm{box}} \in \Psi_{\mathrm{boxes}}$.
  - The box secondary structure free energy, $\overline{g}(\phi_{\mathrm{box}}, s_{\mathrm{box}};\vec{g}_{37})$, for each box secondary structure $s_{\mathrm{box}} \in \overline{\Gamma}_{\mathrm{sample}}(\phi_{\mathrm{box}};J)$ for each box ensemble $\phi_{\mathrm{box}} \in \Psi_{\mathrm{boxes}}$.

We favor using the box ensemble to evaluate the representational capacity of transfer matrix models for reference NN models because the box ensemble includes competition between alternative compositions, ensuring that $g^{\mathrm{assoc}}$ is properly exercised by the training and test data (for example, a box with alternative compositions $c_{\mathrm{box}} = \{A, B\}$ and $c_{\mathrm{box}} = \{AB\}$ such that strands $A$ and $B$ can form either monomer or dimer complexes in alternative compositions).

#### S8.3 Stochastic gradient descent method for K-L divergence optimization

We optimize the K-L divergence for a transfer matrix model with respect to a reference NN model using a stochastic gradient descent method (e.g., Reference 36). The gradient and Hessian of $\mathcal{D}_{\mathrm{KL}}$ with respect to the parameters, $\vec{g}_{37}$, have the forms:

$$
\begin{aligned}
v_i(\vec{g}_{37}) &\coloneqq \frac{\partial \mathcal{D}_{\mathrm{KL}}}{\partial \vec{g}_{37_i}} = -\mathbb{E}_{\phi_{\mathrm{box}}\sim\Phi_{\mathrm{box}}}\left[\frac{\partial \overline{g}(\phi_{\mathrm{box}};\vec{g}_{37})}{\partial \vec{g}_{37_i}} - \mathbb{E}_{s_{\mathrm{box}}\sim p^{\mathrm{lit}}(\phi_{\mathrm{box}})}\left[\frac{\partial g(\phi_{\mathrm{box}}, s_{\mathrm{box}};\vec{g}_{37})}{\partial \vec{g}_{37_i}}\right]\right] \\
H_{i,j}(\vec{g}_{37}) &\coloneqq \frac{\partial^2 \mathcal{D}_{\mathrm{KL}}}{\partial \vec{g}_{37_i}\,\partial \vec{g}_{37_j}} = \hat{H}_{i,j}(\vec{g}_{37}) \; - \; \mathbb{E}_{s_{\mathrm{box}}\sim p^{\mathrm{lit}}(\phi_{\mathrm{box}})}\left[\frac{\partial^2 \overline{g}(\phi_{\mathrm{box}};\vec{g}_{37})}{\partial \vec{g}_{37_i}\,\partial \vec{g}_{37_j}}\right] \; + \; \mathbb{E}_{s_{\mathrm{box}}\sim p^{\mathrm{lit}}(\phi_{\mathrm{box}})}\left[\frac{\partial^2 \overline{g}(\phi_{\mathrm{box}}, s_{\mathrm{box}};\vec{g}_{37})}{\partial \vec{g}_{37_i}\,\partial \vec{g}_{37_j}}\right] \\
\hat{H}_{i,j}(\vec{g}_{37}) &\coloneqq \mathbb{E}_{s_{\mathrm{box}}\sim p^{\mathrm{lit}}(\phi_{\mathrm{box}})}\left[\frac{\partial \overline{g}(\phi_{\mathrm{box}}, s_{\mathrm{box}};\vec{g}_{37})}{\partial \vec{g}_{37_i}}\frac{\partial \overline{g}(\phi_{\mathrm{box}}, s_{\mathrm{box}};\vec{g}_{37})}{\partial \vec{g}_{37_j}}\right] \\
&\quad - \mathbb{E}_{s_{\mathrm{box}}\sim p^{\mathrm{lit}}(\phi_{\mathrm{box}})}\left[\frac{\partial \overline{g}(\phi_{\mathrm{box}}, s_{\mathrm{box}};\vec{g}_{37})}{\partial \vec{g}_{37_i}}\right]\mathbb{E}_{s_{\mathrm{box}}\sim p^{\mathrm{lit}}(\phi_{\mathrm{box}})}\left[\frac{\partial \overline{g}(\phi_{\mathrm{box}}, s_{\mathrm{box}};\vec{g}_{37})}{\partial \vec{g}_{37_j}}\right]
\end{aligned}
\tag{S87}
$$

For a given box, these terms may be evaluated using dynamic programs to calculate the complex partition function (using the SumProduct evaluation algebra) and its derivatives (using the ∂SumProduct evaluation algebra). Analogous to the Gauss-Newton approximation for nonlinear least-squares,[47] our optimization algorithm uses $\hat{H}(\vec{g})$ as a positive semi-definite approximation to the full Hessian $H(\vec{g})$. This property follows from the fact that $\hat{H}_{i,j}(\vec{g})$ is the covariance of the vector-valued function $\nabla \overline{g}(\phi, s; \vec{g}_{37})$.

The K-L divergence, $\mathcal{D}_{\mathrm{KL}}$, is reduced using the stochastic gradient descent method of Algorithm S13 to optimize the parameter set, $\vec{g}_{37}$, for a given transfer matrix model. With this approach, at each optimization step, rather than calculate all components of the gradient, $v_i(\vec{g}_{37})\ \forall i$, and all components of the approximate Hessian, $\hat{H}_{i,j}(\vec{g}_{37})\ \forall i, j$, we calculate a single component of these quantities for stochastically sampled values of $i$ and $j$. $\vec{x}$ is then used to calculate a moving average of the gradient $\nabla$ (with decay scale $\beta_1$), $\hat{H}$ is used to calculate a moving average of the approximate Hessian $\nabla\nabla$ (with decay scale $\beta_2$ and a regularization parameter $\epsilon$ to ensure non-singularity). This approach is motivated by the ADAM optimizer,[36] which tracks moving averages of the gradient and squared gradient at each optimization step. Here, we incorporated a moving average of the approximate Hessian to enable a Newton step. While this approach necessitates a linear solve to invert the regularized approximate Hessian at each optimization step, we observed significantly faster convergence of the optimization process using this approach than using the standard ADAM methodology.

**Algorithm S13** Stochastic gradient descent with Hessian information

**function** HessianSGD($\mathcal{D}_{\mathrm{KL}}, \vec{g}, K, \alpha, \beta_1, \beta_2, \epsilon$)
  $\vec{x} \leftarrow 0$
  $\hat{H} \leftarrow 0$
  **for** $k = 0, 1, \ldots, K-1$ **do** ▷ *$\mathcal{L}^{(k)}$ stochastically sampled s.t. $\mathbb{E}_k[\mathcal{D}_{\mathrm{KL}}^{(k)}(\vec{g})] = \mathcal{D}_{\mathrm{KL}}(\vec{g})$*
    $\vec{x} \leftarrow \beta_1\vec{x} + (1-\beta_1)\nabla\mathcal{D}_{\mathrm{KL}}^{(k)}(\vec{g})$ ▷ *Moving average of gradient*
    $\hat{H} \leftarrow \beta_2\hat{H} + (1-\beta_2)\nabla\nabla\mathcal{D}_{\mathrm{KL}}^{(k)}(\vec{g})$ ▷ *Moving average of approximate Hessian*
    $\vec{g} \leftarrow \vec{g} - \alpha\left(\frac{1-\beta_1^k}{1-\beta_2^k}\hat{H} + \epsilon||\vec{x}||\mathbb{I}\right)^{-1}\vec{x}$ ▷ *Newton step using regularized approximate Hessian*
  **return** $\vec{g}$

Using NUPACK 4.1.0.1, the RNA reference model is `rna06` with the `stacking` ensemble and the DNA reference model is `dna04.3` with the `stacking` ensemble. Conditions were 37 °C with 1.0 M $Na^+$. The training set, $\Psi_{\mathrm{boxes}}^{\mathrm{train}}$, consisted of 180,000 random box ensembles. To generate each random box ensemble, $\phi_{\mathrm{box}} \in \Psi_{\mathrm{boxes}}^{\mathrm{train}}$, we use the following approach:

- generate a random number of strands to include the box ($L = 1 + x \,|\, x \sim G(0.5)$).
- generate a random number of nucleotides for each strand ($|\phi| = 1 + x \,|\, x \sim G(9)$).
- generate a random sequence $\phi$ for each base generated i.i.d. with equal probability.

where $G(x)$ is the geometric distribution with mean $x$. We define a complex of size $L$ with strand ordering $\pi$ and define the set of complexes in the box, $\Psi$, to contain all complexes that are subsets of $\pi$. For example, if $\pi = ABC$, then $\Psi = \{ABC, AB, BC, CA, A, B, C\}$. We set $M_{\mathrm{sol}} = 1$, corresponding to the high-concentration limit, to encourage reactions between these short random sequences.

To optimize parameters for a given transfer matrix model, all parameters in $\vec{g}_{37}$ were initialized to random independent Gaussians with mean 0 with standard deviation 0.01. 180,000 optimization steps were performed to reduce $\mathcal{D}_{\text{KL}}$ using stochastic gradient descent (Algorithm S13). Algorithm parameters were $\alpha = 10^{-3}$, $\beta_1 = 10^{-2}$, $\beta_2 = 10^{-3}$, and $\epsilon = 10^{-2}$. At each optimization step, $J = 100$ structures were Boltzmann sampled for each complex $\phi \in \mathbf{\Psi}_{\text{train}}$ to define $\overline{\Gamma}_{\text{sample}}(\phi; J)$ using the NN reference model.

To assess representational capacity and overfitting, a box test set $\mathbf{\Psi}^{\text{test}}_{\text{boxes}}$, of 100,000 random boxes was generated as above for $\mathbf{\Psi}^{\text{train}}_{\text{boxes}}$, and $\mathcal{D}_{\text{KL}}$, was calculated for $\mathbf{\Psi}^{\text{test}}_{\text{boxes}}$ without performing parameter optimization. Likewise, to compare different-material and historical NN models to NN reference models, $\mathcal{D}_{\text{KL}}$ was calculated for the box test set $\mathbf{\Psi}^{\text{test}}_{\text{boxes}}$ without performing parameter optimization.

#### S8.4 Divergence optimization results

Tables S5 (RNA) and S6 (DNA) and Figure S10 (both) display the K-L divergence with respect to NN reference models for the fixed-weight formulation, the floating-weight formulation, and the floating-block formulation for models sizes: $\nu \in \{1, 2, 3\}$ blocks and $\mu \in \{1, 2, 3, 4, 5\}$ block size with $\xi = 1$ (independent loop subensembles). As a basis for comparison, Table S7 displays the K-L divergence between historical and state-of-the-art NN models for a given material ($\mathcal{D}_{\text{KL}} \approx 0.5$) and between state-of-the-art NN models for different materials ($\mathcal{D}_{\text{KL}} > 1.5$).

Figure S10 demonstrates that RNA and DNA transfer matrix models achieve the representational capacity of existing NN models ($\mathcal{D}_{\text{KL}} < 0.5$) as the number of blocks, $\nu$, and block size, $\mu$, are increased. The fixed-weight formulation is somewhat constrained from approaching the existing NN RNA and DNA reference models given that it seeks to approximate logarithmic entropic costs for closed loops, a feature not present in the reference models for small hairpin or interior loops or for multiloops of any size. The floating-weight formulation has more flexibility to closely approximate the NN RNA and DNA reference models, and the floating-block formulation has the most flexibility to closely approximate the NN RNA and DNA reference models.

| | | Fixed Weights | | | | | | Floating Weights | | | | | | Floating Blocks | | | | | |
|---|---|---|---|---|---|---|---|---|---|---|---|---|---|---|---|---|---|---|---|
| $\nu$ | $\mu$ | $\lvert\vec{g}\rvert$ | train | test | ratio | STD | SEM | $\lvert\vec{g}\rvert$ | train | test | ratio | STD | SEM | $\lvert\vec{g}\rvert$ | train | test | ratio | STD | SEM |
| 1 | 1 | 17 | 3.77 | 3.75 | 0.99 | 5.37 | 0.017 | 34 | 1.67 | 1.67 | 1.00 | 2.88 | 0.009 | 24 | 1.67 | 1.67 | 1.00 | 2.88 | 0.009 |
| | 2 | 49 | 1.09 | 1.08 | 0.99 | 2.22 | 0.007 | 66 | 0.89 | 0.89 | 1.00 | 1.73 | 0.005 | 86 | 1.15 | 1.15 | 1.01 | 2.07 | 0.007 |
| | 3 | 101 | 0.63 | 0.62 | 0.99 | 1.39 | 0.004 | 118 | 0.55 | 0.55 | 1.00 | 1.13 | 0.004 | 188 | 0.40 | 0.40 | 1.00 | 1.00 | 0.003 |
| | 4 | 173 | 0.82 | 0.81 | 0.99 | 1.78 | 0.006 | 190 | 0.59 | 0.59 | 1.00 | 1.28 | 0.004 | 330 | 0.21 | 0.22 | 1.02 | 0.66 | 0.002 |
| | 5 | 265 | 0.36 | 0.36 | 1.00 | 0.85 | 0.003 | 282 | 0.51 | 0.51 | 0.99 | 1.29 | 0.004 | 512 | 0.09 | 0.10 | 1.01 | 0.22 | 0.001 |
| 2 | 1 | 19 | 2.72 | 2.70 | 0.99 | 3.89 | 0.012 | 45 | 0.46 | 0.46 | 1.00 | 0.92 | 0.003 | 35 | 0.46 | 0.46 | 1.00 | 0.92 | 0.003 |
| | 2 | 51 | 0.88 | 0.87 | 1.00 | 1.71 | 0.005 | 77 | 0.31 | 0.31 | 1.00 | 0.65 | 0.002 | 127 | 0.22 | 0.22 | 1.01 | 0.49 | 0.002 |
| | 3 | 103 | 0.52 | 0.53 | 1.00 | 1.10 | 0.003 | 129 | 0.24 | 0.24 | 1.00 | 0.55 | 0.002 | 279 | 0.15 | 0.15 | 1.00 | 0.40 | 0.001 |
| | 4 | 175 | 0.44 | 0.44 | 1.00 | 1.01 | 0.003 | 201 | 0.20 | 0.21 | 1.01 | 0.48 | 0.002 | 491 | 0.09 | 0.09 | 1.01 | 0.25 | 0.001 |
| | 5 | 267 | 0.39 | 0.39 | 1.01 | 0.91 | 0.003 | 293 | 0.14 | 0.14 | 1.00 | 0.32 | 0.001 | 763 | 0.06 | 0.06 | 1.02 | 0.15 | 0.000 |
| 3 | 1 | 21 | 2.38 | 2.37 | 0.99 | 3.37 | 0.011 | 56 | 0.44 | 0.44 | 1.00 | 0.85 | 0.003 | 46 | 0.44 | 0.44 | 1.00 | 0.85 | 0.003 |
| | 2 | 53 | 0.78 | 0.77 | 0.99 | 1.53 | 0.005 | 88 | 0.30 | 0.30 | 1.00 | 0.65 | 0.002 | 168 | 0.29 | 0.29 | 1.01 | 0.65 | 0.002 |
| | 3 | 105 | 0.47 | 0.47 | 1.00 | 1.00 | 0.003 | 140 | 0.22 | 0.22 | 1.00 | 0.49 | 0.002 | 370 | 0.13 | 0.13 | 1.01 | 0.33 | 0.001 |
| | 4 | 177 | 0.40 | 0.40 | 1.00 | 0.92 | 0.003 | 212 | 0.19 | 0.19 | 1.01 | 0.45 | 0.001 | 652 | 0.08 | 0.08 | 1.01 | 0.25 | 0.001 |
| | 5 | 269 | 0.34 | 0.35 | 1.03 | 0.80 | 0.003 | 304 | 0.13 | 0.13 | 1.01 | 0.31 | 0.001 | 1014 | 0.11 | 0.11 | 1.00 | 0.35 | 0.001 |

**Table S5:** Representational capacity of new RNA transfer matrix models relative to a literature reference model (cf. Figure 8): fixed-weight formulation (2.41), floating-weight formulation (2.42), floating-block formulation (S2.5). K-L divergence ($\mathcal{D}_{\mathrm{KL}}$): training set ($\Psi^{\mathrm{train}}_{\mathrm{boxes}}$) mean over 180,000 box ensembles, test set ($\Psi^{\mathrm{test}}_{\mathrm{boxes}}$) mean, STD, SEM over 100,000 box ensembles; ratio denotes the ratio of test to training means. Transfer matrix model parameters optimized for $\nu \in \{1, 2, 3\}$ blocks and $\mu \in \{1, 2, 3, 4, 5\}$ block size with $\xi = 1$ (independent loop subensembles) on training set. Reference model: `rna06` with `stacking` subensemble. DNA reference model `dna04.3` parameters with `stacking` subensemble. NUPACK 4.1.0.1. Conditions: 37 °C, 1.0 M $Na^{+}$.

| $\nu$ | $\mu$ | Fixed Weights | | | | | | Floating Weights | | | | | | Floating Blocks | | | | | |
|---|---|---|---|---|---|---|---|---|---|---|---|---|---|---|---|---|---|---|---|
| | | $\|\vec{g}\|$ | train | test | ratio | STD | SEM | $\|\vec{g}\|$ | train | test | ratio | STD | SEM | $\|\vec{g}\|$ | train | test | ratio | STD | SEM |
| 1 | 1 | 15 | 2.54 | 2.54 | 1.00 | 4.04 | 0.013 | 28 | 1.36 | 1.36 | 1.00 | 2.27 | 0.007 | 20 | 1.36 | 1.36 | 1.00 | 2.27 | 0.007 |
| | 2 | 41 | 0.79 | 0.79 | 1.00 | 1.52 | 0.005 | 54 | 0.59 | 0.59 | 1.01 | 1.10 | 0.003 | 70 | 0.55 | 0.55 | 1.01 | 1.07 | 0.003 |
| | 3 | 83 | 0.48 | 0.48 | 1.00 | 0.93 | 0.003 | 96 | 0.38 | 0.38 | 1.00 | 0.70 | 0.002 | 152 | 0.19 | 0.19 | 0.99 | 0.40 | 0.001 |
| | 4 | 141 | 0.59 | 0.59 | 1.00 | 1.19 | 0.004 | 154 | 0.36 | 0.36 | 1.00 | 0.70 | 0.002 | 266 | 0.14 | 0.14 | 1.01 | 0.28 | 0.001 |
| | 5 | 215 | 0.34 | 0.34 | 1.00 | 0.65 | 0.002 | 228 | 0.30 | 0.31 | 1.00 | 0.65 | 0.002 | 412 | 0.13 | 0.13 | 1.01 | 0.26 | 0.001 |
| 2 | 1 | 17 | 1.76 | 1.76 | 1.00 | 2.78 | 0.009 | 37 | 0.40 | 0.41 | 1.01 | 0.70 | 0.002 | 29 | 0.40 | 0.41 | 1.01 | 0.70 | 0.002 |
| | 2 | 43 | 1.09 | 1.09 | 1.00 | 1.81 | 0.006 | 63 | 0.33 | 0.34 | 1.01 | 0.62 | 0.002 | 103 | 0.18 | 0.18 | 1.00 | 0.29 | 0.001 |
| | 3 | 85 | 0.39 | 0.39 | 1.00 | 0.70 | 0.002 | 105 | 0.22 | 0.22 | 1.00 | 0.37 | 0.001 | 225 | 0.13 | 0.13 | 1.00 | 0.24 | 0.001 |
| | 4 | 143 | 0.35 | 0.35 | 1.00 | 0.62 | 0.002 | 163 | 0.17 | 0.17 | 1.01 | 0.29 | 0.001 | 395 | 0.11 | 0.11 | 1.01 | 0.19 | 0.001 |
| | 5 | 217 | 0.32 | 0.32 | 1.00 | 0.62 | 0.002 | 237 | 0.15 | 0.15 | 1.01 | 0.27 | 0.001 | 613 | 0.10 | 0.10 | 1.01 | 0.19 | 0.001 |
| 3 | 1 | 19 | 1.55 | 1.55 | 1.00 | 2.43 | 0.008 | 46 | 0.38 | 0.38 | 1.01 | 0.64 | 0.002 | 38 | 0.38 | 0.38 | 1.01 | 0.64 | 0.002 |
| | 2 | 45 | 0.56 | 0.57 | 1.01 | 1.03 | 0.003 | 72 | 0.32 | 0.32 | 1.01 | 0.57 | 0.002 | 136 | 0.17 | 0.17 | 1.00 | 0.28 | 0.001 |
| | 3 | 87 | 0.36 | 0.36 | 1.00 | 0.65 | 0.002 | 114 | 0.21 | 0.21 | 1.01 | 0.34 | 0.001 | 298 | 0.13 | 0.13 | 1.00 | 0.21 | 0.001 |
| | 4 | 145 | 0.41 | 0.42 | 1.01 | 0.79 | 0.003 | 172 | 0.17 | 0.17 | 1.01 | 0.29 | 0.001 | 524 | 0.10 | 0.10 | 1.00 | 0.18 | 0.001 |
| | 5 | 219 | 0.29 | 0.29 | 1.00 | 0.56 | 0.002 | 246 | 0.14 | 0.14 | 1.00 | 0.25 | 0.001 | 814 | 0.08 | 0.08 | 0.99 | 0.15 | 0.000 |

**Table S6:** Representational capacity of new DNA transfer matrix models relative to a literature reference model (cf. Figure 8): fixed-weight formulation (2.41), floating-weight formulation (2.42), floating-block formulation (S2.5). K-L divergence ($\mathcal{D}_{\mathrm{KL}}$): training set ($\Psi^{\mathrm{train}}_{\mathrm{boxes}}$) mean over 180,000 box ensembles, test set ($\Psi^{\mathrm{test}}_{\mathrm{boxes}}$) mean, STD, SEM over 100,000 box ensembles; ratio denotes the ratio of test to training means. Transfer matrix model parameters optimized for $\nu \in \{1, 2, 3\}$ blocks and $\mu \in \{1, 2, 3, 4, 5\}$ block size with $\xi = 1$ (independent loop subensembles) on training set. Reference model `dna04.3` parameters with `stacking` subensemble. NUPACK 4.1.0.1. Conditions: 37 °C, 1.0 M $Na^+$.

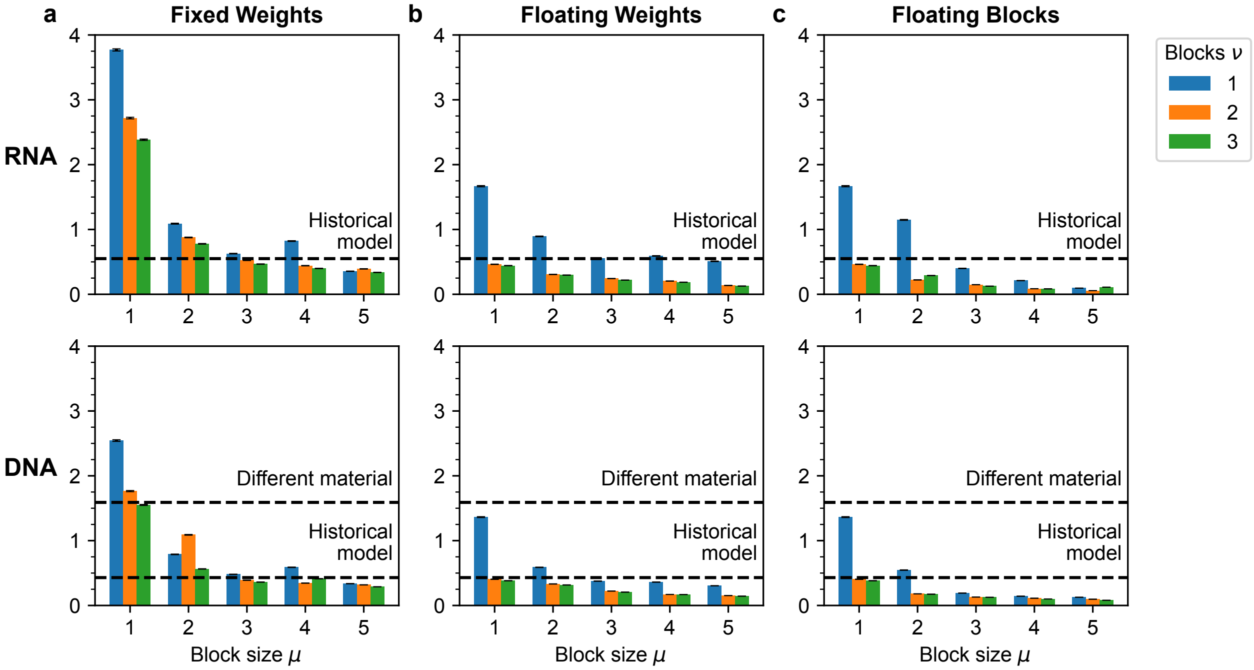

**Figure S10:** Representation capacity of new transfer matrix models relative to literature reference models (cf. Figure 8): a) Fixed-weight formulation (2.41), b) Floating-weight formulation (2.42), c) Floating-block formulation (S2.5). K-L divergence ($\mathcal{D}_{\mathrm{KL}}$): mean ± SEM over test set ($\Psi^{\mathrm{test}}_{\mathrm{boxes}}$) of 100,000 box ensembles. Transfer matrix model parameters optimized for $\nu \in \{1, 2, 3\}$ blocks and $\mu \in \{1, 2, 3, 4, 5\}$ block size with $\xi = 1$ (independent loop subensembles). Dashed lines denote mean values from Table S7. RNA studies (top): reference model: `rna06` (RNA) with `stacking` subensemble; historical model: `rna95` (RNA) with `no stacking` subensemble; different material model: `dna04.3` (DNA) with `stacking` subensemble. DNA studies (bottom): reference model `dna04.3` (DNA) parameters with `stacking` subensemble; historical model: `dna04.1` (DNA) parameters with `no stacking` subensemble; different material model: `rna06` (RNA) with `stacking` subensemble. NUPACK 4.1.0.1. Conditions: 37 °C, 1.0 M $Na^{+}$.

| Reference model | | | Test model | | | | $\mathcal{D}_{\mathrm{KL}}$ | | |
|---|---|---|---|---|---|---|---|---|---|
| Material | Parameters | Ensemble | Material | Parameters | Ensemble | Notes | Mean | STD | SEM |
| RNA | `rna06` | `stacking` | RNA | `rna95` | `nostacking` | Historical | 0.55 | 0.92 | 0.0029 |
| RNA | `rna06` | `stacking` | DNA | `dna04.1` | `nostacking` | Different material | — | — | — |
| DNA | `dna04.3` | `stacking` | DNA | `dna04.1` | `nostacking` | Historical | 0.43 | 0.63 | 0.0020 |
| DNA | `dna04.3` | `stacking` | RNA | `rna95` | `nostacking` | Different material | 1.59 | 2.66 | 0.0084 |

**Table S7:** Kullback-Leibler divergence ($\mathcal{D}_{\mathrm{KL}}$) between historical and reference NN models or between different-material and reference NN models. K-L divergence ($\mathcal{D}_{\mathrm{KL}}$): mean, STD, and SEM over test set ($\Psi^{\mathrm{test}}_{\mathrm{boxes}}$) of 100,000 box ensembles. Mean values are displayed as dashed lines in Figure S10. NUPACK 4.1.0.1. Conditions: 37 °C, 1.0 M $Na^{+}$. Note that for the RNA reference model, the different material (DNA) test model is not evaluated because the RNA ensemble contains wobble pairs that are not in the DNA ensemble, leading $\mathcal{D}_{\mathrm{KL}}$ to be formally infinite.

#### S9 Regression of transfer matrix and tensor model parameters based on experimental melt data

##### S9.1 Database of reaction free energies and enthalpies for literature melt experiments

To enable regression of parameters for the new class of tensor models, we assembled databases of reaction free energies and enthalpies for literature melt experiments for both RNA[2,3,7,9,10,17,20–31,33–35,37,38,42,44–46,50,51,53,56,57,60–66,68–70,74–76,78,80], and DNA.[1–3,5,6,15,16,29,32,48,49,54,55,59,71,72,79] In these literature studies, UV absorbance data were fit with a two-state model as described previously[51] and briefly summarized here.

**1-stranded experiments.** For 1-stranded melt experiments, the strand was assumed to be either in an unpaired state (u) or in a paired state (p):

$$\underbrace{\mathrm{A_u}}_{\text{unpaired state}} \quad \leftrightarrow \quad \underbrace{\mathrm{A_p}}_{\text{paired state}} \quad . \tag{S88}$$

The reaction enthalpy and entropy (both assumed to be temperature-independent),

$$\Delta H^{\text{lit}}_{\text{rxn}} = \Delta H_{\text{p}} - \Delta H_{\text{u}}, \tag{S89}$$

$$\Delta S^{\text{lit}}_{\text{rxn}} = \Delta S_{\text{p}} - \Delta S_{\text{u}}, \tag{S90}$$

were regressed in the literature from the melt data.

**2-stranded experiments.** For 2-stranded melt experiments with non-self-complementary sequences ($\mathrm{A + B \leftrightarrow AB}$), each strand was assumed to be either in a unpaired state (A and B) or in a paired state (AB):

$$\underbrace{\mathrm{A+B}}_{\text{unpaired state}} \quad \leftrightarrow \quad \underbrace{\mathrm{AB}}_{\text{paired state}} \quad . \tag{S91}$$

The reaction enthalpy and entropy (both assumed to be temperature-independent),

$$\Delta H^{\text{lit}}_{\text{rxn}} = \Delta H_{\text{AB}} - \Delta H_{\text{A}} - \Delta H_{\text{B}}, \tag{S92}$$

$$\Delta S^{\text{lit}}_{\text{rxn}} = \Delta S_{\text{AB}} - \Delta S_{\text{A}} - \Delta S_{\text{B}}, \tag{S93}$$

were regressed in the literature from the melt data.

For 2-stranded melt experiments with self-complementary sequences ($\mathrm{2A \leftrightarrow AA}$), each strand was assumed to be either in a unpaired state (A) or in a paired state (AA):

$$\underbrace{\mathrm{2A}}_{\text{unpaired state}} \quad \leftrightarrow \quad \underbrace{\mathrm{AA}}_{\text{paired state}} \quad . \tag{S94}$$

The reaction enthalpy and entropy (both assumed to be temperature-independent),

$$\Delta H^{\text{lit}}_{\text{rxn}} = \Delta H_{\text{AA}} - 2\Delta H_{\text{A}}, \tag{S95}$$

$$\Delta S^{\text{lit}}_{\text{rxn}} = \Delta S_{\text{AA}} - 2\Delta S_{\text{A}}, \tag{S96}$$

were regressed in the literature from the melt data.

**Reaction free energies.** For either 1-stranded or 2-stranded reactions, the reaction enthalpy and entropy were used to calculate the reaction free energy:

$$\Delta G^{\text{lit}}_{\text{rxn},37} = \Delta H^{\text{lit}}_{\text{rxn}} - T_{37}\Delta S^{\text{lit}}_{\text{rxn}}, \tag{S97}$$

at $T_{37} \coloneqq 310.15$ K. Literature studies typically report $\Delta G^{\text{lit}}_{\text{rxn},37}$ and $\Delta H^{\text{lit}}_{\text{rxn}}$ for each melt experiment. The reaction free energies and enthalpies for multiple melt experiments were then used to regress $\Delta G_{37}(\text{loop})$ and $\Delta H(\text{loop})$ values for existing loop-based free energy models based on the 2-state definitions above where each state was defined to be a single secondary structure (no base pairs formed for the unpaired state, all base pairs formed for the paired state), yielding a temperature dependent free energy model (see Section S2.3).

**Experimental data sets.** Here, to provide a basis for regressing parameters for new transfer matrix and tensor models, we compiled $\Delta G^{\mathrm{lit}}_{\mathrm{rxn},37}$ and $\Delta H^{\mathrm{lit}}_{\mathrm{rxn}}$ values for large sets of literature experiments for RNA (895 melts) and DNA (464 melts) in standard salt conditions (1 M $Na^{+}$). For each material, we assigned ≈80% of experiments to a training set to use for parameter regression, and the other ≈20% of experiments to a test set to assess model accuracy as well as to monitor overfitting.

#### S9.2 Calculation of reaction free energies and enthalpies using transfer matrix and tensor models

UV melt experiments measure temperature-dependent absorbance, which may be expressed as a sum over all secondary structures $s \in \Gamma(\phi)$:

$$A(\phi,T) = \sum_{s\in\Gamma(\phi)} A(\phi,s,T)p(\phi,s,T), \tag{S98}$$

where $A(\phi,s,T)$ is the absorbance of sequence $\phi$ in secondary structure $s$ at temperature $T$ and $p(\phi,s,T)$ is the equilibrium probability of secondary structure $s$ for sequence $\phi$ at temperature $T$. A detailed sequence- and structure-dependent model for absorbance is unknown, but it is well-established that the absorbance of unpaired bases is higher than for base pairs. Literature models are regressed using a 2-state approximation using a single secondary structure to represent the unpaired state and a single secondary structure to represent the paired state. In reality, we would expect the absorbance of the unpaired and paired states to depend on an equilibrium distribution of secondary structures. For this reason, we regress parameters for transfer matrix and tensor models using a 2-ensemble approximation where the unpaired and paired states are each modeled by an ensemble of structures.

**1-stranded experiments.** For 1-stranded experiments, the strand was assumed to be either in an unpaired state with no base pairs formed (u; a single structure) or in a paired state with one or more base pairs formed (p; all other structures in the complex ensemble of the single strand):

$$\underbrace{\mathrm{A_u}}_{\text{unpaired state}} \quad \leftrightarrow \quad \underbrace{\mathrm{A_p}}_{\text{paired state}} \quad . \tag{S99}$$

The reaction free energy is calculated as follows:

$$\begin{aligned}\Delta G_{\mathrm{rxn}}(\phi;\vec{g}) &\coloneqq \Delta G_{\mathrm{p}}(\phi;\vec{g}) - \Delta G_{\mathrm{u}}(\phi;\vec{g}),\\ &= -k_BT\log[Q(\phi;\vec{g}) - Q_u(\phi;\vec{g})] + k_BT\log Q_u(\phi;\vec{g}),\\ &= -k_BT\log[Q(\phi;\vec{g}) - e^{-\Delta G(\phi,s_u;\vec{g})/k_BT}] - \Delta G(\phi,s_u;\vec{g}).\end{aligned} \tag{S100}$$

Let $\vec{g}_{37}$ denote $\vec{g}$ at reference temperature $T_{37} \coloneqq 310.15$ K so that:

$$\Delta G_{\mathrm{rxn},37}(\phi;\vec{g}_{37}) \coloneqq \Delta G_{\mathrm{rxn}}(\phi;\vec{g})\big|_{T_{37}}. \tag{S101}$$

The complex partition function $\overline{Q}(\phi;\vec{g})$ is calculated using the SumProduct evaluation algebra. Note that no distinguishability correction[13] is required for a 1-stranded complex, so $Q(\phi) = \overline{Q}(\phi)$. The reaction enthalpy (assumed to be temperature-independent) is calculated as follows:

$$\begin{aligned}\Delta H_{\mathrm{rxn}} &= \frac{\partial(\beta\Delta G_{\mathrm{rxn}}(\phi;\vec{g}))}{\partial\beta}\bigg|_{T_{37}} = \frac{\partial g_{\mathrm{rxn}}(\phi;\vec{g})}{\partial\beta}\bigg|_{T_{37}}\\ &= \beta^{-1}\sum_i \frac{\partial g_{\mathrm{rxn}}(\phi_r;\vec{g})}{\partial\vec{g}_i}\vec{h}_i\bigg|_{T_{37}}\end{aligned} \tag{S102}$$

where

$$\vec{h}_i = \beta\frac{\partial\vec{g}_i}{\partial\beta}\bigg|_{T_{37}}. \tag{S103}$$

and $\vec{g}_i$ is the $i$th entry of $\vec{g}$ and $\vec{h}_i$ is the $i$th entry of $\vec{h}$. The $\partial$SumProduct evaluation algebra is used to evaluate $\partial\overline{Q}(\phi;\vec{g})/\partial\vec{g}_i$ as an intermediate step to evaluating $\partial g_{\mathrm{rxn}}(\phi_r;\vec{g})/\partial\vec{g}_i$ for each parameter $i$.

**2-stranded experiments.** For 2-stranded experiments with non-self-complementary sequences (A + B ↔ AB), each strand was assumed to be either in a monomer state (A or B) or dimer state (AB):

$$\underbrace{\mathrm{A+B}}_{\text{monomer state}} \quad \leftrightarrow \quad \underbrace{\mathrm{AB}}_{\text{dimer state}} \quad . \tag{S104}$$

The reaction free energy is calculated as:

$$\Delta G_{\mathrm{rxn}}(\phi;\vec{g}) \coloneqq \Delta G(\phi_{\mathrm{AB}};\vec{g}) - \Delta G(\phi_{\mathrm{A}};\vec{g}) - \Delta G(\phi_{\mathrm{B}};\vec{g}). \tag{S105}$$

At reference temperature $T_{37} \coloneqq 310.15$ K, we have:

$$\Delta G_{\mathrm{rxn},37}(\phi;\vec{g}_{37}) \coloneqq \Delta G_{\mathrm{rxn}}(\phi;\vec{g})\big|_{T_{37}}. \tag{S106}$$

The complex partition functions $\overline{Q}(\phi_{\mathrm{A}};\vec{g})$, $\overline{Q}(\phi_{\mathrm{B}};\vec{g})$, and $\overline{Q}(\phi_{\mathrm{AB}};\vec{g})$ are calculated using the SumProduct evaluation algebra. Note that no distinguishability correction[13] is required for a 1-stranded complex or for a 2-stranded complex with non-self-complementary strands, so $Q(\phi_{\mathrm{A}}) = \overline{Q}(\phi_{\mathrm{A}})$, $Q(\phi_{\mathrm{B}}) = \overline{Q}(\phi_{\mathrm{B}})$, and $Q(\phi_{\mathrm{AB}}) = \overline{Q}(\phi_{\mathrm{AB}})$. In each case, the complex free energy is defined by $\Delta G(\phi;\vec{g}) \coloneqq -k_B T \log Q(\phi)$.

For 2-stranded experiments with self-complementary sequences (2A ↔ AA), each strand was assumed to be either in a monomer state (A) or dimer state (AA):

$$\underbrace{\mathrm{2A}}_{\text{monomer state}} \quad \leftrightarrow \quad \underbrace{\mathrm{AA}}_{\text{dimer state}} \quad . \tag{S107}$$

The reaction free energy is calculated as:

$$\Delta G_{\mathrm{rxn}}(\phi;\vec{g}) \coloneqq \Delta G(\phi_{\mathrm{AA}};\vec{g}) - 2\Delta G(\phi_{\mathrm{A}};\vec{g}). \tag{S108}$$

At reference temperature $T_{37} \coloneqq 310.15$ K, we have:

$$\Delta G_{\mathrm{rxn},37}(\phi;\vec{g}_{37}) \coloneqq \Delta G_{\mathrm{rxn}}(\phi;\vec{g})\big|_{T_{37}}. \tag{S109}$$

The complex partition functions $\overline{Q}(\phi_{\mathrm{A}};\vec{g})$ and $\overline{Q}(\phi_{\mathrm{AA}};\vec{g})$ are calculated using the SumProduct evaluation algebra. Note that no distinguishability correction[13] is required for a 1-stranded complex, so $Q(\phi_{\mathrm{A}}) = \overline{Q}(\phi_{\mathrm{A}})$. On the other hand, there is a distinguishability correction[13] for a 2-stranded complex with self-complementary strands, so $Q(\phi_{\mathrm{AA}}) = \overline{Q}(\phi_{\mathrm{AA}})/2$. In each case, the complex free energy is defined by $\Delta G(\phi;\vec{g}) \coloneqq -k_B T \log Q(\phi)$.

For 2-stranded experiments with either non-self-complementary or self-complementary strands, the reaction enthalpy (assumed to be temperature-independent) is calculated as in (S102) and the ∂SumProduct evaluation algebra is used to evaluate $\partial\overline{Q}(\phi_{\mathrm{A}};\vec{g})/\partial\vec{g}_i$, $\partial\overline{Q}(\phi_{\mathrm{B}};\vec{g})/\partial\vec{g}_i$, and $\partial\overline{Q}(\phi_{\mathrm{AB}};\vec{g})/\partial\vec{g}_i$ for each parameter $i$ at $T_{37}$. For a 2-stranded complex with self-complementary strands, a distinguishability correction[13] is applied as above (i.e., $\partial Q(\phi_{\mathrm{AA}};\vec{g})/\partial\vec{g}_i = \frac{1}{2}\partial\overline{Q}(\phi_{\mathrm{AA}};\vec{g})/\partial\vec{g}_i$).

#### S9.3 Calculation of reaction free energies and enthalpies for nearest-neighbor reference models

For NN reference models, the reaction free energy, $\Delta G_{\mathrm{rxn}}$, was calculated as in Section S9.2 using an NN reference model and algorithm[19] and the reaction enthalpies were estimated using a finite difference

$$\Delta H_{\mathrm{rxn}} \approx \frac{T_{38}\Delta G_{\mathrm{rxn}}(\phi;T_{38}) - T_{36}\Delta G_{\mathrm{rxn}}(\phi;T_{36})}{2\,\mathrm{K}} \tag{S110}$$

with $T_{38} \coloneqq T_{37} + 1\,\mathrm{K}$ and $T_{36} \coloneqq T_{37} - 1\,\mathrm{K}$.

#### S9.4 Objective functions for regressing transfer matrix and tensor model free energy and enthalpy parameters

Parameter regression is performed in a two-step process. First, we regress the free energy parameters $\vec{g}_{37}$ corresponding to reference temperature $T_{37}$. Second, we regress the enthalpy parameters $\vec{h}$, which are assumed to be temperature-independent. Temperature-dependent free energy parameters $\vec{g}$ are then defined by (S3).

The objective function for regressing free energy parameter set, $\vec{g}_{37}$, via nonlinear least squares optimization takes the form

$$\min_{\vec{g}_{37}} \mathcal{L}(\vec{g}_{37}, \mathcal{R}) \quad \text{where} \quad \mathcal{L}(\vec{g}_{37}) \coloneqq |\mathcal{R}|^{-1} \sum_{r\in\mathcal{R}} \left[\Delta G_{\text{rxn},37}(\phi_r; \vec{g}_{37}) - \Delta G^{\text{lit}}_{\text{rxn},37,r}\right]^2, \tag{S111}$$

where for reaction $r$ in reaction set $\mathcal{R}$, $\Delta G^{\text{lit}}_{\text{rxn},37,r}$ is the literature reaction free energy regressed from an experimental melt curve (see Section S9.1) and $\Delta G_{\text{rxn},37}(\phi_r; \vec{g}_{37})$ is the reaction free energy calculated using a transfer matrix or tensor model (see Section S9.2) using $\vec{g}_{37}$. Optimization is performed using a Gauss-Newton trust-region method (see Section S9.5).

The objective function for regressing enthalpy parameter set, $\vec{h}$, via linear least squares takes the form

$$\min_{\vec{h}} \mathcal{L}(\vec{h}, \vec{g}_{37}, \mathcal{R}) \quad \text{where} \quad \mathcal{L}(\vec{h}, \vec{g}_{37}, \mathcal{R}) \coloneqq \alpha\|\vec{h}\|_2^2 + |\mathcal{R}|^{-1} \sum_{r\in\mathcal{R}} (\Delta H_{\text{rxn}}(\phi_r; \vec{h}, \vec{g}_{37}) - \Delta H^{\text{lit}}_{\text{rxn},r})^2, \tag{S112}$$

where $\Delta H^{\text{lit}}_{\text{rxn},r}$ is the literature reaction enthalpy regressed from an experimental melt curve (see Section S9.1) and $\Delta H_{\text{rxn}}(\phi_r; \vec{h}, \vec{g}_{37})$ is the reaction enthalpy calculated using a transfer matrix or tensor model (see Section S9.2) using $\vec{g}_{37}$ and $\vec{h}$, and we have introduced a small regularization term, $\alpha\|\vec{h}\|_2^2$, to avoid ill-conditioning (using $\alpha \coloneqq 10^{-3}$). Note that in calculating the reaction enthalpy,

$$\Delta H_{\text{rxn}}(\phi_r; \vec{h}, \vec{g}_{37}) = \frac{\partial g_{\text{rxn}}(\phi_r; \vec{g})}{\partial \beta}\bigg|_{T_{37}} = \sum_i \left(\beta^{-1} \frac{\partial g_{\text{rxn}}(\phi_r; \vec{g})}{\partial \vec{g}_i}\bigg|_{T_{37}}\right) \vec{h}_i, \tag{S113}$$

with

$$\vec{h}_i = \beta \frac{\partial \vec{g}_i}{\partial \beta}\bigg|_{T_{37}}, \tag{S114}$$

the $\partial g_{\text{rxn}}/\partial \vec{g}_i$ terms are independent of $\vec{h}$, so regression of the enthalpy parameter set, $\vec{h}$, takes the form of standard linear least squares. Collecting a row of $\partial g_{\text{rxn}}/\partial \vec{g}_i$ for each experiment into a matrix $M$ and the experimental reaction enthalpies into a column vector $b$, our regressed parameter set is given by

$$\vec{h} = (M^\top M + \alpha|\mathcal{R}|\mathbb{I})^{-1} M^\top b. \tag{S115}$$

#### S9.5 Gauss-Newton trust region method for free energy parameter regression

We implemented fits to experimental melt data free energies using a Gauss-Newton trust-region method.[47] In this approach, we consider our objective function (S111) and its Hessian:

$$\begin{aligned} H_{i,j}(\vec{g}) &\coloneqq \frac{\partial^2}{\partial \vec{g}_{37_i} \partial \vec{g}_{37_j}} |\mathcal{R}|^{-1} \sum_{r\in\mathcal{R}} \left[\Delta G_{\text{rxn},37}(\phi_r; \vec{g}_{37}) - \Delta G^{\text{lit}}_{\text{rxn},37,r}\right]^2 \\ &= 2|\mathcal{R}|^{-1} \sum_{r\in\mathcal{R}} \left( \frac{\partial \Delta G_{\text{rxn},37}(\phi_r; \vec{g}_{37})}{\partial \vec{g}_{37_i}} \frac{\partial \Delta G_{\text{rxn},37}(\phi_r; \vec{g}_{37})}{\partial \vec{g}_{37_j}} + \left[\Delta G_{\text{rxn},37}(\phi_r; \vec{g}_{37}) - \Delta G^{\text{lit}}_{\text{rxn},37,r}\right] \frac{\partial^2 \Delta G_{\text{rxn},37}(\phi_r; \vec{g}_{37})}{\partial \vec{g}_{37_i} \partial \vec{g}_{37_j}} \right) \end{aligned} \tag{S116}$$

from which the Gauss-Newton approximation yields:

$$\hat{H}_{i,j}(\vec{g}) \coloneqq 2|\mathcal{R}|^{-1} \sum_{r\in\mathcal{R}} \frac{\partial \Delta G_{\text{rxn},37}(\phi_r; \vec{g}_{37})}{\partial \vec{g}_{37_i}} \frac{\partial \Delta G_{\text{rxn},37}(\phi_r; \vec{g}_{37})}{\partial \vec{g}_{37_j}}, \tag{S117}$$

such that $\hat{H}_{i,j}(\vec{g})$ is a symmetric positive-semidefinite approximation to $H_{i,j}(\vec{g})$.

For regression to experimental data, each parameter set $\vec{g}_{37}$ was initialized to the values obtained from divergence-based regression to the existing NN reference model in Section S8. In each iteration, we then compute the objective gradient

$$v_i(\vec{g}) \coloneqq \frac{\partial \Delta G_{\text{rxn},37}(\phi_r; \vec{g}_{37})}{\partial \vec{g}_{37_i}} \tag{S118}$$

and the approximate Hessian $\hat{H}$. Given a trust-region radius $\delta$, we then solve the quadratic problem

$$\min_{\substack{x \\ \|x\| \le \delta}} \quad v^\top x + \tfrac{1}{2} x^\top \hat{H} x, \tag{S119}$$

to yield the parameter update:

$$x = \begin{cases} -\hat{H}^{-1}v & \|\hat{H}^{-1}v\| \leq \delta \\ -(\hat{H} + \lambda\mathbb{I})^{-1}v & \text{otherwise} \end{cases}, \tag{S120}$$

with $\vec{g} \leftarrow \vec{g} + x$. In the second case, scalar $\lambda$ is the unique solution of $\|(\hat{H} + \lambda\mathbb{I})^{-1}v\| = \delta$; $\lambda$ is calculated by eigendecomposition of $\hat{H}$ and subsequent bisection in $\lambda$ to numerical precision (starting from values $\lambda_{\text{low}} = 0$ and $\lambda_{\text{high}} = 1$, doubling $\lambda_{\text{high}}$ until $\|(\hat{H} + \lambda_{\text{high}}\mathbb{I})^{-1}v\| < \delta$). The trust-region radius $\delta$ is then updated using Algorithm S14. Within each iteration, this parameter and trust region update procedure is repeated until a decrease in the objective is achieved.

**Algorithm S14** Trust region radius update

```
function NextTrustRegionRadius(δ, ρ)
    if ρ < 1/4 then
        return δ/4
    else if ρ > 3/4 then
        return 2δ
    else
        return δ
```

#### S9.6 Free energy and enthalpy parameter regression results

For the fixed-weight formulation, floating-weight formulation, and floating-block formulation, root mean square errors (RMSE) for reaction free energy, $\Delta G_{\text{rxn},37}$, and reaction enthalpy, $\Delta H_{\text{rxn}}$ are reported for training sets and test sets for a variety of model sizes $(\nu, \mu, \xi)$ in Table S8 for RNA and Table S10 for DNA. The corresponding RMSEs for NN reference models are shown in Table S9 for RNA and Table S11 for DNA. Scatter plots of predicted vs experimental values of $\Delta G_{\text{rxn},37}$ and $\Delta H_{\text{rxn}}$ are shown for NN reference models and representative transfer matrix models with $(\nu, \mu, \xi) = 2, 2, 1$ and either fixed weights, floating weights, or floating blocks in Figure S11 for RNA and Figure S12 for DNA.

| | | | Fixed Weights | | | | | | | Floating Weights | | | | | | | Floating Blocks | | | | | | |
|---|---|---|---|---|---|---|---|---|---|---|---|---|---|---|---|---|---|---|---|---|---|
| | | | | $\Delta G_{\mathrm{rxn},37}$ (kcal/mol) | | | $\Delta H_{\mathrm{rxn}}$ (kcal/mol) | | | | $\Delta G_{\mathrm{rxn},37}$ (kcal/mol) | | | $\Delta H_{\mathrm{rxn}}$ (kcal/mol) | | | | $\Delta G_{\mathrm{rxn},37}$ (kcal/mol) | | | $\Delta H_{\mathrm{rxn}}$ (kcal/mol) | | |
| $\nu$ | $\mu$ | $\xi$ | $\lvert\vec{g}\rvert$ | train | test | ratio | train | test | ratio | $\lvert\vec{g}\rvert$ | train | test | ratio | train | test | ratio | $\lvert\vec{g}\rvert$ | train | test | ratio | train | test | ratio |
| 1 | 1 | 1 | 17 | 2.36 | 2.08 | 0.88 | 17.44 | 17.51 | 1.00 | 34 | 1.65 | 1.51 | 0.92 | 12.63 | 12.16 | 0.96 | 24 | 1.65 | 1.51 | 0.92 | 12.63 | 12.16 | 0.96 |
| | 2 | 1 | 49 | 1.36 | 1.05 | 0.77 | 12.16 | 11.48 | 0.94 | 66 | 1.29 | 1.11 | 0.86 | 11.10 | 11.67 | 1.05 | 86 | 1.32 | 1.23 | 0.93 | 10.69 | 10.89 | 1.02 |
| | | 2 | 73 | 1.35 | 1.06 | 0.79 | 12.16 | 11.53 | 0.95 | 90 | 1.27 | 1.09 | 0.86 | 10.97 | 11.34 | 1.03 | 134 | 1.25 | 1.20 | 0.96 | 10.81 | 11.50 | 1.06 |
| | 3 | 1 | 101 | 1.12 | 1.08 | 0.96 | 10.78 | 11.33 | 1.05 | 118 | 1.12 | 1.04 | 0.93 | 10.41 | 10.68 | 1.03 | 188 | 0.98 | 1.06 | 1.08 | 9.78 | 10.21 | 1.04 |
| | | 2 | 155 | 1.05 | 1.08 | 1.03 | 9.26 | 10.75 | 1.16 | 172 | 1.11 | 1.00 | 0.90 | 9.98 | 10.68 | 1.07 | 296 | 0.94 | 0.98 | 1.04 | 9.03 | 9.74 | 1.08 |
| | | 3 | 209 | 1.09 | 1.01 | 0.93 | 10.23 | 10.43 | 1.02 | 226 | 1.05 | 1.05 | 1.00 | 10.12 | 10.96 | 1.08 | 404 | 0.93 | 1.11 | 1.19 | 9.29 | 11.40 | 1.23 |
| | 4 | 1 | 173 | 0.94 | 1.00 | 1.06 | 9.77 | 11.41 | 1.17 | 190 | 0.99 | 1.16 | 1.17 | 9.81 | 12.43 | 1.27 | 330 | 0.83 | 1.16 | 1.40 | 10.07 | 12.87 | 1.28 |
| | | 2 | 269 | 0.92 | 0.97 | 1.05 | 9.61 | 10.52 | 1.09 | 286 | 0.96 | 1.15 | 1.20 | 9.77 | 11.04 | 1.13 | 522 | 0.90 | 0.88 | 0.98 | 9.39 | 9.74 | 1.04 |
| | | 3 | 365 | 0.85 | 0.91 | 1.07 | 8.91 | 10.66 | 1.20 | 382 | 0.91 | 0.93 | 1.02 | 9.47 | 9.81 | 1.04 | 714 | 0.88 | 1.01 | 1.15 | 8.93 | 11.02 | 1.23 |
| 2 | 1 | 1 | 19 | 2.24 | 1.97 | 0.88 | 16.19 | 16.12 | 1.00 | 45 | 1.33 | 1.12 | 0.84 | 11.99 | 11.87 | 0.99 | 35 | 1.33 | 1.12 | 0.84 | 11.99 | 11.87 | 0.99 |
| | 2 | 1 | 51 | 1.34 | 1.07 | 0.80 | 12.28 | 11.60 | 0.94 | 77 | 1.01 | 0.94 | 0.93 | 10.17 | 10.23 | 1.01 | 127 | 0.89 | 0.98 | 1.10 | 9.28 | 10.55 | 1.14 |
| | | 2 | 75 | 1.33 | 1.07 | 0.80 | 12.12 | 11.53 | 0.95 | 101 | 0.99 | 0.93 | 0.94 | 9.80 | 9.64 | 0.98 | 199 | 0.85 | 0.92 | 1.08 | 8.76 | 9.57 | 1.09 |
| | 3 | 1 | 103 | 1.15 | 1.01 | 0.88 | 11.22 | 11.67 | 1.04 | 129 | 0.88 | 0.99 | 1.12 | 9.27 | 10.45 | 1.13 | 279 | 0.79 | 1.10 | 1.39 | 8.83 | 11.28 | 1.28 |
| | | 2 | 157 | 1.11 | 1.00 | 0.90 | 10.14 | 11.03 | 1.09 | 183 | 0.81 | 1.02 | 1.26 | 8.76 | 10.37 | 1.18 | 441 | 0.73 | 1.01 | 1.38 | 8.52 | 10.63 | 1.25 |
| | | 3 | 211 | 1.07 | 0.99 | 0.93 | 10.28 | 11.09 | 1.08 | 237 | 0.80 | 0.92 | 1.15 | 8.84 | 10.01 | 1.13 | 603 | 0.65 | 0.86 | 1.32 | 8.11 | 10.00 | 1.23 |
| | 4 | 1 | 175 | 0.91 | 0.95 | 1.04 | 9.41 | 10.81 | 1.15 | 201 | 0.80 | 0.89 | 1.11 | 9.20 | 10.65 | 1.16 | 491 | 0.69 | 0.94 | 1.36 | 8.38 | 10.64 | 1.27 |
| | | 2 | 271 | 1.02 | 1.03 | 1.01 | 10.50 | 11.12 | 1.06 | 297 | 0.74 | 0.97 | 1.31 | 8.84 | 11.48 | 1.30 | 779 | 0.55 | 0.97 | 1.76 | 7.72 | 10.82 | 1.40 |
| | | 3 | 367 | 1.04 | 1.18 | 1.13 | 9.96 | 12.20 | 1.22 | 393 | 0.79 | 0.99 | 1.25 | 8.83 | 10.11 | 1.14 | 1067 | 0.51 | 1.01 | 1.98 | 7.42 | 11.66 | 1.57 |
| | | 4 | 463 | 0.99 | 1.16 | 1.17 | 10.02 | 11.96 | 1.19 | 489 | 0.78 | 1.00 | 1.28 | 8.75 | 11.18 | 1.28 | 1355 | 0.52 | 0.90 | 1.73 | 7.54 | 10.68 | 1.42 |
| 3 | 1 | 1 | 21 | 2.18 | 1.90 | 0.87 | 15.78 | 15.58 | 0.99 | 56 | 1.19 | 1.05 | 0.88 | 10.24 | 11.09 | 1.08 | 46 | 1.19 | 1.05 | 0.88 | 10.24 | 11.10 | 1.08 |
| | 2 | 1 | 53 | 1.32 | 1.04 | 0.79 | 12.19 | 11.45 | 0.94 | 88 | 0.95 | 0.91 | 0.96 | 9.78 | 10.42 | 1.07 | 168 | 0.89 | 0.92 | 1.03 | 9.42 | 10.23 | 1.09 |
| | | 2 | 77 | 1.31 | 1.05 | 0.80 | 12.04 | 11.42 | 0.95 | 112 | 0.92 | 0.91 | 0.99 | 9.46 | 9.57 | 1.01 | 264 | 0.82 | 0.89 | 1.09 | 9.09 | 9.86 | 1.08 |
| | 3 | 1 | 105 | 1.14 | 1.01 | 0.89 | 11.42 | 11.88 | 1.04 | 140 | 0.83 | 1.01 | 1.22 | 9.10 | 10.59 | 1.16 | 370 | 0.77 | 1.14 | 1.48 | 9.22 | 12.01 | 1.30 |
| | | 2 | 159 | 1.07 | 1.07 | 1.00 | 9.97 | 10.86 | 1.09 | 194 | 0.74 | 0.97 | 1.31 | 8.48 | 10.51 | 1.24 | 586 | 0.60 | 0.95 | 1.58 | 7.81 | 10.67 | 1.37 |
| | | 3 | 213 | 1.07 | 1.00 | 0.93 | 10.86 | 11.25 | 1.04 | 248 | 0.71 | 0.89 | 1.25 | 8.35 | 10.75 | 1.29 | 802 | 0.68 | 0.93 | 1.37 | 8.09 | 10.21 | 1.26 |
| | 4 | 1 | 177 | 0.90 | 0.94 | 1.04 | 9.39 | 10.90 | 1.16 | 212 | 0.71 | 1.10 | 1.55 | 7.97 | 10.59 | 1.33 | 652 | 0.62 | 1.15 | 1.85 | 8.50 | 10.93 | 1.29 |
| | | 2 | 273 | 0.95 | 1.08 | 1.14 | 10.32 | 11.15 | 1.08 | 308 | 0.75 | 0.91 | 1.21 | 8.36 | 9.67 | 1.16 | 1036 | 0.62 | 0.86 | 1.39 | 8.07 | 10.01 | 1.24 |
| | | 3 | 369 | 0.96 | 0.95 | 0.99 | 9.33 | 10.33 | 1.11 | 404 | 0.71 | 0.86 | 1.21 | 8.04 | 9.80 | 1.22 | 1420 | 0.53 | 1.02 | 1.92 | 7.83 | 10.33 | 1.32 |
| | | 4 | 465 | 0.89 | 0.92 | 1.03 | 9.50 | 10.89 | 1.15 | 500 | 0.65 | 0.99 | 1.52 | 7.94 | 10.18 | 1.28 | 1804 | 0.55 | 1.02 | 1.85 | 7.92 | 10.92 | 1.38 |

**Table S8:** RNA transfer matrix and tensor model parameter regression: root mean square errors (RMSE) for reaction free energy, $\Delta G_{\mathrm{rxn},37}$, and reaction enthalpy, $\Delta H_{\mathrm{rxn}}$, for training set (716 experimental melts) and test set (179 experimental melts) for a variety of model sizes $(\nu, \mu, \xi)$ using fixed-weight formulation, floating-weight formulation, or floating-block formulation. $\lvert\vec{g}\rvert$ denotes the number of parameters for each model (Section S2.7). Ratio denotes the ratio of test to training errors. Salt conditions: 1.0 M $Na^{+}$. Scatter plots for models in the highlighted row are displayed in Figure S11.

| Parameters | Ensemble | $\lvert\vec{g}\rvert$ | $\Delta G_{\text{rxn},37}$ (kcal/mol) train | test | ratio | $\Delta H_{\text{rxn}}$ (kcal/mol) train | test | ratio |
|---|---|---|---|---|---|---|---|---|
| rna06 | nostacking | ≈553 | 1.77 | 1.58 | 0.89 | 16.65 | 16.39 | 0.98 |
| rna06 | stacking | ≈553 | 1.44 | 1.32 | 0.91 | 17.77 | 17.70 | 1.00 |
| rna95 | nostacking | ≈553 | 2.16 | 2.00 | 0.92 | 17.40 | 17.11 | 0.98 |
| rna95 | stacking | ≈553 | 1.50 | 1.36 | 0.91 | 19.57 | 19.16 | 0.98 |
| dna04.3 | nostacking | ≈297 | 3.58 | 3.53 | 0.99 | 27.67 | 26.62 | 0.96 |
| dna04.3 | stacking | ≈297 | 2.52 | 2.54 | 1.00 | 28.30 | 27.51 | 0.97 |
| dna04.1 | nostacking | ≈297 | 3.62 | 3.39 | 0.94 | 28.81 | 27.01 | 0.94 |
| dna04.1 | stacking | ≈297 | 2.44 | 2.23 | 0.91 | 29.37 | 28.29 | 0.96 |

**Table S9:** RNA nearest neighbor models: root mean square errors (RMSE) for reaction free energy, $\Delta G_{\text{rxn},37}$, and reaction enthalpy, $\Delta H_{\text{rxn}}$, for training set (716 experimental melts) and test set (179 experimental melts) for an RNA reference model (`rna06`), an RNA historical model (`rna95`) and two models for a different material (`dna04.3` and `dna04.1`). $\lvert\vec{g}\rvert$ denotes the number of parameters for each model. For RNA models, there are 12,596 parameters of which ≈ 553 have unique values.[18,39–41,67,73,81,84] Ratio denotes the ratio of test to training errors (note that no training was done for these models in the present work and this partitioning into training and test sets is solely for comparison to Table S8. Salt conditions: 1.0 M $Na^+$. Scatter plots for the model in the highlighted row are displayed in Figure S11.

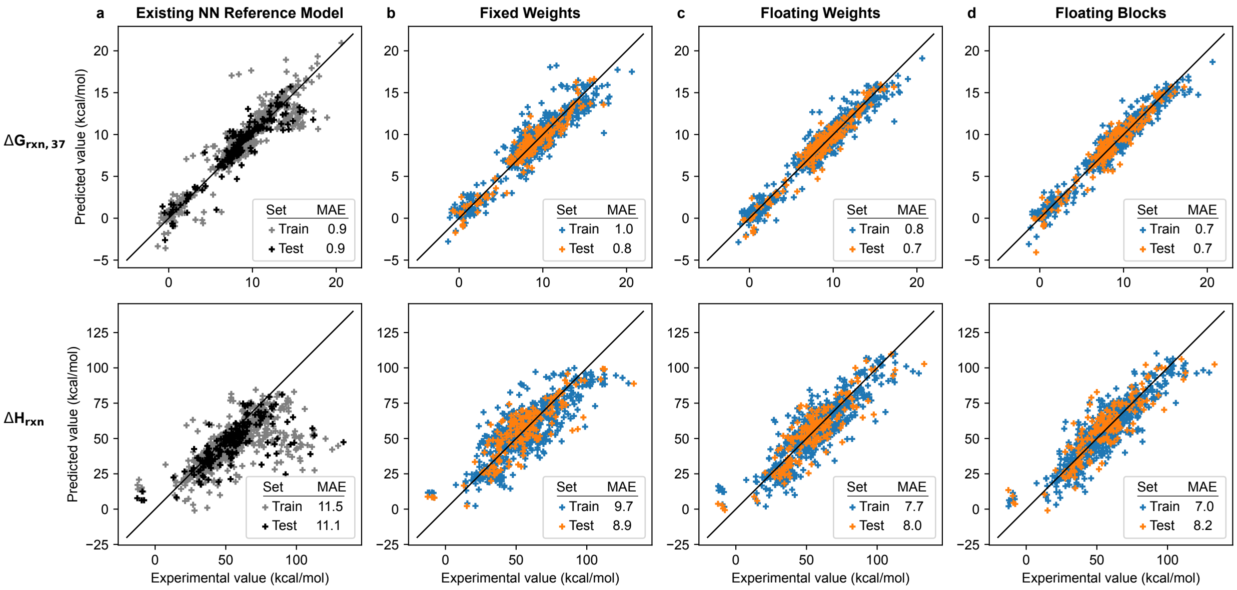

**Figure S11:** Predicted vs experimental RNA reaction free energies ($\Delta G_{\text{rxn},37}$; top) and enthalpies ($\Delta H_{\text{rxn}}$; bottom). a) Existing loop-based free energy model (panel a; NUPACK 4.1.0 with `rna06` parameters and `stacking` subensemble). b,c,d) New transfer matrix models $(\nu, \mu, \xi) = (2, 2, 1)$ using either the fixed-weight formulation (panel b), floating-weight formulation (panel c), or floating block formulation (panel d). Mean absolute error (MAE) in kcal/mol. Training set: 716 melts. Test set: 179 melts. Salt conditions: 1.0 M $Na^+$.

| | | | Fixed Weights | | | | | | | Floating Weights | | | | | | | Floating Blocks | | | | | | |
|---|---|---|---|---|---|---|---|---|---|---|---|---|---|---|---|---|---|---|---|---|---|---|
| | | | | $\Delta G_{\mathrm{rxn},37}$ (kcal/mol) | | | $\Delta H_{\mathrm{rxn}}$ (kcal/mol) | | | | $\Delta G_{\mathrm{rxn},37}$ (kcal/mol) | | | $\Delta H_{\mathrm{rxn}}$ (kcal/mol) | | | | $\Delta G_{\mathrm{rxn},37}$ (kcal/mol) | | | $\Delta H_{\mathrm{rxn}}$ (kcal/mol) | | |
| $\nu$ | $\mu$ | $\xi$ | $\lvert\vec{g}\rvert$ | train | test | ratio | train | test | ratio | $\lvert\vec{g}\rvert$ | train | test | ratio | train | test | ratio | $\lvert\vec{g}\rvert$ | train | test | ratio | train | test | ratio |
| 1 | 1 | 1 | 15 | 3.27 | 3.29 | 1.01 | 18.32 | 20.11 | 1.10 | 28 | 1.04 | 1.08 | 1.04 | 9.48 | 10.11 | 1.07 | 20 | 1.04 | 1.08 | 1.04 | 9.48 | 10.11 | 1.07 |
| | 2 | 1 | 41 | 0.83 | 0.93 | 1.12 | 10.45 | 13.13 | 1.26 | 54 | 0.74 | 0.87 | 1.18 | 8.94 | 10.35 | 1.16 | 70 | 0.69 | 0.79 | 1.14 | 8.62 | 11.54 | 1.34 |
| | | 2 | 57 | 0.78 | 0.92 | 1.18 | 9.86 | 11.75 | 1.19 | 70 | 0.64 | 0.83 | 1.30 | 9.58 | 12.60 | 1.32 | 102 | 0.58 | 0.81 | 1.40 | 9.10 | 13.12 | 1.44 |
| | 3 | 1 | 83 | 0.67 | 0.91 | 1.36 | 8.92 | 12.27 | 1.38 | 96 | 0.56 | 0.76 | 1.36 | 7.80 | 11.14 | 1.43 | 152 | 0.45 | 0.88 | 1.96 | 6.42 | 10.36 | 1.61 |
| | | 2 | 119 | 0.61 | 0.85 | 1.39 | 8.51 | 11.07 | 1.30 | 132 | 0.55 | 0.85 | 1.55 | 7.83 | 11.36 | 1.45 | 224 | 0.42 | 0.75 | 1.79 | 6.65 | 10.53 | 1.58 |
| | | 3 | 155 | 0.62 | 0.97 | 1.56 | 8.73 | 12.63 | 1.45 | 168 | 0.52 | 0.78 | 1.50 | 8.11 | 12.36 | 1.52 | 296 | 0.44 | 0.78 | 1.77 | 6.44 | 11.63 | 1.81 |
| | 4 | 1 | 141 | 0.49 | 0.78 | 1.59 | 7.77 | 11.83 | 1.52 | 154 | 0.49 | 0.98 | 2.00 | 7.75 | 14.97 | 1.93 | 266 | 0.34 | 0.92 | 2.71 | 6.05 | 11.81 | 1.95 |
| | | 2 | 205 | 0.47 | 0.76 | 1.62 | 7.16 | 11.74 | 1.64 | 218 | 0.46 | 0.64 | 1.39 | 7.32 | 10.44 | 1.43 | 394 | 0.36 | 0.98 | 2.72 | 5.95 | 14.39 | 2.42 |
| | | 3 | 269 | 0.45 | 0.93 | 2.07 | 7.09 | 11.02 | 1.55 | 282 | 0.46 | 0.72 | 1.57 | 7.22 | 11.97 | 1.66 | 522 | 0.35 | 0.89 | 2.54 | 6.12 | 12.95 | 2.12 |
| | | 4 | 333 | 0.49 | 0.92 | 1.88 | 7.63 | 13.84 | 1.81 | 346 | 0.42 | 0.86 | 2.05 | 6.94 | 12.67 | 1.83 | 650 | 0.32 | 0.86 | 2.69 | 6.12 | 13.96 | 2.28 |
| 2 | 1 | 1 | 17 | 2.74 | 2.84 | 1.04 | 17.78 | 20.38 | 1.15 | 37 | 0.87 | 0.87 | 1.00 | 8.82 | 9.38 | 1.06 | 29 | 0.87 | 0.87 | 1.00 | 8.82 | 9.37 | 1.06 |
| | 2 | 1 | 43 | 0.84 | 0.95 | 1.13 | 10.39 | 13.11 | 1.26 | 63 | 0.68 | 0.86 | 1.26 | 8.47 | 10.44 | 1.23 | 103 | 0.55 | 0.80 | 1.45 | 7.44 | 11.05 | 1.49 |
| | | 2 | 59 | 0.81 | 0.90 | 1.11 | 9.84 | 11.96 | 1.22 | 79 | 0.64 | 0.78 | 1.22 | 8.17 | 11.81 | 1.45 | 151 | 0.48 | 0.79 | 1.65 | 7.28 | 10.81 | 1.48 |
| | 3 | 1 | 85 | 0.65 | 1.01 | 1.55 | 8.87 | 14.48 | 1.63 | 105 | 0.56 | 0.78 | 1.39 | 8.02 | 11.43 | 1.43 | 225 | 0.42 | 0.91 | 2.17 | 6.44 | 11.92 | 1.85 |
| | | 2 | 121 | 0.60 | 0.76 | 1.27 | 8.02 | 11.80 | 1.47 | 141 | 0.48 | 0.72 | 1.50 | 7.08 | 11.87 | 1.68 | 333 | 0.34 | 0.97 | 2.85 | 6.42 | 13.28 | 2.07 |
| | | 3 | 157 | 0.59 | 0.95 | 1.61 | 8.85 | 12.06 | 1.36 | 177 | 0.45 | 0.75 | 1.67 | 6.85 | 11.20 | 1.64 | 441 | 0.30 | 1.42 | 4.73 | 5.48 | 14.39 | 2.63 |
| | 4 | 1 | 143 | 0.51 | 0.71 | 1.39 | 7.85 | 11.60 | 1.48 | 163 | 0.42 | 0.93 | 2.21 | 6.72 | 11.15 | 1.66 | 395 | 0.31 | 0.99 | 3.19 | 6.01 | 11.82 | 1.97 |
| | | 2 | 207 | 0.47 | 0.85 | 1.81 | 7.66 | 13.33 | 1.74 | 227 | 0.38 | 0.80 | 2.11 | 6.51 | 12.75 | 1.96 | 587 | 0.26 | 0.96 | 3.69 | 4.80 | 13.14 | 2.74 |
| | | 3 | 271 | 0.46 | 0.73 | 1.59 | 7.58 | 12.56 | 1.66 | 291 | 0.39 | 1.05 | 2.69 | 6.38 | 14.33 | 2.25 | 779 | 0.29 | 0.98 | 3.38 | 5.77 | 12.89 | 2.23 |
| | | 4 | 335 | 0.60 | 0.80 | 1.33 | 8.61 | 13.66 | 1.59 | 355 | 0.38 | 0.82 | 2.16 | 6.50 | 13.51 | 2.08 | 971 | 0.21 | 1.23 | 5.86 | 4.61 | 12.95 | 2.81 |
| 3 | 1 | 1 | 19 | 2.46 | 2.57 | 1.04 | 17.06 | 19.93 | 1.17 | 46 | 0.77 | 0.86 | 1.12 | 8.66 | 9.68 | 1.12 | 38 | 0.77 | 0.86 | 1.12 | 8.66 | 9.69 | 1.12 |
| | 2 | 1 | 45 | 0.85 | 0.97 | 1.14 | 10.39 | 13.08 | 1.26 | 72 | 0.63 | 0.80 | 1.27 | 8.28 | 10.28 | 1.24 | 136 | 0.45 | 0.75 | 1.67 | 7.07 | 10.78 | 1.52 |
| | | 2 | 61 | 0.85 | 0.97 | 1.14 | 10.38 | 13.07 | 1.26 | 88 | 0.57 | 0.82 | 1.44 | 8.08 | 11.47 | 1.42 | 200 | 0.40 | 0.84 | 2.10 | 6.43 | 12.45 | 1.94 |
| | 3 | 1 | 87 | 0.65 | 1.03 | 1.58 | 9.06 | 13.62 | 1.50 | 114 | 0.48 | 0.69 | 1.44 | 6.90 | 9.37 | 1.36 | 298 | 0.33 | 0.92 | 2.79 | 5.87 | 12.70 | 2.16 |
| | | 2 | 123 | 0.61 | 0.72 | 1.18 | 8.28 | 11.46 | 1.38 | 150 | 0.45 | 0.76 | 1.69 | 6.85 | 10.91 | 1.59 | 442 | 0.31 | 0.97 | 3.13 | 5.39 | 13.60 | 2.52 |
| | | 3 | 159 | 0.58 | 0.88 | 1.52 | 8.19 | 13.44 | 1.64 | 186 | 0.43 | 0.79 | 1.84 | 6.79 | 11.13 | 1.64 | 586 | 0.27 | 0.91 | 3.37 | 5.26 | 13.81 | 2.63 |
| | 4 | 1 | 145 | 0.51 | 0.79 | 1.55 | 7.52 | 12.21 | 1.62 | 172 | 0.43 | 0.77 | 1.79 | 6.82 | 10.54 | 1.55 | 524 | 0.29 | 1.27 | 4.38 | 5.32 | 12.07 | 2.27 |
| | | 2 | 209 | 0.47 | 0.81 | 1.72 | 7.49 | 15.61 | 2.08 | 236 | 0.41 | 0.80 | 1.95 | 6.43 | 11.14 | 1.73 | 780 | 0.21 | 1.34 | 6.38 | 4.85 | 16.64 | 3.43 |
| | | 3 | 273 | 0.48 | 0.80 | 1.67 | 7.15 | 10.41 | 1.46 | 300 | 0.36 | 1.01 | 2.81 | 6.04 | 11.98 | 1.98 | 1036 | 0.23 | 1.33 | 5.78 | 5.16 | 14.68 | 2.84 |
| | | 4 | 337 | 0.53 | 0.92 | 1.74 | 7.80 | 13.38 | 1.72 | 364 | 0.33 | 0.86 | 2.61 | 6.73 | 12.63 | 1.88 | 1292 | 0.22 | 1.04 | 4.73 | 4.65 | 14.90 | 3.20 |

**Table S10:** DNA transfer matrix and tensor model parameter regression: root mean square errors (RMSE) for reaction free energy, $\Delta G_{\mathrm{rxn},37}$, and reaction enthalpy, $\Delta H_{\mathrm{rxn}}$, for training set (371 experimental melts) and test set (93 experimental melts) for a variety of model sizes $(\nu, \mu, \xi)$ using fixed-weight formulation, floating-weight formulation, or floating-block formulation. $\lvert\vec{g}\rvert$ denotes the number of parameters for each model (Section S2.7). Ratio denotes the ratio of test to training errors. Salt conditions: 1.0 M $Na^{+}$. Scatter plots for models in the highlighted row are displayed in Figure S12.

| | | | $\Delta G_{\mathrm{rxn},37}$ (kcal/mol) | | | $\Delta H_{\mathrm{rxn}}$ (kcal/mol) | | |
|---|---|---|---|---|---|---|---|---|
| Parameters | Ensemble | $\lvert\vec{g}\rvert$ | train | test | ratio | train | test | ratio |
| `dna04.3` | `nostacking` | ≈297 | 1.22 | 1.28 | 1.05 | 13.21 | 16.15 | 1.22 |
| `dna04.3` | `stacking` | ≈297 | 1.30 | 1.36 | 1.05 | 19.38 | 21.63 | 1.12 |
| `dna04.1` | `nostacking` | ≈297 | 1.64 | 1.72 | 1.05 | 15.63 | 20.16 | 1.29 |
| `dna04.1` | `stacking` | ≈297 | 1.59 | 1.70 | 1.06 | 23.88 | 27.13 | 1.14 |
| `rna06` | `nostacking` | ≈553 | 4.78 | 4.44 | 0.93 | 19.75 | 19.96 | 1.01 |
| `rna06` | `stacking` | ≈553 | 4.81 | 4.46 | 0.93 | 15.63 | 17.82 | 1.14 |
| `rna95` | `nostacking` | ≈553 | 2.68 | 2.51 | 0.94 | 15.66 | 18.37 | 1.17 |
| `rna95` | `stacking` | ≈553 | 2.48 | 2.29 | 0.93 | 18.83 | 22.26 | 1.18 |

**Table S11:** DNA nearest neighbor models: root mean square errors (RMSE) for reaction free energy, $\Delta G_{\mathrm{rxn},37}$, and reaction enthalpy, $\Delta H_{\mathrm{rxn}}$, for training set (371 experimental melts) and test set (93 experimental melts) for a DNA reference model (`dna04.3`), a DNA historical model (`dna04.1`) and two models for a different material (`rna06` and `rna95`). $\lvert\vec{g}\rvert$ denotes the number of parameters for each model. For DNA models, there are 12,756 parameters of which ≈ 297 have unique values.[5, 18, 52, 58, 59, 84] Ratio denotes the ratio of test to training errors (note that no training was done for these models in the present work and this partitioning into training and test sets is solely for comparison to Table S10). Salt conditions: 1.0 M $Na^+$. Scatter plots for the model in the highlighted row are displayed in Figure S12.

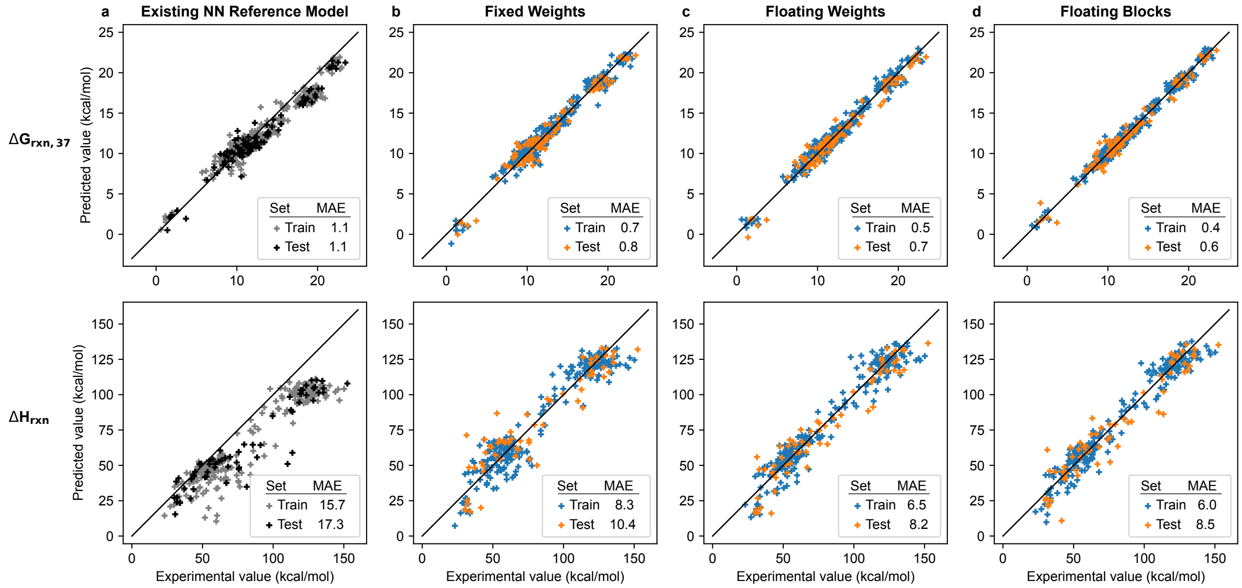

**Figure S12:** Predicted vs experimental DNA reaction free energies ($\Delta G_{\mathrm{rxn},37}$; top) and enthalpies ($\Delta H_{\mathrm{rxn}}$; bottom) (cf. Figure 9). a) Existing loop-based free energy model (panel a; NUPACK 4.1.0 with `dna04.3` parameters and `stacking` subensemble). b,c,d) New transfer matrix models $(\nu, \mu, \xi) = (2, 2, 1)$ using either the fixed-weight formulation (panel b), floating-weight formulation (panel c), or floating block formulation (panel d). Mean absolute error (MAE) in kcal/mol. Training set: 371 melts. Test set: 93 melts. Salt conditions: 1.0 M $Na^+$.

#### S10 Additional studies

##### S10.1 Empirical dependence of ensemble size on complex size

We performed calculations to measure the number of secondary structures, $|\overline{\Gamma}(\phi)|$, and subensemble states, $|\overline{\Gamma}^{\shortparallel}(\phi)|$, for a set of RNA complexes with random sequences. Empirically, $|\overline{\Gamma}(\phi)|$ and $|\overline{\Gamma}^{\shortparallel}(\phi)|$ grow exponentially with the number of nucleotides in the complex (Figure S13). Least-squares linear regressions on the log-linear data yield the fits:

$$|\overline{\Gamma}(\phi)| = 0.00834 \cdot 1.832^N, \qquad (r = 0.9999997), \tag{S121}$$

$$|\overline{\Gamma}^{\shortparallel}(\phi)| = 0.00388 \cdot 2.106^N, \qquad (r = 0.9999999) \text{ for } \nu, \mu, \xi = (2, 1, 1), \tag{S122}$$

$$|\overline{\Gamma}^{\shortparallel}(\phi)| = 0.0687 \cdot 3.664^N, \qquad (r = 0.9999999) \text{ for } \nu, \mu, \xi = (1, 2, 1), \tag{S123}$$

$$|\overline{\Gamma}^{\shortparallel}(\phi)| = 0.0287 \cdot 4.212^N, \qquad (r = 0.9999999) \text{ for } \nu, \mu, \xi = (2, 2, 1), \tag{S124}$$

$$|\overline{\Gamma}^{\shortparallel}(\phi)| = 0.0655 \cdot 4.942^N, \qquad (r = 0.9999999) \text{ for } \nu, \mu, \xi = (2, 2, 2). \tag{S125}$$

Note that these results are sequence-dependent (e.g., $\phi$ = AAAAA... will have an ensemble size of $|\overline{\Gamma}(\phi)| = 1$ independent of complex size). Unlike existing NN free energy models, which approximate single-stranded secondary structures as having a zero free energy independent of sequence, the new transfer matrix and tensor models calculate the secondary structure free energy over a subensemble of states for all unpaired bases so $|\overline{\Gamma}^{\shortparallel}(\phi)| > 1$ even for a sequence with only one secondary structure.

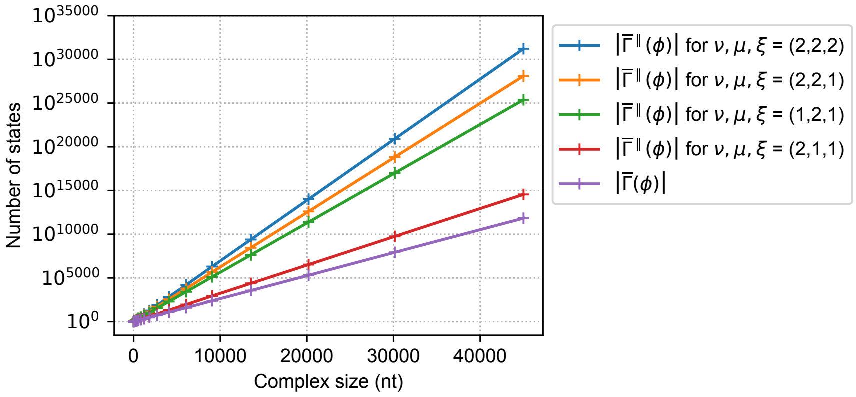

**Figure S13:** Empirical dependence of ensemble size on complex size. Each RNA complex comprises 3 random sequences of equal length. The number of secondary structures $|\overline{\Gamma}(\phi)|$ is independent of the transfer matrix or tensor model size $(\nu, \mu, \xi)$. The number of subensemble states, $|\overline{\Gamma}^{\shortparallel}(\phi)|$, exceeds the number of secondary structures, $|\overline{\Gamma}(\phi)|$, and increases with model size. Transfer matrix or tensor models using the fixed-weight formulation for model sizes $\nu, \mu, \xi$ noted in the legend.

#### S10.2 Empirical dependence of partition function cost on model dimensions

Table S12 displays the cost of partition function calculations for a range of transfer matrix and tensor model sizes $(\nu, \mu, \xi)$ across a range of complex sizes up to 30,000 nt. The dependence of algorithm cost on matrix size is mild, with all tested models being within a factor of 3 in cost to the representative $\nu, \mu, \xi = (2, 2, 1)$ model, indicating that the choice of model can be made based on accuracy rather than cost.

| Model | | | Complex size (nt) | | | | | | | | | | | | | |
|---|---|---|---|---|---|---|---|---|---|---|---|---|---|---|---|---|
| $\nu$ | $\mu$ | $\xi$ | 18 | 30 | 57 | 99 | 174 | 309 | 549 | 969 | 1719 | 3045 | 5394 | 9558 | 16935 | 30000 |
| 1 | 1 | 1 | 0.43 | 0.61 | 0.84 | 0.63 | 0.61 | 0.49 | 0.52 | 0.77 | 0.60 | 0.56 | 0.45 | 0.37 | 0.33 | 0.23 |
| 2 | 1 | 1 | 0.46 | 0.90 | 0.97 | 0.70 | 0.71 | 0.53 | 0.56 | 0.83 | 0.70 | 0.62 | 0.50 | 0.40 | 0.36 | 0.24 |
| | 2 | 1 | 1.00 | 1.00 | 1.00 | 1.00 | 1.00 | 1.00 | 1.00 | 1.00 | 1.00 | 1.00 | 1.00 | 1.00 | 1.00 | 1.00 |
| | | 2 | 0.64 | 0.98 | 1.05 | 1.10 | 1.12 | 0.95 | 0.79 | 1.02 | 1.00 | 1.00 | 1.00 | 0.99 | 1.00 | 1.00 |
| 3 | 1 | 1 | 0.64 | 0.83 | 1.03 | 0.82 | 0.80 | 0.63 | 0.92 | 1.35 | 1.03 | 0.80 | 0.58 | 0.50 | 0.43 | 0.31 |
| | 2 | 1 | 0.78 | 1.13 | 1.26 | 1.09 | 1.21 | 1.20 | 1.09 | 1.18 | 1.14 | 1.19 | 1.22 | 1.23 | 1.29 | 1.32 |
| | | 2 | 0.79 | 1.46 | 1.25 | 1.26 | 1.32 | 1.13 | 0.83 | 1.27 | 1.21 | 1.18 | 1.22 | 1.27 | 1.29 | 1.31 |
| | 3 | 1 | 0.97 | 1.58 | 1.86 | 2.05 | 2.54 | 1.84 | 1.26 | 1.54 | 1.59 | 1.88 | 2.09 | 2.37 | 2.74 | |
| | | 2 | 1.14 | 1.40 | 1.95 | 2.24 | 2.62 | 2.08 | 1.21 | 1.67 | 1.65 | 1.83 | 2.08 | 2.38 | 2.74 | |
| | | 3 | 1.40 | 2.13 | 2.17 | 2.06 | 2.17 | 1.36 | 0.93 | 1.56 | 1.62 | 1.85 | 2.10 | 2.38 | 2.74 | |

**Table S12:** Cost of calculating the partition function using different transfer matrix and tensor model dimensions $(\nu, \mu, \xi)$ normalized by the cost of the representative model $\nu, \mu, \xi = (2, 2, 1)$ (highlighted row). Production implementation (CPU parallel for small complexes and GPU for large complexes) for all algorithms. Timings omitted for 30,000 nt complexes using the largest model dimensions due to insufficient GPU memory. Fixed-weight formulation for all models. Each RNA complex comprises 3 random sequences of equal length. Mean wall clock time over 3 sets of random sequences per complex size. Conditions: RNA, 37° C, 1.0 M $Na^{+}$.

#### S10.3 Relative cost of partition function, equilibrium pair probability, and MFE calculations

The cost of calculating the partition function, $\overline{Q}(\phi)$, equilibrium pair probability matrix, $\overline{P}(\phi)$, and minimum free energy (MFE) of a subensemble state, $\overline{\Delta G}(\phi, s^{\shortparallel}_{\text{MFE}})$, are profiled in Figure S14.

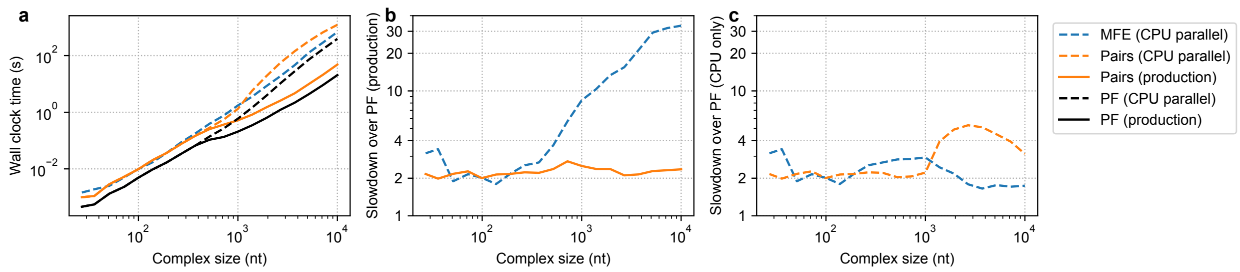

**Figure S14:** Relative cost of partition function, equilibrium pair probability, and MFE calculations. (a) Computational cost. (b) Relative cost of calculating the MFE (CPU parallel implementation) or pair probability matrix (production implementation) relative to the partition function (production implementation). (c) Relative cost of calculating the MFE (CPU parallel implementation) or pair probability matrix (CPU parallel implemenation) relative to the partition function (CPU parallel implementation). Each RNA complex comprises 3 random sequences of equal length. Transfer matrix model with $\nu, \mu, \xi = (2, 2, 1)$ using the fixed-weight formulation. Conditions: RNA, 37° C, 1.0 M $Na^+$.

### S11 Algorithm validation

#### S11.1 Tests

Over 20 types of C++ tests were run to validate the new transfer matrix and tensor models. Some unit tests compare the output from a dynamic program to output from an exhaustive enumeration algorithm. This type of comparison can be done only for small complexes because the time complexity of exhaustive enumeration is exponential in the number of nucleotides.

- **Comparisons between dynamic programming and exhaustive enumeration for the following physical quantities:**
  - **Partition function:** exhaustive enumeration (Algorithm S15 below) is used to calculate $\overline{Q}(\phi)$ using the sum of (S11) and post-processed to calculate $Q(\phi)$ (see Section S5 of Reference 19). Compare to dynamic programming post-processed output $Q(\phi)$ to the relevant floating point precision.
  - **Equilibrium base-pairing probability matrix:** exhaustive enumeration is used to calculate $\overline{P}(\phi)$ using the sum of (S13). Compare to dynamic programming output to the relevant floating point precision.
  - **MFE:** exhaustive enumeration is used to calculate the free energy of the MFE subensemble state, $\overline{\Delta G}(\phi, s_{\text{MFE}'})$, using the minimization of (S14). Compare to dynamic programming output to the relevant floating point precision.
  - **MFE proxy structure:** exhaustive enumeration is used to calculate the MFE proxy structure(s) $s_{\text{MFE}'}$ using equation (S15). The dynamic program for MFE proxy structure determination is the suboptimal secondary structure algorithm with $\Delta G_{\text{gap}} = 0$ kcal/mol. Compare to dynamic programming output.
  - **Suboptimal structures:** exhaustive enumeration is used to calculate the set of suboptimal secondary structures, $\overline{\Gamma}_{\text{subopt}}(\phi, \Delta G_{\text{gap}})$ within a specified $\Delta G_{\text{gap}} \geq 0$ of the MFE subsensemble state using equation (S18). Tests are performed using $\Delta G_{\text{gap}} = 0.4$ kcal/mol. Compare to dynamic programming output.
- **Comparison of equilibrium base-pairing probability and Boltzmann sampled structure dynamic programs.** Estimate equilibrium the base-pairing probability matrix from Boltzmann-sampled structures and check convergence to the exact base-pairing probability matrix as the number of sampled structures increases.
- **Comparison of MFE and free energy of MFE proxy structure.** Calculate the free energy of the MFE stacking state, $\overline{\Delta G}(\phi, s_{\text{MFE}'})$ using the forward dynamic program. Calculate the MFE proxy structure, $s_{\text{MFE}'}$, using the backtracking dynamic program for suboptimal structure generation with $\Delta G_{\text{gap}} = 0$ kcal/mol. Calculate the free energy of the MFE proxy structure using equation (1) and compare it to the value calculated using the forward algorithm to the relevant floating point precision.
- **Comparison to simplified reference implementations.** Block-based implementations for calculating the secondary structure free energy, $\overline{g}(\phi, s)$, and all of the above physical quantities were compared to results calculated using fully dense $\gamma \times \gamma$ transfer matrices or tensors. These block-based implementations were also compared to reference implementations using no vectorization and no BLAS/CuBLAS invocations.
- **Comparison of GPU-accelerated dynamic programs to CPU implementations.** GPU-accelerated dynamic programs for calculating the partition function and the equilibrium base pairing probability matrix are compared to CPU implementations to the relevant floating point precision.
- **Comparison of analytic gradients to finite differences.** Analytic gradients (e.g., of $\vec{g}_{37}$) are compared to finite difference gradients to the relevant floating point precision.

#### S11.2 Exhaustive enumeration algorithm

For a complex with sequence $\phi$, Algorithm S15 enumerates all unpseudoknotted secondary structures $s \in \overline{\Gamma}(\phi)$. The algorithm implicitly relies on a lexicographic ordering of base pairs by their indices $(i, j)$.

**Algorithm S15** Exhaustive enumeration of all secondary structures $s \in \overline{\Gamma}(\phi)$ for sequence $\phi$

```
procedure RecurseStructure(b, v, s, φ)
    v ← v ∪ {s}                                                      ▷ Add structure to set
    for j ∈ [0:|φ|), i ∈ [0:j) do                                     ▷ For each base pair
        if (i, j) < b and i unpaired in s and j unpaired in s then    ▷ Omit base pairs ordered above bound b
            s′ ← s with base pair i · j
            if s′ is a valid unpseudoknotted secondary structure for φ then   ▷ Omit pseudoknots, impossible pairs
                RecurseStructure((i, j), v, s′, φ)

function EnumerateStructures(φ)
    v ← ∅
    s ← unpaired secondary structure of size |φ|
    RecurseStructure((|φ|, |φ|), v, s, φ)     ▷ Initialize with a bound greater than any possible base pair
    v ← {s | s ∈ v, Connected(v)}             ▷ Remove disconnected structures
    return v
```

### S12 Data files

#### S12.1 Literature reaction free energies and enthalpies for experimental melts

Literature $\Delta G^{\text{lit}}_{\text{rxn},37}$ and $\Delta H^{\text{lit}}_{\text{rxn}}$ values for experimental melts for RNA or DNA (see Section S9.1):

- Files: `{rna,dna}-lit-melts.json`

#### S12.2 Free energy parameters from regression to existing NN reference models

Free energy parameters, $\vec{g}_{37}$, at reference temperature 37 °C for new RNA or DNA transfer matrix models of various sizes $(\nu, \mu, \xi)$ using either the fixed-weight formulation, floating-weight formulation, or floating-block formulation, regressed to existing NN reference models via KL divergence optimization (see Section S8):

- $\vec{g}_{37}$ parameters:

  `{rna,dna}-{fixed-weights,floating-weights,floating-blocks}-kl-g37-params.json`

- $\vec{g}_{37}$ matrices $(\tilde{V}, \tilde{B}, Z, S)$:

  `{rna,dna}-{fixed-weights,floating-weights,floating-blocks}-kl-g37-matrices.json`

- Text file containing detailed regression information:

  `{rna,dna}-kl-fit.csv`

#### S12.3 Free energy and enthalpy parameters from regression to experimental melts

Free energy parameters, $\vec{g}_{37}$, at reference temperature 37 °C, and enthalpy parameters, $\vec{h}$, for new RNA or DNA transfer matrix models of various sizes $(\nu, \mu, \xi)$ using either the fixed-weight formulation, floating-weight formulation, or floating-block formulation, regressed to experimental melt data (see Section S9.6):

- $\vec{g}_{37}$ parameters:

  `{rna,dna}-{fixed-weights,floating-weights,floating-blocks}-melt-g37-params.json`

- $\vec{g}_{37}$ matrices $(\tilde{V}, \tilde{B}, Z, S)$:

  `{rna,dna}-{fixed-weights,floating-weights,floating-blocks}-melt-g37-matrices.json`

- $\vec{h}$ parameters:

  `{rna,dna}-{fixed-weights,floating-weights,floating-blocks}-melt-h-params.json`

- $\vec{h}$ matrices $(\tilde{V}, \tilde{B}, Z, S)$:

  `{rna,dna}-{fixed-weights,floating-weights,floating-blocks}-melt-h-matrices.json`

- Text file containing detailed regression information:

  `{rna,dna}-melt-fit.csv`
